# Native-like 3D colonoids reveal mechanisms of colonic drug disposition

**DOI:** 10.64898/2026.11.16.766943

**Authors:** Rebekkah Hammar, Patrik Lundquist, Dinh Son Vo, Foteini Tzioufa, Jens Eriksson, Ana C.C. Lopes, Helen S. Hammer, Oliver Pötz, Maria Letizia di Martino, Wilhelm Graf, Madlen Hubert, Magnus Sundbom, Mikael E. Sellin, Per Artursson

## Abstract

Intestinal organoid research in drug absorption, distribution, metabolism, excretion, and toxicity (ADME-T) has focused predominantly on small intestinal enteroids, leaving colonic drug handling comparatively unexplored. Here, we established and characterized adult colonic stem cell-based three-dimensional (3D) organoid (colonoid) models as physiologically relevant New Approach Methodologies (NAMs). Donor-matched mucosal tissue, isolated primary epithelial cells, and 3D colonoids were characterized using global and targeted proteomics, fixed and live-cell microscopy, functional ADME-T assays and targeted lipidomics. This complementary analysis identified isolated primary epithelial cells as a more representative native epithelial reference than excised mucosal tissue. Suspended differentiated apical- and basal-out colonoid configurations more closely matched this reference across the assessed characteristics. Together, these two configurations provided access to both apical and basolateral epithelial surfaces, enabling functional studies of key drug transporters (P-gp, BCRP) and metabolizing enzymes (CYP3A4, UGT1A, SULT1A) in a native-like setting. Measurement of intracellular bioavailability (F_ic_), reflecting drug availability to intracellular targets, revealed lipophilicity-dependent microvillar absorption mechanisms. Hydrophilic compounds exploited the enlarged membrane surface provided by microvilli, whereas highly permeable lipophilic compounds seemingly entered only at the microvillus tips. Compared with HEK293 cells, an established reference model for F_ic_ measurements, colonoids combined native-like epithelial geometry with higher phospholipid content and generally stronger intracellular drug binding, highlighting the importance of both architecture and cellular composition for studying colonic drug disposition. These colonoid models provide a physiologically relevant platform for investigating colonic ADME-T and reveal how microvillar architecture shapes intracellular drug bioavailability.

**Highlights:**

- Donor-matched 3D colonoids benchmarked against native colonic epithelium
- Native-like 3D colonoids reveal mechanisms of colonic drug disposition
- Adapted 3D assays determine ADME-T, including intracellular bioavailability
- Microvilli shape drug uptake in a lipophilicity-dependent manner
- Lipidomics identifies lipid class correlates with cellular drug binding

**Graphical abstract:** 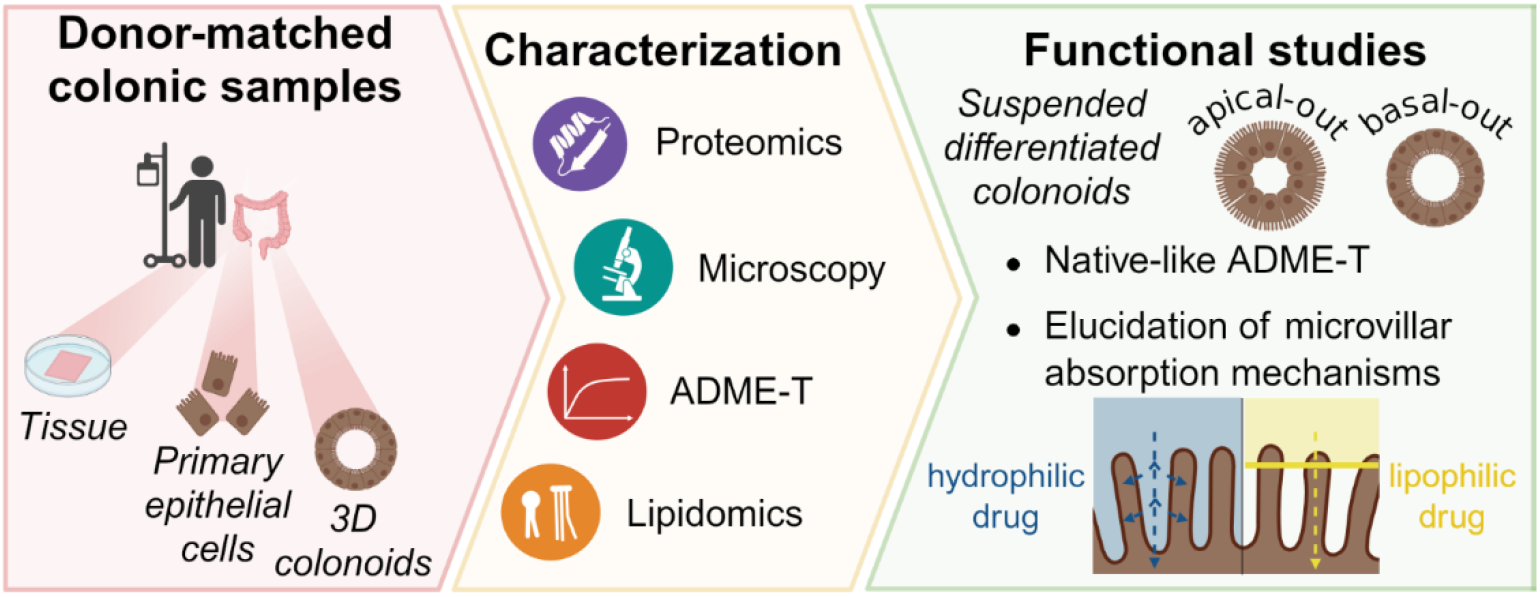

## 1 Introduction

With colonic diseases on the rise globally, the need for assessing colonic drug exposure is increasing [1]. Although the relevance of colon-mediated exposure to both local and systemic drug action has been confirmed [2–8], its underlying mechanisms remain poorly understood. Specifically, mechanisms governing colonic absorption, distribution, metabolism, excretion, and toxicity (ADME-T) have received limited attention.

This knowledge gap partly reflects established models’ failure to recapitulate key aspects of colonic physiology [1]. Organ explants, animal studies, and immortalized cell lines are limited by their culturability [9], species transferability [10], and lack of organotypic phenotypes [11,12], respectively. The rise of organoid technology [13] has, however, provided access to culturable human colonic epithelium. These models, termed colonoids when established from adult intestinal epithelial stem cells, have the potential to become New Approach Methodologies (NAMs) for ADME-T research, provided they are suitably engineered and validated. To fulfill this potential, colonoids must 1) reflect native physiology, including its underlying proteomes and lipidomes, 2) be reproducibly culturable, and 3) permit the study of absorptive and secretory processes while maintaining cellular and barrier integrity. Knowledge gaps remain in each of these areas.

1. Establishing physiological relevance requires representative native reference material against which colonoids can be benchmarked. However, efforts to standardize colonoids have been constrained not only by cell handling but also by a lack of representative reference proteomes. The clinical relevance of ADME-T transcriptomic data [14] is inherently debatable due to poor RNA transcript-protein correlation [15–17]. Moreover, colonic ADME-T characterization using either transcriptomics or proteomics [14,18,19] has relied exclusively on colonic tissue as a native reference, in the cases a reference was included. Such references are biased by submucosal cell types. Intestinal biopsies consist of approximately 50% mucosal cells and 50% submucosal cells, skewing proteomic or transcriptomic data away from actual epithelial cell expression [20,21]. Partly because of the use of such bulk tissue, ADME-T protein quantification has typically required targeted approaches, limiting the number of proteins that could be studied [19]. Together, these limitations have left the colonic ADME-T proteome incompletely mapped. Cell isolation reduces unwanted proteomic background signal [22], but enzymatic isolation can damage surface localized proteins, including transporters and enzymes [23]. Additionally, few studies report donor-matched proteomes comparing source colonic material with their derivative cultured systems, and none of these focus on ADME-T [24]. Matched healthy lipidomic references are also lacking, as most lipid profiling in this field has focused on colorectal cancer [25–27].
2. Beyond appropriate benchmarking, generating organoids reproducibly at comparable differentiation states generally remains a challenge, especially across passages [28]. Culture conditions also determine epithelial orientation and accessibility. In basement membrane matrix, colonoids are basal-out, or apical-in, 3D structures, limiting access to the apical membrane involved in the absorption of orally administered drugs [29]. Colonoid-derived monolayers provide easy access to the apical membrane [30], but do not reproduce the curved, 3D geometry of the native intestine [29] unless grown on specialized inserts [31]. Furthermore, transcriptomic [32,33] and proteomic [30] analysis indicate that these monolayers do not fully differentiate unless maintained in air-liquid interface culture.
3. Although a 3D format addresses the issues of geometry and differentiation [29], it introduces challenges for functional studies. Microinjecting 3D colonoids without compromising their integrity is technically challenging and difficult to standardize [34]. Suspension culture is therefore preferable, allowing intestinal organoids to invert their geometry and expose their apical surface [29,35,36]. However, standardized characterization assays in the field use an adherent two-dimensional (2D) format [37], and require substantial adaptation to a suspended 3D model. Thus, realizing the potential of colonoids for ADME-T research requires both suitable culture conditions and functional assays adapted to their architecture.

To address these knowledge gaps, we first developed protocols to reproducibly generate suspended 3D human colonoids of varying degrees of differentiation. We established donor-matched reference proteomes, using purified, non-enzymatically isolated colonic epithelial cells. We assessed the epithelial representativeness of the resulting colonoids through global proteomics, morphology, viability, barrier integrity, and functional ADME-T assays using methods adapted to a 3D format. The best-performing models, apical-out and basal-out differentiated (AD and BD), were then used to assess colonic drug exposure to a representative drug panel. Intracellular bioavailability (F_ic_), which describes intracellular unbound drug exposure relative to the extracellular concentration [38], was determined in AD and BD colonoids and compared with values in human embryonic kidney 293 (HEK293) cells, an established model for F_ic_ measurements in drug discovery [38–41]. Finally, lipidomic analysis complemented model-specific intracellular binding measurements to investigate how differences in lipid content and composition relate to drug binding in colonoids and HEK293 cells.

## 2 Materials and methods

Unless otherwise stated, reagents and materials were purchased from Sigma-Aldrich/Merck (Darmstadt, Germany) with the exception of commercial cell media, which were purchased from Gibco (Thermo Fisher Scientific, Waltham, MA, USA).

### 2.1 Tissue and cell samples

#### 2.1.1 Krebs-Ringer buffer recipes

Krebs-Ringer Bicarbonate Buffer powder with 1800 mg/L glucose was prepared according to manufacturer’s instructions. Sodium pyruvate, sodium fumarate, and sodium L-glutamate were supplemented to 5 mM. The Krebs-Ringer (KR) buffer was perfused with carbogen for 20 min and the pH adjusted to 7.3. Cell-culture grade penicillin/streptomycin was added to 1x. For KR containing Ca^2+^ and Mg^2+^ (KR+), cell culture grade CaCl_2_ solution was first added to 1.25 mM. For Ca^2+^ and Mg^2+^-free KR (KR-), we proceeded straight to an osmolarity adjustment to ≈ 290 mOsmol, using NaCl and KCl, followed by sterile filtration. The buffers were stored for up to one week at 4°C. Tissue biopsies were collected in freshly carbogen-perfused KR+. For cell dissociation from tissue, dithiothreitol (to 1 mM) and EDTA (to 3 mM) were added to KR-, followed by sterile filtration. For centrifugation steps, a 1% (V/V) Matrigel solution (Corning, Corning, NY, USA) in KR+ was prepared, followed by sterile filtration. This solution was used to prepare 20% and 40% (V/V) solutions of Percoll (Cytiva, Marlborough, MA, USA), layered to generate a density gradient for centrifugation.

#### 2.1.2 Human colon biopsy collection

Biopsy donors (S1:1A) underwent surgical resection of primary colorectal cancer. From the resection margin, a small amount of macroscopically healthy, tumor-adjacent colon tissue was obtained. All donors gave written informed consent in accordance with the Swedish Ethical Review Authority (Etikprövningsmyndigheten), Sweden, ethics permit number 2010-157 with addendums 2010-157-1 and 2020-05754, and permit number 2023-01524-01. Biopsies, pseudonymized before analysis to protect donor details, were rapidly transported to the lab in oxygenated, ice-cold KR+. The mucosa was dissected out. A portion was snap-frozen and transferred to storage at -150°C until proteomics analysis. The remainder was immediately used to isolate primary cells and/or establish colonoid lines.

#### 2.1.3 Primary colonic epithelial cell isolation

Cells were isolated according to an adapted protocol based on van der Post and Hansson [22], followed by density gradient centrifugation. Approx. 1 g of mucosa was rinsed with KR+, transferred to a 50 mL Falcon tube containing 10 mL of KR-, and shaken at 350 rpm for 30 min on ice. The mucosa was then transferred to a new 50 mL Falcon tube containing 10 mL of KR- and shaken at 350 rpm for 60 min at 37°C [22]. The tube was vortexed 5-10x for 10 s at medium speed, after which remaining tissue was carefully removed. The cell suspension was strained with a 70 µm, then a 40 µm cell strainer. All centrifugation steps proceeded at 300 rcf for 10 min at 4°C [22]. Following the first centrifugation step, the cells were resuspended in 3 mL of 20% Percoll. This suspension was layered on top of 3 mL of 40% Percoll in a 15 mL Falcon tube and centrifuged. The top phase was carefully aspirated, transferred to a new Falcon tube, and diluted with 1% Matrigel (Corning) in KR+ to 8 mL. The cells were washed twice with buffer and once with phosphate buffered saline (PBS) prior to being flash-frozen and stored at -150°C until proteomics analysis.

#### 2.1.4 Colonoid culture

Colonoid lines were established and maintained as previously described for enteroids [34] following an adapted protocol. Colonoids were established and maintained with a passaging ratio of approx. 1:4. For transfer to suspension culture, culture medium was replaced with 0.5 mL of ice-cold Cell Recovery Solution (Corning) and incubated for 1 min. Well contents were transferred to 15 mL Falcon tubes and incubated at 20 rpm on a nutating table on ice for 1 h. After allowing the colonoids to settle by gravity, the supernatant was removed and the organoids were carefully washed with Dulbecco’s Modified Eagle Medium (DMEM)/F12 supplemented with 0.25% bovine serum albumin (BSA). The colonoids were again allowed to settle by gravity before removing the supernatant and taking them up in Human Organoid Growth Medium (OGM, STEMCELL Technologies, Vancouver, BC, Canada) or Human Organoid Differentiation Medium (ODM, STEMCELL Technologies, supplemented with 10 µM DAPT). To obtain basal-out colonoids, the medium was supplemented with 7.5% Matrigel (Corning), otherwise apical-out colonoids were obtained. 500 µL/well of the colonoid suspension were then transferred to an ultra-low attachment 24-well plate (Corning) and maintained at 37°C, 5% CO_2_. After 1-3 days in suspension culture (4-6 days post-passaging), the colonoids were harvested by carefully rinsing 3x with PBS on 25 µm strainers (apical-out colonoids) or 40 µm cell strainers (basal-out colonoids) from Funakoshi (Tokyo, Japan) [42]. Experiments were conducted at colonoid passages 4-14.

#### 2.1.5 HEK293 cell culture

Human embryonic kidney 293 (HEK293 cells) were cultured as previously described [38]. Mycoplasma-tested HEK293 cells were obtained from AstraZeneca (Mölndal, Sweden) and cultured as described in the literature [43]. For experiments, performed at passages 18 and 21, cells were seeded at 0.5 million/well in antibiotic-free and phenol-red free DMEM supplemented with 10% fetal bovine serum in CellBIND 24-well plates (Corning) 48 h prior to use.

### 2.2 Microscopy

Unless otherwise stated, colonoid samples were washed with DMEM/F12 + 0.25% BSA on strainers (25 µm for apical-out, 40 µm for basal-out samples) prior to imaging. All reagent dilutions were done in PBS or Hanks’ Balanced Salt Solution (HBSS). Colonoids were allowed to settle by gravity between washing/transfer steps.

#### 2.2.1 Image analysis

ImageJ (version 1.54) was used for image analysis.

#### 2.2.2 Cell viability assay for primary cells

Apoptosis activity in primary cells was measured using CaspGLOW Fluorescein Active Caspase Staining Kits from BioVision (Milpitas, CA, USA), following the protocol provided by the supplier. Briefly, fluorescent markers for activated caspase-3 and -8 were incubated with the primary cells at a temperature of 37°C for 2 hours. After incubation, 20 μL of the cell suspension was analyzed using a Cellometer Vision CBA, using the VB-535-402 filter (Revvity, Waltham, MA, USA).

#### 2.2.3 Size and shape determination

Images of colonoids were captured on an Eclipse TS100 Inverted Routine Microscope with a DS-Fi2 camera mount microscope (10x/0.25 objective) running NIS-Elements F (version 4.00.00) software (all from Nikon, Tokyo, Japan). Size and shape were determined as previously described for enteroids [29], determining cross-sectional area, Feret’s diameter, and circularity by masking and segmenting in ImageJ.

#### 2.2.4 C1-BODIPY-C12 kinetics

Three wells of colonoids were transferred to a falcon tube in 100 µL of PBS, before removing 50 µL and adding 50 µL 5 µM C1-BODIPY-C12 in 5 µM BSA in PBS. Colonoids were then live-imaged in sandwich microscopy slides over 30 min on a custom-built microscope based on an Eclipse Ti2 body (Nikon) [34]. A 10x/0.45 Plan Apo Lambda air objective (Nikon) and a back-lit sCMOS camera with a physical pixel size of 11 μm (Prime 95B, Photometrics, Tucson, AZ, USA) and a final image pixel size of 1.12 μm were used [44]. The imaging chamber was maintained at 37°C and 5% CO2 in a moisturized atmosphere. Widefield images were acquired by differential interference contrast (DIC). Fluorescence imaging relied on the Spectra-X excitation light engine (Lumencor, Beaverton, OR, USA) and emission collection via quadruple band pass filters (89402 & 89403; Chroma Technology Corp, Bellows Falls, VT, USA) and a spinning disk module (X-light V2, CrestOptics, Rome, Italy).

#### 2.2.5 Fluorescence-based immunocytochemistry (ICC) stainings

Colonoids were fixed, stained, and imaged as described in Tzioufa et al. [29], using the staining reagents listed in S1:2A. Briefly, colonoids were fixed with paraformaldehyde, permeabilized with saponin, and incubated overnight with primary antibodies prepared in blocking/staining buffer prior to washing and treatment with secondary antibodies. After washing and mounting, colonoids were imaged using a Zeiss (Oberkochen, Germany) LSM800 confocal microscope equipped with a 20x objective and a 40x immersion oil objective. ICC images were used to quantify how many colonoids per type displayed the desired polarity, represented as a percentage of 100 randomly selected colonoids (compare Fig. 1E).

**Fig. 1:**
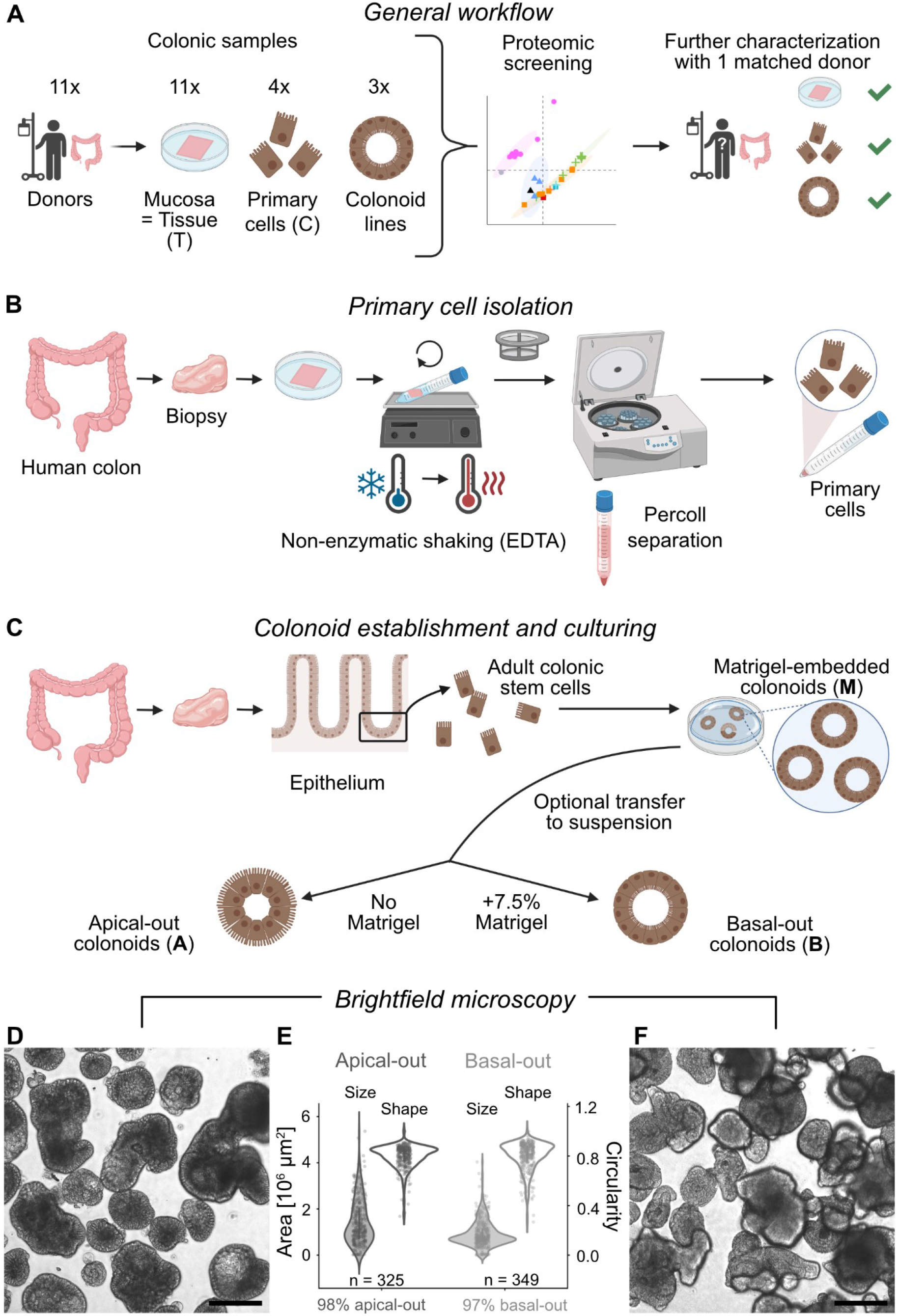
Generation of donor-matched colonic mucosal tissue, primary cells, and colonoid lines. (A) General workflow. (B) Primary colonic epithelial cell isolation. (C) Colonoid establishment and culturing. (D, F) Micrographs (scale bar 200 µm) and (E) size and shape of apical-out and basal-out suspended differentiated colonoids, n denotes the number of colonoids included in the size and shape analyses Graphics in A-C produced in BioRender.

#### 2.2.6 Goblet cell determination

Goblet cell proportion was determined using the images from MUC2 ICC stainings, identifying cells via their nuclei (DAPI channel). The percentage is expressed as the mean number of cells filled with MUC2 (MUC2 channel) vs. the total mean number of cells.

#### 2.2.7 Scanning electron microscopy (SEM)

SEM samples were processed and handled as described in Tzioufa et al. [29]. Briefly, the samples were fixed overnight, washed, dehydrated, mounted, and platinum coated. Samples were then imaged by field-emission scanning electron microscopy (FESEM, Zeiss Merlin) using an Inlens secondary electron detector.

#### 2.2.8 Breast Cancer Resistance Protein (BCRP) and barrier integrity assay

Per sample, 3 wells of colonoids were transferred to 0.5 mL tubes in DMEM/F12 + 0.25% BSA. The supernatant was removed and replaced with 400 µL of 100 µM of the BCRP substrate Lucifer Yellow (LY), optionally supplemented with 0.1% saponin. Samples were incubated for 2 h at 37°C at 100 rpm. The supernatant was then removed and samples washed twice with HBSS. Samples were resuspended in DMEM/F12 + 0.25% BSA and imaged in widefield and confocal mode on the custom-built microscope described in section 2.2.4. Data were analyzed by manually drawing regions of interest (ROIs) around colonoids via touchscreen input, creating a mask for the colonoid area. For each colonoid, a matched background area was selected by copying the mask to the adjacent background. This background area was used to correct for background fluorescence, see S1:3A for further details.

### 2.3 Plate reader assays

#### 2.3.1 ATP viability assay

A CellTiter Glo 2.0 kit was purchased from Promega (Madison, WI, USA) and used according to manufacturer’s instructions, using 2 colonoid wells per replicate in a white 96-well plate. Overnight treatment with toxin (25 µM temozolomide) was used to generate positive controls for apoptosis. The assay was read out with a Spark plate reader running SparkControl software (version 2.2) (Tecan, Männedorf, Switzerland).

#### 2.3.2 P-glycoprotein (P-gp) assay

Colonoid samples were incubated with 1 µM calcein AM in ODM, optionally supplemented with 1 µM of the P-gp inhibitor elacridar, for 20 h [45]. Per replicate, 2 wells of colonoids were transferred directly to 0.5 mL tubes on ice and rapidly rinsed with ice-cold HBSS. The colonoids were taken up in 200 µL of HBSS and transferred to a black-wall clear bottom 96-well plate maintained at RT. Fluorescence (excitation at 494 nm, emission at 514 nm) was measured at 37°C over 30 min on a Spark plate reader running SparkControl software (version 2.2) (Tecan). Cells in each well were then lysed and analyzed using a Nucleocounter NC-200 (ChemoMetec, Allerød, Denmark).

#### 2.3.3 Phospholipid content determination

Cell homogenates from cellular unbound fraction (f_u_) assays were used. Phospholipid concentrations were determined using a MAK122 kit according to manufacturer’s instructions. The assay was read out with a Safire 2 plate reader running Magellan (version 7.2) and SparkControl software (version 2.2) (Tecan).

### 2.4 Compound selection

A chemically representative small-molecule drug panel (S1:4) incorporating reference compounds [38] as well as drugs with publicly available *in vivo* colonic absorption data [3] (S1:4C) was used for determining intracellular bioavailability (see section 2.5.4). Chemical space analysis was performed using the FactoMineR package in R [46]. The approved drug database available via DrugBank [47] was subjected to principal component analysis (PCA). The eigenvalues were plotted using GraphPad Prism 10 (San Diego, CA, USA).

### 2.5 UHPLC-MS/MS-based assays

Compound concentrations were determined by UHPLC-MS/MS using an Acquity UHPLC coupled to a XEVO TQ-S micro triple quadrupole mass spectrometer in positive electrospray ionization mode (Waters, Milford, MA, USA). For chromatographic separation, a T3 HSS 2.1×50 mm 1.7 μm column (Waters) was used. The injection volume was 5 µL. Multiple reaction monitoring methods were optimized for each compound, metabolite, and internal standard (warfarin). Method parameters are described in S1:5A-B. Compounds were considered to be quantified if stability and mass balance criteria of 75%-125% were met.

#### 2.5.1 Global proteomics

Proteomic screening of donor samples was performed using the following replicates per donor and sample type: a single injection per tissue piece, triplicate injection per primary cell isolate (due to the low number of primary cell isolates of n = 4), and duplicate injection of each possible combination of colonoid medium and configuration (compare Fig. 2A). All further characterization of donor 4-derived samples was performed in technical triplicate (repeat injections on LC-MS/MS) of the following biological replicates: 2 tissue pieces, 2 primary cell isolates, and 3 biological replicates of each possible combination of colonoid medium and configuration. For mucosa samples, approx. 0.1 g was minced and homogenized using an Ultra-Turrax disperser (IKA-Werke, Staufen im Breisgau, Germany) in 300 µL of lysis buffer. Alternatively, Precellys 0.5 mL soft tissue homogenizing kits (Bertin Technologies, Montigny-le-Bretonneux, France) and 100 µL of lysis buffer [48,49] was used for homogenization. Primary colonic epithelial cells and colonoids were lysed directly by addition of 100 µL of lysis buffer. Single-pot, solid-phase-enhanced sample preparation (SP3) was performed [48,49], followed by UHPLC-MS/MS analysis as previously described [50]. MaxQuant (version 2.6.2.0) [51] was used for raw data analysis, matching against the UniProt database [52], followed by an in-house pipeline in R (see S2AO for additional details). Proteins with a minimum of 3 identified razor/unique peptides were quantified using the Total Protein Approach [50]. The mass spectrometry proteomics data have been deposited to the ProteomeXchange Consortium via the PRIDE [53] partner repository with the dataset identifier PXD086179.

**Fig. 2:**
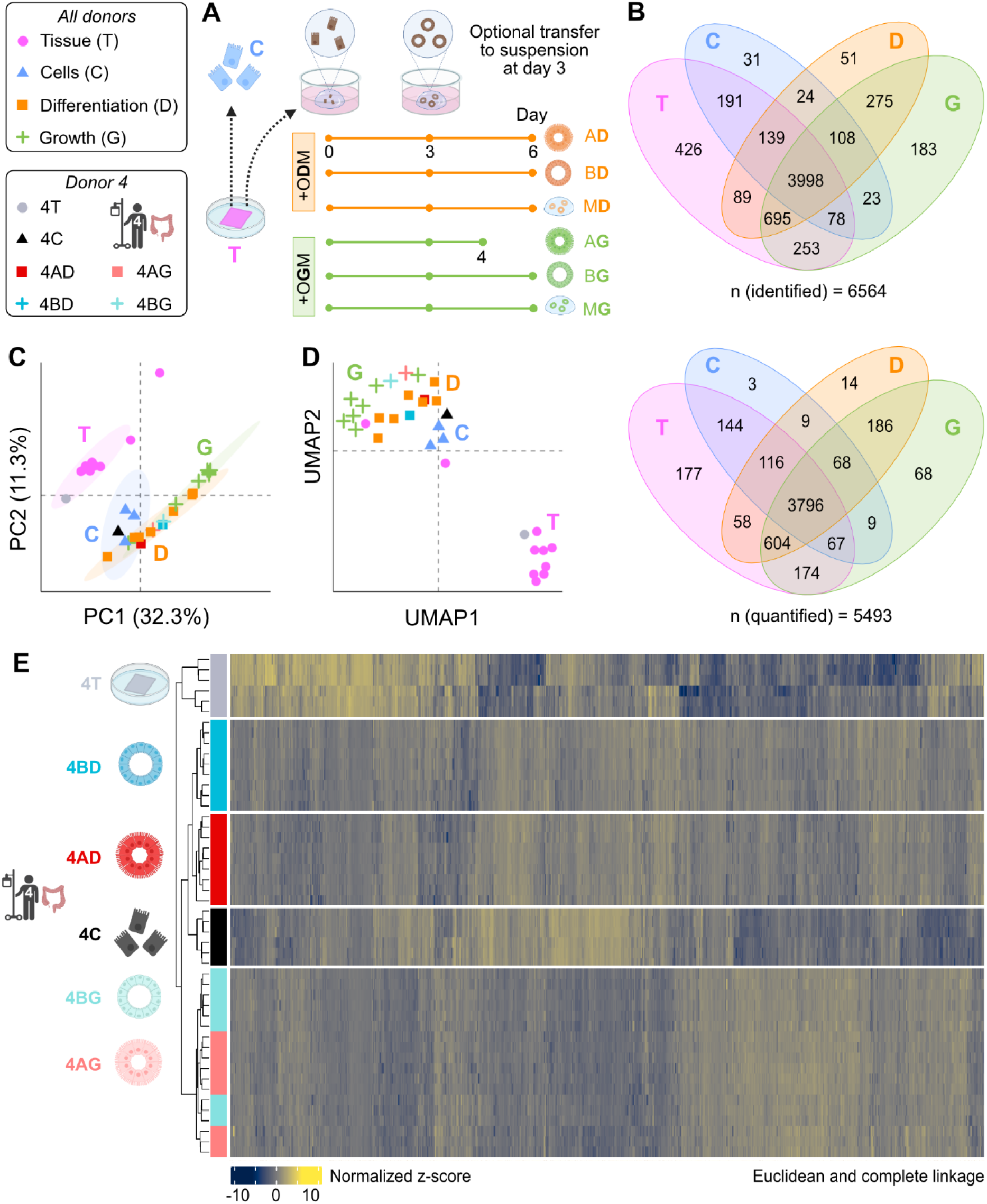
Proteomic screening to identify a suitable colonoid line and culture conditions. (A) Colonoid culturing timeline. (B) Venn diagrams of identified and quantified proteins across donors and sample types. (C) PCA with 95% confidence ellipses for each cluster and (D) UMAP across donors and sample types, with donor 4 highlighted. (E) Hierarchical clustering (Euclidean, complete linkage) of tissue (4T), primary epithelial cells (4C), and suspended colonoids (4AD, 4BD, 4AG, 4BG) from donor 4. Graphics in A and E produced in BioRender.

#### 2.5.2 Targeted proteomics

Protein quantification was conducted in accordance with previously published methodology [54,55]. Frozen tissue (≈ 10 mg) was ground using a ball mill (Micro-dismembrator S, Sartorius, Göttingen, Germany). Pulverized tissue and organoids were incubated for one hour with lysis buffer, after which the resulting protein concentration was determined by BCA assay (Thermo Fisher Scientific). The lysate was then proteolyzed using trypsin. Surrogate peptides and internal standard peptides were precipitated using triple X proteomics (TXP) antibodies. Subsequently, peptides were eluted and quantified using modified 6-minute parallel reaction monitoring (PRM) LC–MS methods [56,57]. Raw data were processed using TraceFinder 4.1 (Thermo Fisher Scientific). Peptide amounts were calculated by forming the ratios of the integrated peaks of the endogenous peptides and the isotope-labeled standards, with protein quantities reported in normalized units (fmol/µg).

#### 2.5.3 Metabolism assay

Metabolic cocktails (midazolam, acetaminophen, and 4-methylumbelliferone) were diluted to 1 µM in HBSS. Per replicate, 2 wells of colonoids were transferred to 3 mL glass tubes in 500 µL of probe mix. As a negative control, colonoids were incubated for 5 min at 95°C and cooled on ice before the experiment. Samples were incubated at 37°C at 300 rpm for 60 min. At each timepoint, 80 µL were sampled (after ensuring homogenous dispersion of colonoids) and lysed with 120 µL of 10 nM warfarin (internal standard) in 100% acetonitrile or methanol prior to UHPLC-MS/MS analysis. Colonoids remaining in each tube after 60 min were lysed and analyzed using a Nucleocounter NC-200 (ChemoMetec) to determine cell count.

#### 2.5.4 Intracellular bioavailability (F_ic_)

Cell assays were performed using cassette dosing (10 µM compound) with all parameters calculated as described in Mateus et al. [38]. The equations for these calculations are listed in S1:6A. Briefly, cellular accumulation (Kp) was determined in cells over time and the intracellular unbound fraction (f_u_) was determined via rapid equilibrium dialysis (RED) of cell homogenates. Multiplication of these parameters yielded the intracellular bioavailability (F_ic_). For colonoid samples, the following adaptation was made: The EDTA (or organoid cavity) fraction (see section 2.5.4.1) was handled as part of the extracellular fraction in Kp calculations.

##### 2.5.4.1 Cellular accumulation (Kp) assay

To minimize breakage, 1000 µL pipette tips were used for all colonoid handling steps. During all washing steps, colonoids were allowed to settle by gravity for 2 min. Per replicate and timepoint, 3 wells of colonoids were used. They were rinsed thrice with HBSS on strainers (25 µm for apical-out, 40 µm for basal-out samples) before being transferred to 0.5 mL Eppendorf tubes in 500 µL 0.1% BSA in HBSS. After allowing the colonoids to settle by gravity for 2 min, 450 µL of liquid was carefully removed. For the washing step, performed twice, HBSS was carefully added up to the 0.5 mL fill line, followed by removal of 450 µL of liquid. To the 50 µL remaining in the tube, 450 µL of 10 µM compound solution in HBSS was added. From the supernatant in each tube, 20 µL was sampled to determine the respective initial drug concentration. The tubes were then placed on an orbital shaker in a shallow 37°C water bath. After incubation for a total of 15, 30, or 60 minutes, the tubes were placed on ice and 20 µL was sampled from the supernatant to determine its drug concentration. The supernatant (410 µL) was removed and discarded. The colonoids were then washed as described above. Cell culture grade EDTA in HBSS (10 mM) was then carefully added up to the 0.5 mL fill line. To determine the initial concentration in EDTA solution, 20 µL was sampled. The samples were kept on ice for 15 min, after which they were pipetted up and down 10x to ensure access to the organoid cavity. Cells were spun down for 1 min at 3000 rpm using a benchtop microcentrifuge (MiniSpin Plus, Eppendorf) after which 20 µL was sampled from the supernatant to determine the concentration in the EDTA solution. A volume of 410 µL of EDTA solution was then removed and discarded. The samples were again washed as described above, this time adding HBSS up to the 0.5 mL fill line. Cells were then spun down for 1 min at 3000 rpm using the previously described benchtop microcentrifuge, after which 450 µL of supernatant was removed and discarded. Samples were then lysed by adding 450 µL 10 nM warfarin in ACN/H_2_O (60/40), pipetting up and down 10x to ensure lysis. Samples were then stored upright in a -20°C freezer for 3 h prior to spinning down cells for 1 min at 3000 rpm using the benchtop microcentrifuge. To determine the cell concentration, 200 µL was sampled, after which all remaining liquid was carefully removed. Samples were then subjected to protein determination and processed as described in Mateus et al. [38].

##### 2.5.4.2 Cellular unbound fraction (f_u,cell_) assay

Up to 20 wells of colonoids were pooled and homogenized in 1.5 mL of HBSS using a Vibra-Cell ultrasonication microtip (Sonics & Materials, Newtown, CT, USA) at 20% intensity for 20 s. Samples were then processed as described in Mateus et al. [38].

#### 2.5.5 Lipidomics

Targeted lipidomics analysis was performed by the Swedish Metabolomics Center, part of the national SciLifeLab infrastructure, in Umeå, Sweden. Briefly, samples were prepared in triplicate. Lipids were extracted according to published methodology [58] and analyzed via UHPLC-MS/MS as described in [59]. The data were then preprocessed using ProFinder 10.0 (MassHunter, Agilent, Santa Clara, CA, USA), performing data quality control using SIMCA version 18 (Sartorius Stedim Biotech, Aubagne, France). Additional details are given in S1:7A.

### 2.6 Statistical analysis

Experiments were performed in triplicate on two independent occasions unless otherwise stated. Visualization was performed in GraphPad Prism 10 or R. A p-value < 0.05 was considered significant.

#### 2.6.1 Global proteomics

Differential protein expression analysis in mass spectrometry (DEqMS) [60] was used as a statistical method, set to a log2 fold change threshold of 1. Overrepresentation analysis (ORA) was run against the Reactome [61] database. Gene set enrichment analysis (GSEA) [62–69] was run against the Gene Ontology (including PANTHER) [70–72] and Reactome [61] databases. For both forms of enrichment analysis, multiple testing correction of p-values was performed according to Benjamini-Hochberg.

#### 2.6.2 Lucifer Yellow (LY) assay

A Kruskal-Wallis test with Dunn’s post-hoc test and Holm’s multiple comparison correction was used.

#### 2.6.3 Intracellular bioavailability (F_ic_) ratio

A Wald test was used to check for a statistically distinguishable difference in upper and lower plateaus based on the curve fit. A Welch two-sample t-test and a Wilcoxon rank-sum test were used to analyze differences in group means based on grouping compounds by these two plateaus.

## 3 Results and discussion

### 3.1 Generation of donor-matched colonic mucosal tissue, primary cells, and colonoid lines

Colonic biopsies were collected from 11 donors by sampling macroscopically healthy tumor-adjacent tissue (Fig. 1A, S1:1A), with different colon segments, donor age ranges, and both biological sexes represented.

From 4 of these biopsies, primary cells were isolated via a gentle, non-enzymatic colonic epithelial cell isolation procedure (Fig. 1A, 1B, S1:1A-C). A lower degree of surface enlargement, tighter junctions, and higher microbial load in the colon [1,73] contributed to an overall lower primary cell yield per biopsy relative to small intestinal samples (S1:1B) [29,74]. The biopsies were also small, often only yielding material for either primary cell isolation or organoid establishment.

Still, from 3 of the biopsies colonoid lines were established from adult colonic stem cells (Fig. 1A), yielding highly viable colonoids of a microscopically uniform differentiation status conducive to reproducibility. Specifically, the colonoids were approximately spherical and consisted exclusively of clear (vs. darkened) cells forming a narrow epithelial band (S1:1D). Of note, this starting material constituted a marked improvement over reports in the literature of colonoids visibly having begun to differentiate to varying degrees prior to exposure to differentiation medium [75].

Through transfer to suspension culture (Fig. 1A, 1C), apical- (Fig. 1D) and basal-out (Fig. 1F) colonoids were obtained, shown here in their differentiated state. In Matrigel-free suspension culture, 98% of colonoids were apical-out. Conversely, in Matrigel-supplemented suspension culture, 97% of colonoids were basal-out (Fig. 1E). Due to the Matrigel supplementation [76], these basal-out colonoids were prone to aggregation (Fig. 1F), making them more challenging to handle. Both types of colonoids exhibited a high degree of size uniformity, with apical-out colonoids having a ≈ 1.6-fold greater cross-sectional area. Both types were also approximately spherical at average circularities of ≈ 0.8 (Fig. 1E), mimicking the curved geometry found in native epithelium [29].

Despite comparable circularities, the colonoids’ shape profiles as viewed by microscopy differed. Protrusions were apparent at the surface of basal-out colonoids (Fig. 1F), creating crypt-like invaginations in the organoid interior. The formation of these invaginations likely restricted growth to some extent, reflected in the smaller overall size (Fig. 1E), as loss of curvature results in a less proliferative phenotype [29,77]. In contrast, apical-out colonoids possessed a relatively smooth profile (Fig. 1D), similar to recent observations on jejunal enteroids [29]. Despite a lack of visible cryptlike structures in the apical-out configuration, both configurations were revealed to produce cryptlike cell populations, discussed in section 3.5. Compared with other human intestinal 3D organoid ADME-T models reported in the literature (e.g., [14,29]), our colonoids were exceptionally homogenous in both size and shape, constituting an important step towards reproducibility and standardization.

### 3.2 Proteomic screening to identify the most suitable colonoid line and culture conditions

#### 3.2.1 Screening of matched colonic samples visualizes relative sample variance and similarity

The three colonoid lines were subjected to screening to identify the culturing conditions leading to the most native-like proteome. When passaging, colonoids were split into either differentiation (-D) or growth (-G) medium. They were then transferred to suspension culture on day 3 to obtain suspended apical-out (A-) and basal-out (B-) colonoids or kept in Matrigel to obtain Matrigel-embedded (M-) colonoids. Colonoids were harvested on day 6 with the exception of AG samples, which were harvested on day 4 (Fig. 2A). Thus, suspended as well as Matrigel-embedded colonoids were obtained.

Global proteomic analysis of excised mucosal tissues (T), isolated primary cells (C), and all types of colonoids (AD, BD, MD, AG, BG, and MG) yielded high numbers of total identified (n = 6564) and quantified proteins (n = 5493). Notably, about half of these proteins were shared across all sample types (n = 3998 and n = 3796, respectively). Tissue had the highest number of unique proteins (426 identified and 177 quantified) (Fig. 2B), setting it apart from the other sample types. Further trends in similarity emerged through use of dimensionality reduction techniques, where distinct clusters were observed for each sample type in both PCA (Fig. 2C) and uniform manifold approximation and projection (UMAP) (Fig. 2D). Both techniques revealed tissues clustering separately whereas primary cells and colonoids had low variance (Fig. 2C) and high similarity (Fig. 2D) relative to one another. Among the colonoids, the differentiated samples appeared to be more similar to the primary cells overall (Fig. 2D).

#### 3.2.2 Tissue, primary cells, and suspended colonoids from donor 4 selected for further characterization

Of all available matched samples, those from donor 4 (denoted by the prefix 4-) were selected for further characterization. This selection was made based on their overall within-group representativeness per sample type as well as the availability of all sample types (Fig. 2C–2E, S1:1, S1:8A). We acknowledge that donor 4 was male (S1:1A), which constitutes a limitation where sex-specific phenotypes are relevant. For this first-of-kind study, we accepted this limitation and aimed to pinpoint the most suitable colonoid types for further characterization.

Matrigel-embedded colonoids were similar to suspended ones but contained a greater amount of residual extracellular matrix proteins. Though it did not impact overall clustering by PCA or UMAP (S1:8B-C), we sought to limit human proteomic contamination introduced by the murine-derived Matrigel [78]. Additionally, Matrigel-embedded colonoids neither yielded direct membrane access nor the possibility of an apical-out configuration. For these reasons, only suspended colonoids were characterized further. Hierarchical clustering confirmed 4T as a separate cluster while all suspended colonoid types clustered together with 4C. Of these, 4AD was closest to 4C, followed by 4BD. Next came 4AG and 4BG, between which there appeared to be little difference due to intermingled clusters (Fig. 2E).

### 3.3 Comparative analysis of differentially expressed proteins highlights differences in cellular function

#### 3.3.1 Volcano plots indicate changes to cellular reorganization in primary cells vs. colonoids

These matched samples were subjected to differential protein expression analysis, visualized in the form of volcano plots. Sample types were compared to the native epithelial reference of 4C (Fig. 3A-C). The total number of statistically significant differentially expressed proteins (matched by their gene names) was highest in 4T (n ≈ 800), followed by 4AG and 4BG (both at n ≈ 750). Next was 4BD (n ≈ 500), with 4AD having the least divergent proteome (n ≈ 400) compared to 4C (Fig. 3B).

**Fig. 3:**
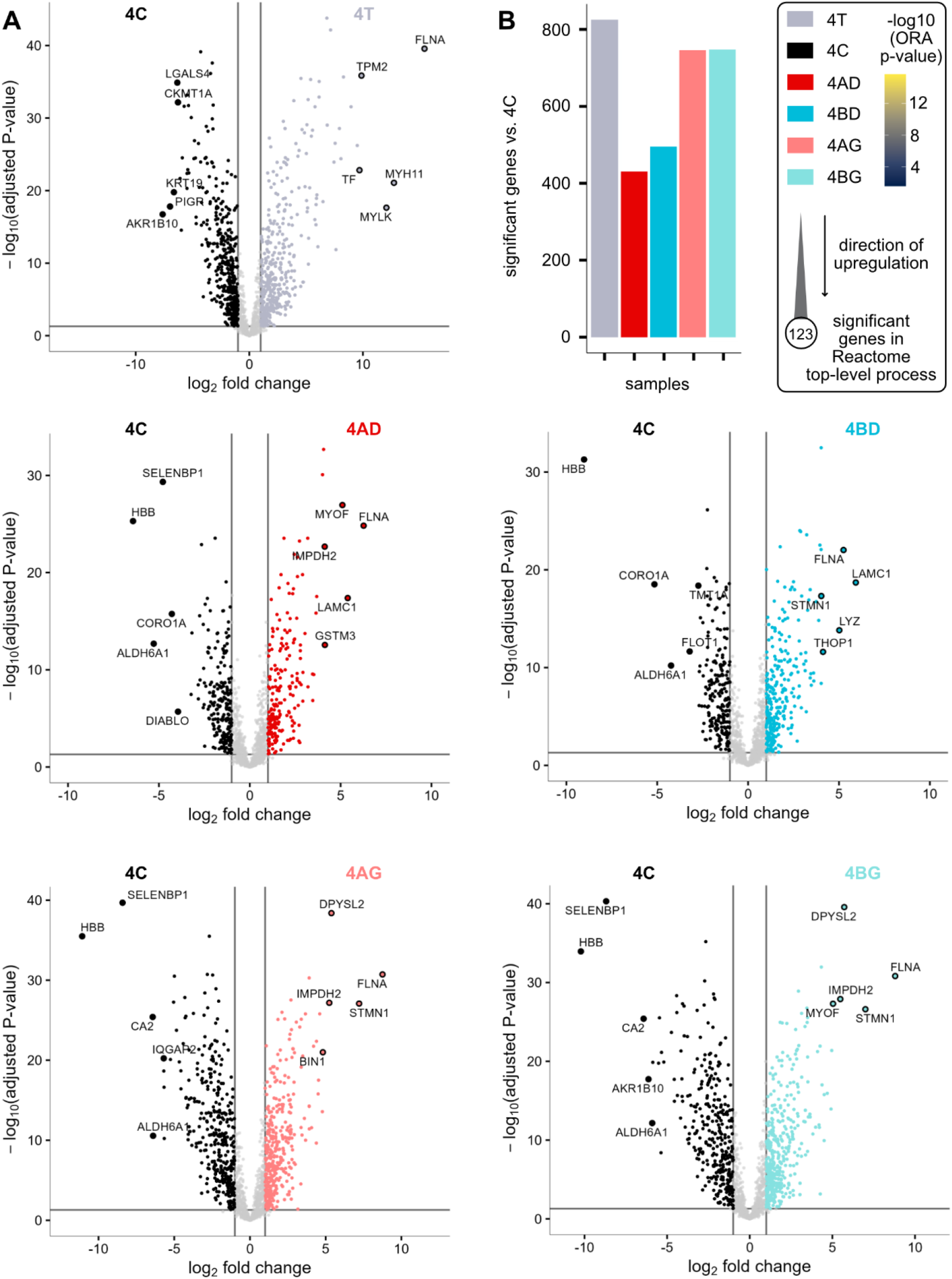
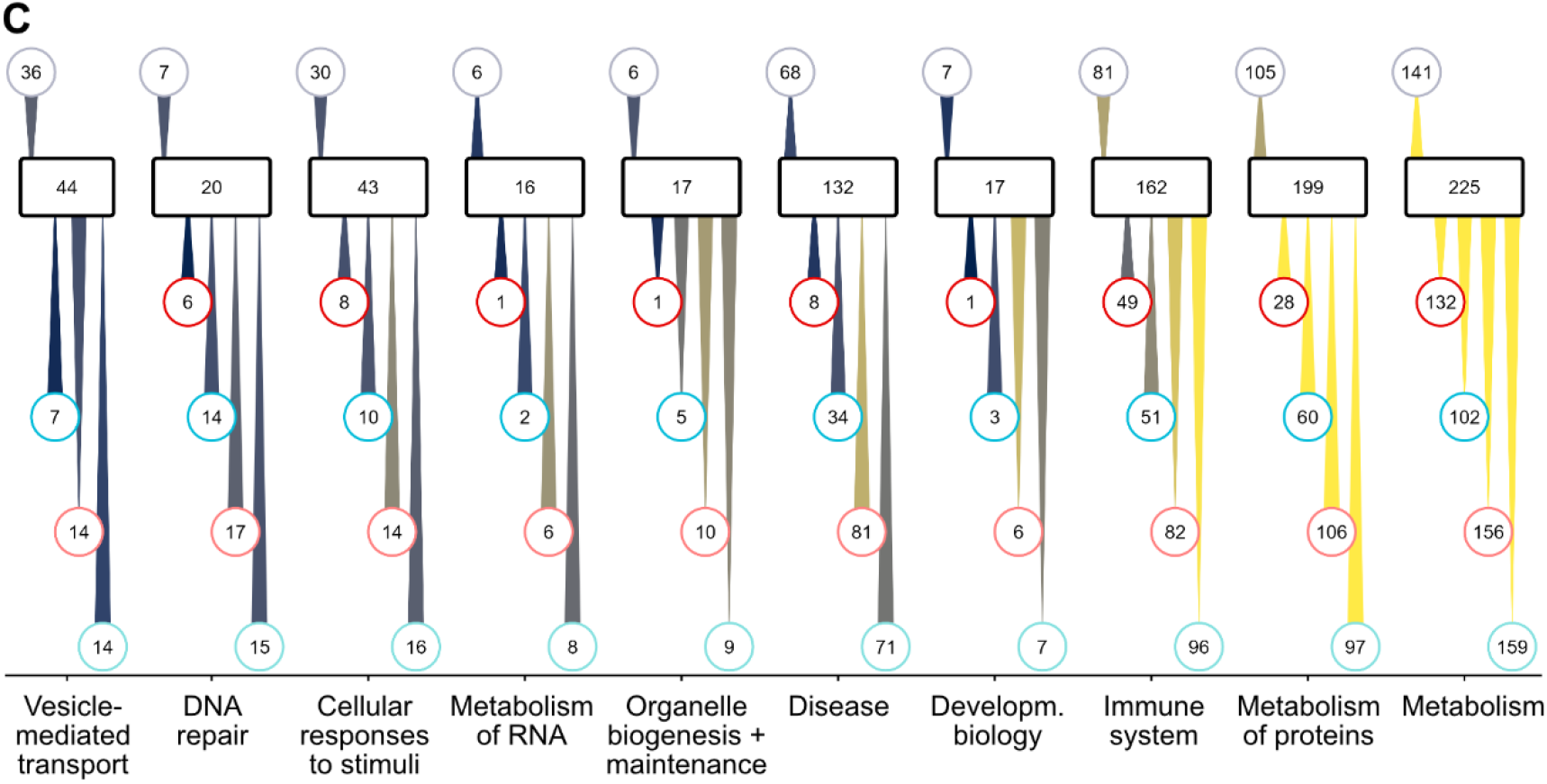
Comparative analysis of differentially expressed proteins highlights differences in cellular function. (A) Volcano plots of 4T, 4AD, 4BD, 4AG, and 4BG compared to 4C. (B) Barplot of the total number of significant proteins (matched via their genes) in each volcano plot. (C) Longitudinal appreciable volcano analysis (LAVA) based on preceding volcano plots with top 10 Reactome top-level processes shown.

Focusing first on only the top five significant proteins in each pair of samples, volcano plots indicated important trends: FLNA (filamin A), an actin crosslinker involved in maintaining apicobasal polarity [79], was overexpressed in all sample types relative to 4C. In differentiated colonoids, LAMC1 (laminin subunit gamma 1), crucial in the assembly of basement membranes [79], was a shared overexpressed protein. STMN1 (stathmin 1), a microtubule destabilizer abundant in proliferative cells, was overexpressed in all colonoid types except 4AD (Fig. 3A). Colonoids in growth medium overexpressed DPYSL2 (dihydropyrimidinase-like protein 2), which promotes microtubule assembly [79]. In short, we observed a trend in changes to proteins relating to cellular reorganization.

In the context of the isolation procedure necessary for obtaining the primary cell reference, such reorganization is expected. During isolation, the primary cells were removed from the basement membrane, resulting in a loss of laminins (S1:8D), and dissociated into a single-cell suspension. This loss of basolateral contact points led to rounded cells, or a loss of apicobasal polarity, over time (S1:1C), implying substantial cellular reorganization processes.

This trend of reorganization remained even when accounting for all significantly differentially expressed proteins. The corresponding gene sets were queried against the Reactome database [61] for overrepresentation, returning the top 10 top-level processes. We termed this plot type “Longitudinal Appreciable Volcano Analysis” (LAVA), as it visualizes key affected processes, the number of associated significant genes per process, and the “flow” in terms of the direction of regulation (denoted by the connecting triangle) relative to the reference 4C. Processes ranging from vesicle mediated transport to cellular metabolism (excluding xenobiotic metabolism) emerged as significantly altered between samples (Fig. 3C). Important for downstream analysis (see section 3.7), ADME-T related processes (annotated as “Drug ADME” in the Reactome database [61]) were conserved in the colonoids relative to 4C.

#### 3.3.2 Gene set enrichment analysis (GSEA) reveals colonoid differentiation status

Additional between-sample differences hinted at in the LAVA plot were explored in-depth via gene set enrichment analysis (GSEA). In line with the previously observed differential clustering (Fig. 2C-E) and highest number of significantly differentially expressed proteins (genes) (Fig. 3B), 4T was the most LAVA-divergent overall (Fig. 3C). GSEA revealed the enrichment of proteins (genes) relating to hemostasis and signaling between different tissue layers (S2A-H), consistent with 4T not purely consisting of epithelium (discussed further in section 3.5.4).

On the opposite end of the spectrum, 4AD was the least disease-associated colonoid type with only 8 LAVA-upregulated proteins (genes). The colonoids thus accurately recapitulated a healthy state despite their origin in tumor-adjacent, macroscopically healthy tissue, which may exhibit a more disease-like phenotype than truly healthy tissue [80]. Isolating and culturing stem cells derived in this manner seems to restore the healthy phenotype to a large extent, in line with existing literature on tumor-adjacent-tissue-derived colonoids [81]. 4AD also had the fewest changes in proteins (genes) relating to immune processes and metabolism of proteins (n = 49 and n = 28, respectively) (Fig. 3C). GSEA indicated 4AD as having a profile consistent with differentiated epithelium, with an overall focus on protein folding and maturation (S2I-N). Pathway analysis confirmed the retention of proliferative activity compared to the inherently dying 4C (S2O-P).

4BD was the second most similar sample to 4C. This sample had slightly more LAVA-significant proteins (genes) overall than 4AD (288 vs. 234) (Fig. 3C) and a GSEA profile suggesting a proto-version of 4AD, namely an upregulation of RNA translation (S2Q-W).

4AG and 4BG had approximately comparable profiles in both LAVA (Fig. 3C) and GSEA (S2Y-AN), matching the intermingled clustering observed in Fig. 2. Cellular processes yet another step upstream of proteins, namely those relating to RNA modification and translation initiation (S2Y-AN), were enriched in both samples. Though this enrichment agreed with a more stem-like or proliferative phenotype as expected from exposure to growth medium, it also positioned 4AG and 4BG as the least suitable colonoid types for subsequent studies.

### 3.4 Suspended differentiated colonoids have native-like morphology and viability

#### 3.4.1 Morphological characterization confirms 3D geometry and successful eversion

Based on the proteomic characterization above (see section 3.3.2), only 4AD and 4BD, schematically represented in Fig. 4F, were subjected to morphological and viability analysis. In line with brightfield microscopy (Fig. 1D-1F), both colonoid types were confirmed to be approximately spherical (Fig. 4A, 4D, 4G, 4H, 4I), mimicking the 3D geometry of native intestine as discussed above (section 3.1).

**Fig. 4:**
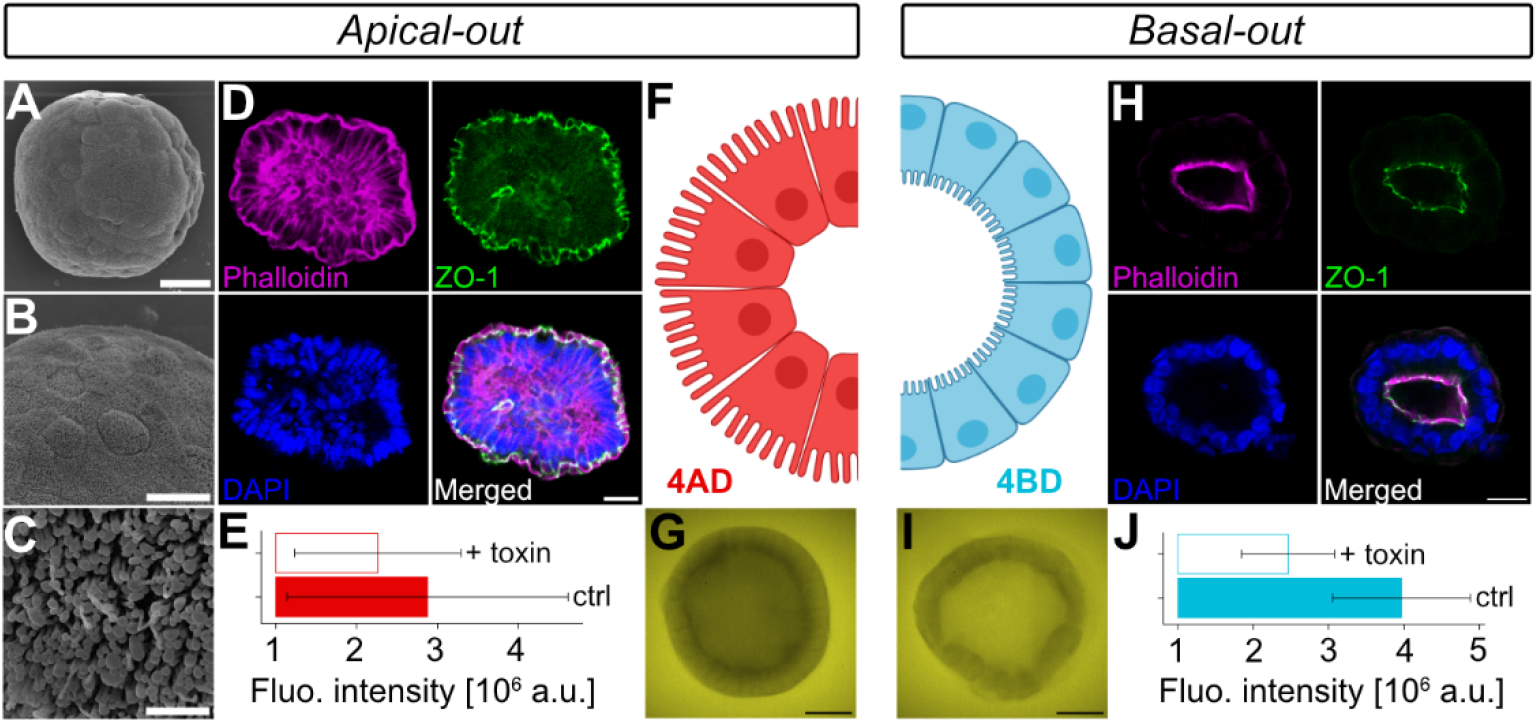
Suspended differentiated colonoids have native-like morphology and viability. (A-C) SEM images of 4AD (scale bars 20, 10, and 1 µm, respectively). (D, H) ICC images of 4AD and 4BD, respectively (scale bars 20 µm). (E, J) ATP viability assay (mean ± SD) for 4AD and 4BD, respectively, with temozolomide used as a toxin. (F) Schematic representation of 4AD and 4BD, produced in BioRender. (G, I) Lucifer yellow exclusion assay in 4AD and 4BD, respectively (scale bars 20 µm).

SEM of 4AD indicated a tight monolayer of individually discernible cells (Fig. 4B) with microvilli (Fig. 4C). For 4BD, similar but inverted morphology was expected, as previously seen in suspended basal-out 3D enteroids [29]. ICC staining confirmed the desired polarity, with actin localizing to the exterior in the case of 4AD (Fig. 4D) and interior in the case of 4BD (Fig. 4H), as denoted by the phalloidin stain. The nuclei (denoted by the DAPI stain) were displaced towards the basal pole, most noticeably so in 4BD (Fig. 4H), aligning with their position *in vivo* [21].

Staining for ZO-1 (zonulin-1, also known as tight junction protein 1), pointed towards an intact barrier function (Fig. 4D, 4H). Exclusion of the small molecular fluorescent permeability marker Lucifer Yellow (LY) by 4AD (Fig. 4G) supported this notion. In the case of 4BD, active transport of LY into the central “lumen” or cavity occurred (Fig. 4I) [29], which is discussed in detail below (see section 3.7.1).

#### 3.4.2 Suspended differentiated colonoids exhibit high viability

For both 4AD and 4BD, an ATP-based viability assay revealed sustained cellular metabolic activity after 24 h, corresponding to a timescale easily accommodating most drug and toxicity assays. 4BD showed slightly higher ATP levels than 4AD, consistent with a slightly more proliferative or less differentiated phenotype (Fig. 4E, 4J).

Due to the high turnover of the colonic epithelium, a small proportion of about 3% of cells entering programmed cell death is inherent even in living tissue [82]. Regrettably, insufficient cells from donor 4 were available to perform an apoptosis assay. Thus, donor 7 primary cells were used, with only a small fraction being dead or dying post-isolation (max. 24% as measured by activated caspase-3, a late apoptosis marker) (S1:2B), indicating high overall viability of the primary cell reference material. The viability of 4C and both colonoid types was additionally supported by low levels of programmed cell death markers, both for general apoptosis (APAF1, BCL2L1, and CASP3/7/8) and those specific to anoikis (ATOP2A3, NAT1, PBK) (S1:2C-D).

### 3.5 Suspended differentiated colonoids contain representative colonic epithelial cell populations

#### 3.5.1 Suspended differentiated colonoids contain all mucosal epithelial cell types

For all colonic epithelial cell types found along the crypt-surface axis (Fig. 5A), we quantified marker proteins (Fig. 5B). Importantly, the two physiologically most abundant cell types were clearly present in our colonoids. Nutrient-absorbing colonocytes and mucus-secreting goblet cells account for ≈ 93% of colonic mucosal cells [1]. As such, confirming their presence and ensuring a representative goblet cell population (Fig. 5B-F, sections 3.5.2 and 3.5.3) were deemed sufficient for the purpose of our studies.

**Fig. 5:**
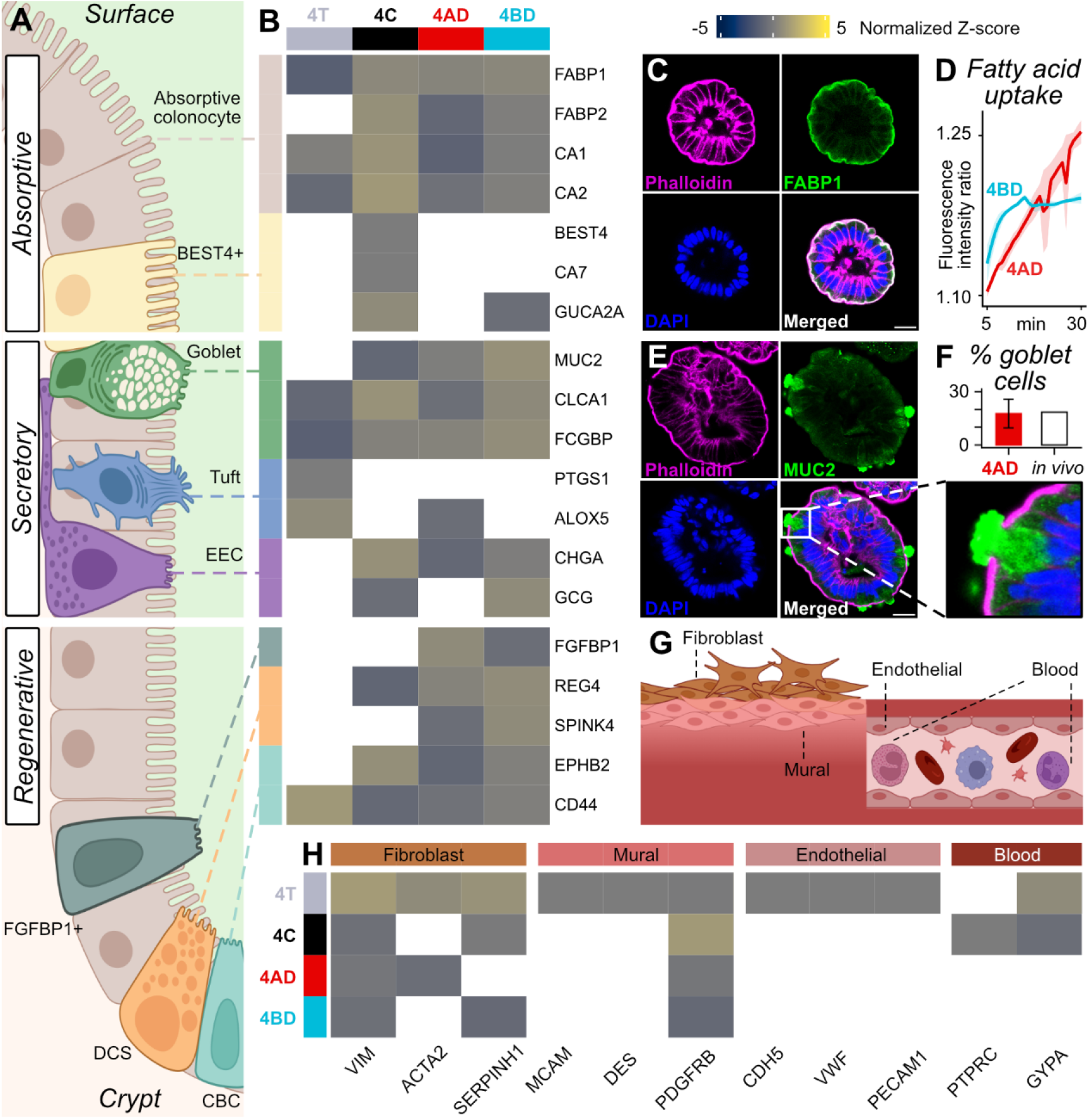
Suspended differentiated colonoids contain representative colonic epithelial cell populations. (A) Schematic representation of the colonic epithelial crypt-surface axis. (B) Heatmap of protein expression of mucosal epithelial cell markers based on global proteomics. (C) ICC image of FABP1 staining in 4AD (scale bar 20 µm). (D) Uptake of C1-BODIPY-C12 in 4AD and 4BD. (E) ICC image of MUC2 staining in 4AD (scale bar 20 µm), inset highlighting extruding mucus plume. (F) Percentage of goblet cells per colonoid in 4AD (based on n = 30 colonoids stained for MUC2) compared to literature. (G) Schematic representation of submucosal cell types present in tissue. (H) Heatmap of protein expression of submucosal cell markers based on global proteomics. Graphics in A and G produced in BioRender.

Though we here focus on the establishment and validation of *in vitro* ADME-T models, the presence of almost all cell types in our colonoids (Fig. 5B, S1:9A) paves the way for extended applications. Such applications could include, e.g., endocrinology (due to the presence of enteroendocrine cells, EECs) or regenerative medicine (due to the presence of the recently described regenerative FGFBP1+ transit-amplifying cells [83] and crypt-base columnar cells, CBCs). Regarding the latter, 4AD seems to be an ideally suited model, as it was enriched for FGFBP1+ cells but CBC-depleted (Fig. 5B, S1:9A). Before other epithelial cell types can be studied in depth in our colonoids, additional work is needed to optimize culturing conditions towards their expression. These cell types include the immunologically relevant tuft cells and the recently discovered absorptive BEST4+ cells (Fig. 5B, S1:9A), whose physiological function is not yet fully understood [84].

#### 3.5.2 Suspended differentiated colonoids display an absorptive phenotype

Often used as a marker of healthy colonocytes [85], FABP1 is a hallmark of an absorptive phenotype [86]. This protein, responsible for shuttling long-chain fatty acids, is also known to shuttle drugs [39,87]. ICC staining for FABP1 confirmed its presence and correct (apical) localization (Fig. 5C). To provide a functional proof of absorptivity, we tracked colonoids’ uptake of a fluorescently labeled long-chain fatty acid, BODIPY-C12. As expected from a more strongly differentiated phenotype, 4AD had a steeper absorption curve than 4BD did (Fig. 5D). Protein expression levels for relevant lipid shuttling machinery (e.g., FABP1 or the C1-BODIPY-C12 uptake transporter SLC27A4) were approximately the same in both 4AD and 4BD (S1:9B). Recent observations in enteroids indicate apical SLC27A4 localization under similar culturing conditions [29], enabling active apical uptake aligning with the high absorptivity seen in 4AD (Fig. 5D).

Though not a focus of our study, we additionally quantified markers of amino acid and glucose transport and processing (S1:9C-D), rounding out an absorptive profile. As expected from cells largely relying on short chain fatty acids for nutrition [1], fatty acid processing machinery dominated in terms of expression levels, followed by amino acid transporters (S1:9B-C). Glucose uptake transporters, known to be weakly expressed in the colon [88,89], were only quantified in 4BD. Sodium-dependent glucose cotransporter 1 (SGLT1), encoded by SLC5A1, was present at low levels (0.067 ± 0.030 fmol/µg) (S1:9D).Trefoil factor 3 (TFF3), a secretory marker [88,89] implicated in glucose regulation [90], was also only quantified in 4BD (S1:9A). Though further exploration is needed, this absorptive and regulatory proteome could serve as a starting point for using our models to study human nutrition.

#### 3.5.3 Suspended differentiated colonoids contain representative goblet cell populations

As thick mucus is a defining feature of the colonic epithelium, representative populations of the goblet cells responsible for its production [1] were desirable to recapitulate in our colonoids. Goblet cell localization was confirmed by staining for MUC2 (Fig. 5E, S1:9E), through which we observed, e.g., a mucus plume mid-extrusion (Fig. 5E). Using several such micrographs (n = 30 colonoids), the mean percentage of goblet cells per 4AD colonoid was determined to be 24 ± 16 %. This percentage aligned well with *in vivo* values found in the literature of ≈ 20% (Fig. 5F), especially considering the longitudinal gradient in mucus production found in the colon [1].

Though they all secrete MUC2, not all goblet cells are the same, with several specialized subpopulations identified to date [91,92]. We quantified proteins known to be enriched in intercrypt goblet cells, sentinel goblet cells, and noncanonical goblet cells (S1:9F-G), suggesting they may be represented. In short, markers of goblet cells (Fig. 5B, E, F, S1:9F-G) and other cell types representative of specific regions along the crypt-surface axis (see section 3.5.1) were found in both colonoid types. This result implies that we sufficiently recapitulated this axis, even in 4AD which lacked the geometry typical of crypts (see section 3.1.1).

#### 3.5.4 Tissue contains proteome-diluting cell types while isolated primary epithelial cells are essentially pure

Across all cell types, the fewest mucosal epithelial markers were detected in 4T (Fig. 5B, S1:9A). This hampered detection in tissue was apparent throughout this study (compare Fig. 6-7). Though excising the mucosa enriched the tissue sample for mucosal epithelium, proteomic analysis revealed the interfering presence of other cell types. Acting as dilutants, hematopoietic, endothelial, fibroblast, and mural cells appeared to still be abundant in 4T (Fig. 5G, 5H, S1:9A).

**Fig. 6:**
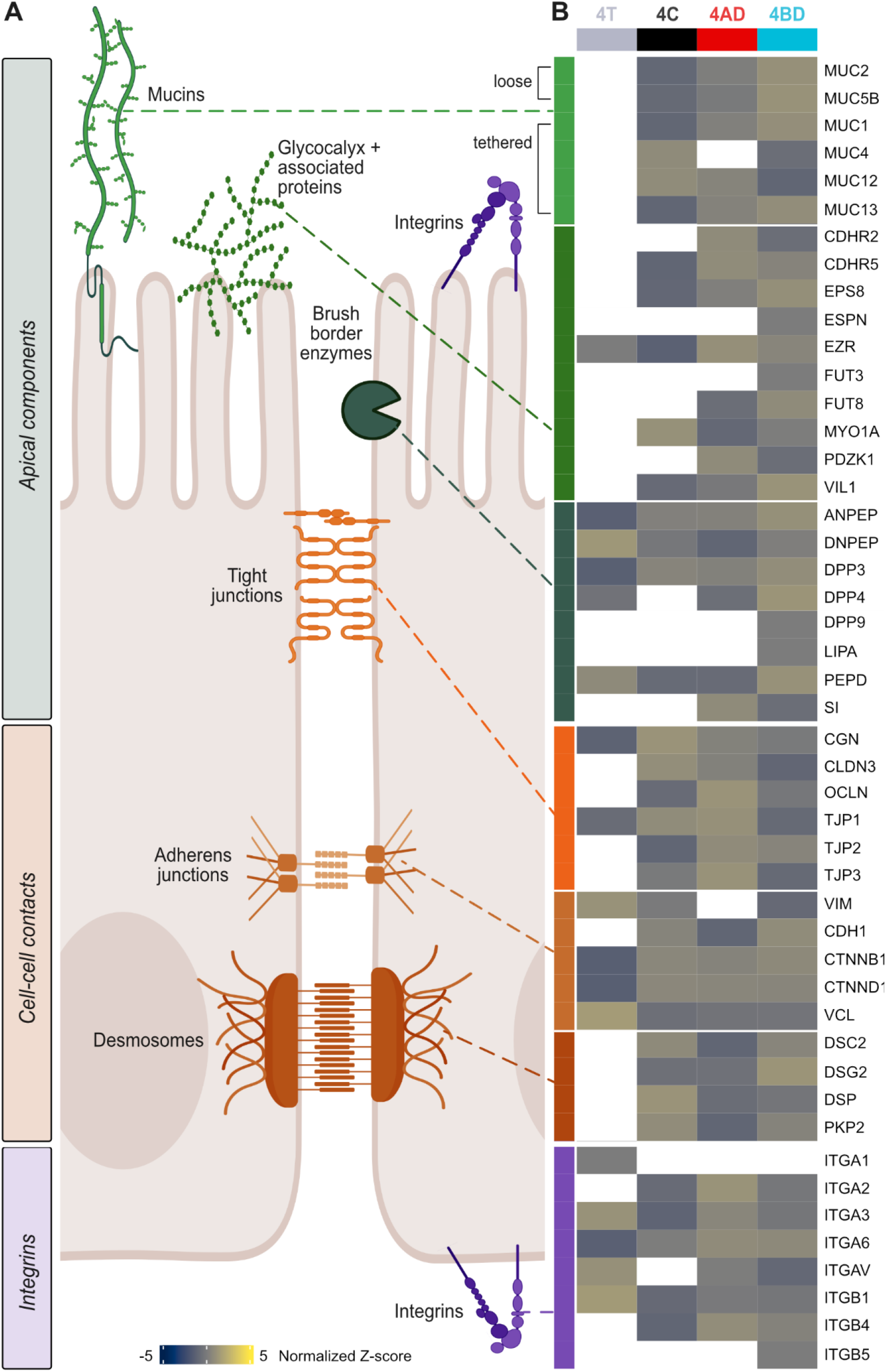
Suspended differentiated colonoids have native-like barrier proteomes. (A) Schematic representation of mucosal barriers. (B) Heatmap of protein expression of barrier markers based on global proteomics. Graphic in A produced in BioRender.

**Fig. 7:**
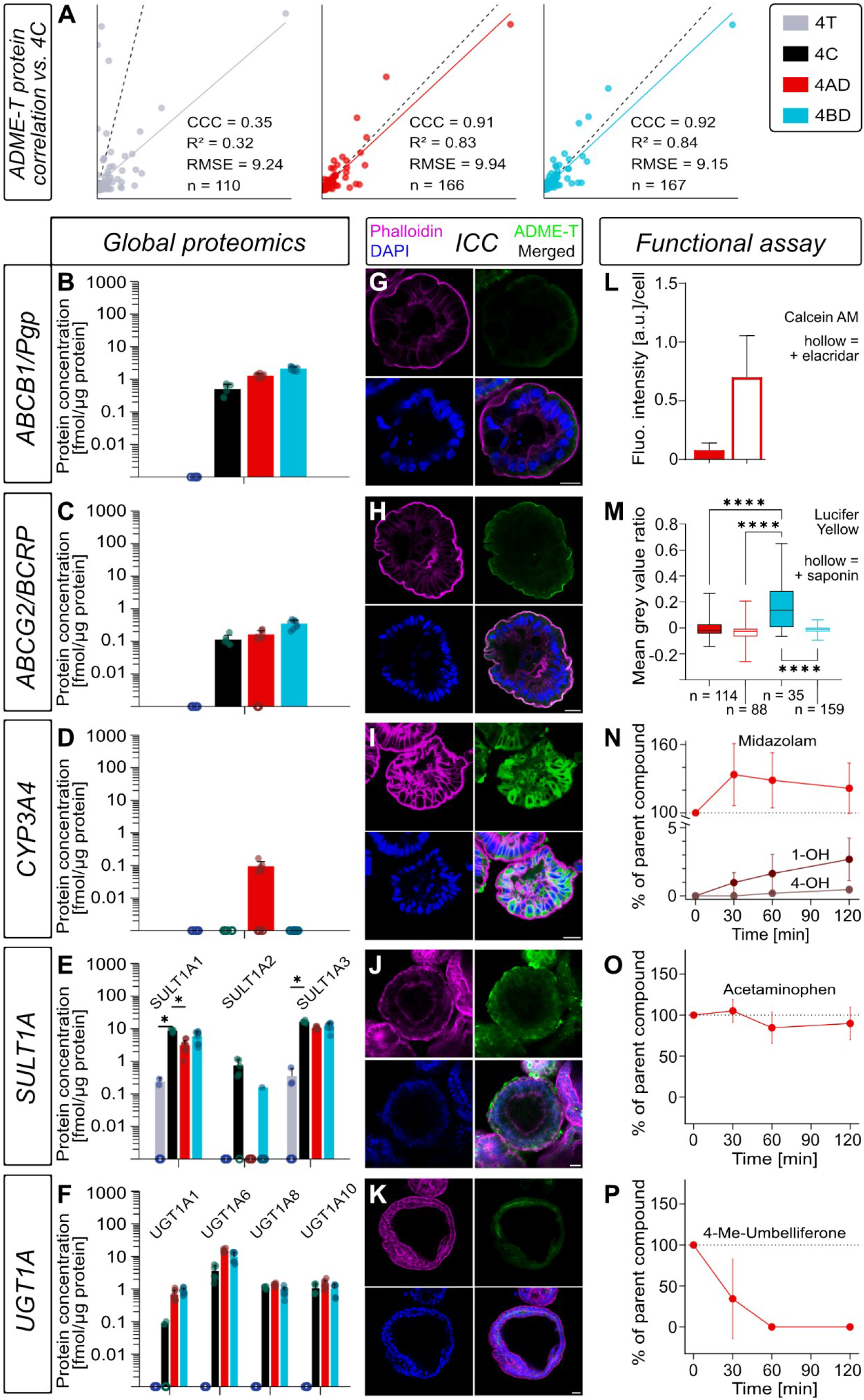
Suspended differentiated colonoids permit study of native ADME-T activity. A) Correlation of all ADME-T proteins relative to 4C, with the line of unity shown as a dashed line. CCC and R^2^ denote Lin’s concordance and Pearson’s correlation coefficients, respectively. (B-F) Protein expression (mean ± SD) in fmol/µg as measured by global proteomics. Hollow circles on the x axis indicate replicates where the protein of interest not quantifiable. (G-K) ICC images with the ADME-T protein of interest shown in the green fluorescent channel. (L-P) Functional ADME-T assays (mean ± SD). Refer to row headers for protein of interest. *p-value < 0.05 (DEqMS with Benjamini-Hochberg multiple testing correction), ****p-value < 0.0001 (Kruskal-Wallis test with Dunn’s post-hoc test and Holm’s multiple comparison correction).

Conversely, they were detected at reduced levels, if at all, in 4C (Fig. 5H, S1:9A), suggesting an epithelial isolate with high purity. Their purity positions isolated primary colonic epithelial cells as an improved proteomic reference, constituting a marked improvement over approaches relying on tissue or, more recently, non-purified isolated cells [22]. The colonoid samples were expected to be devoid of these contaminants but in fact contained some traces of fibroblast and mural proteins (though devoid of endothelial and blood marker proteins) (Fig. 5H, S1:9A). Their presence may be an artifact of, e.g., feeder layers used to generate the proprietary differentiation medium, whose exact composition is unknown.

### 3.6 Suspended differentiated colonoids have native-like barrier proteomes

#### 3.6.1 Suspended differentiated colonoids exhibit physiologically relevant barrier protein expression

In physiological epithelium, apical components (mucins, glycocalyx and associated proteins, brush border enzymes, and selected integrins), cell-cell contacts (tight junctions, adherens junctions, and desmosomes), and integrins contribute to its barrier function. Though many integrins act as a basolateral tether, either to the extracellular matrix or to other cells [35,36,93], these proteins can also localize to the apical membrane to regulate its barrier function [94] (Fig. 6A).

Key markers of each of these physical and chemical barriers were quantified across samples (Fig. 6B, S1:10A-G), but least so in 4T, for reasons discussed above (see section 3.5.4). In 4C, a cell suspension, junctional complex proteins were present but dysfunctional due to the compromised barrier. In both colonoid types, this barrier remained intact. Conversely, 4AD and 4BD were comparable to one another and to the 4C reference in most aspects.

However, differing differentiation status (as discussed in section 3.3.2) was also apparent at a barrier level. In 4AD, the outward-facing tight junctions and integrins were enriched (Fig. 6B), likely due to shear forces in the medium creating the need for increased tightness and rigidity. Presumably also due to shear forces, mucin abundance in 4AD was lower than in 4BD (Fig. 6B), with produced mucus likely being sloughed off and/or diluted by the surrounding medium. Conversely, levels of loose (or secreted) mucins and brush border enzymes were elevated in 4BD (Fig. 6B), where the apical side was shielded within the colonoid. Here, any secreted mucus was trapped within the inner cavity. Lacking access to key physiological mucus clearance processes such as peristalsis and microbial modulation [95], 4BD may have upregulated various peptidases (e.g., ANPEP, DPP3, DPP4, etc.) (Fig. 6B) to expedite mucus breakdown. Of these peptidases, PEPD (peptidase D) was most upregulated in 4BD relative to 4C (2.49 ± 0.53 fmol/µg vs. 0.81 ± 0.29 fmol/µg) (Fig. 6B, S1:10C). PEPD is involved in proline recycling [79], which is linked to ATP production [96]. Thus, its upregulation aligns with the high 4BD ATP turnover observed in Fig. 4J and a more proliferative and immature phenotype compared to 4AD (see section 3.3.2).

In line with this phenotype, 4BD also exhibited significantly lower expression levels of about half as much TJP1 (tight junction protein 1/zonulin-1) than 4C (0.22 fmol/µg vs. 0.53 fmol/µg) (Fig. 6B, S1:10D). In spite of this fact, 4BD colonoids were demonstrated to be tight, which is discussed in further detail below (see section 3.7.1). Moving further in a basal direction, adherens junction proteins were approximately comparable across 4C, 4AD, and 4C. Conversely, desmosome protein and integrin expression patterns were inverted for 4AD relative to 4C and 4BD (Fig. 6B, S1:10F), consistent with inverted apicobasal polarity. In summary, the barrier differences confirmed that polarity [29,35,36], curvature [29,77], and mechanical stimulation [97] impact differentiation.

#### 3.6.2 Suspended differentiated colonoids are equipped with endocytosis machinery

To round out the barrier profile, we also investigated endocytosis machinery, as it plays a role in the transcellular passage of large molecules and new modalities [98,99]. Coat and adaptor proteins (such as clathrins and adaptor-related protein complexes), scission and regulation proteins (such as dynamin), and proteins related to the cytoskeleton and downstream processes (such as actin) (S1:10H-I) were present in both 4AD and 4BD. Of note, most caveolins were not detected in the samples consisting of pure epithelium (S1:10I). The absence of epithelial caveolins mirrors both native tissue architecture, where expression is predominantly restricted to the stroma and smooth muscle [100], and reported baseline profiles of the colon carcinoma-derived Caco-2 epithelial cell line [99]. Consequently, endocytic transport within this pure epithelial layer is characteristically driven by caveolin-independent mechanisms [99].

As such, our colonoids appear suitable for future mechanistic characterization of non-small molecular drugs. Given the recent trend in oral and rectal delivery of such drugs [101], we anticipate that our colonoids will prove useful in their assessment. The colonoids’ low, native-like expression of, e.g., relevant peptidases (see section 3.6.1) implies their suitability for studying ADME-T relevant processes in this extended context.

### 3.7 Suspended differentiated colonoids permit study of native ADME-T activity

After deep proteomic and morphological characterization to establish fundamental workability of our models, we proceeded with their functional ADME-T characterization. Overall ADME-T protein [102] expression in 4AD and 4BD correlated well with the 4C reference (Lin’s concordance correlation coefficient = CCC = 0.91 and 0.92, respectively) (Fig. 7A, S1:3), highlighting their similarity to native epithelium. Conversely, 4T was once again highly divergent from 4C (CCC = 0.35) (Fig. 7A). This tissue divergence contextualizes and calls into question existing knowledge of the colonic ADME-T proteome derived from tissue (as highlighted in the introduction).

To remedy this gap and characterize the colonoid models’ core ADME-T functionality in depth, we utilized global proteomics (Fig. 7B–7F), providing additional validation via targeted proteomics (S1:3E, I, P), followed by ICC to visualize protein localization (Fig. 7G–7K). Finally, we performed functional ADME-T assays where we focused mainly on 4AD (Fig. 7L-P), having established that these colonoids had the most physiologically relevant proteome. Due to insufficient remaining primary cell material, it was unfortunately not possible to include 4C (or cells from any other donor) as a reference in targeted proteomic or metabolic characterization.

#### 3.7.1 High P-gp and BCRP efflux activity measured in suspended differentiated colonoids

Clinically relevant efflux transporters P-glycoprotein (ABCB1/P-gp) and Breast Cancer Resistance Protein (ABCG2/BCRP) were quantified at reference-like levels (0.50 ± 0.20 fmol/µg and 0.11 ± 0.04 fmol/µg, respectively, in 4C) in both 4AD and 4BD by global proteomics (Fig. 7B, 7C). These values were further validated by targeted proteomics, though higher values were measured for BCRP for 4AD (1.46 fmol/µg) and 4BD (1.77 fmol/µg) (S1:3E). As such, targeted or data-independent acquisition [103] approaches may be needed to accurately quantify this protein.

Both transporters were also shown to localize to the apical membrane (Fig. 7G, 7H, S1:3C). 4AD colonoids pre-treated with the P-gp inhibitor elacridar retained the P-gp substrate calcein AM intracellularly, hydrolyzing the acetomethoxy (AM) groups to generate a fluorescent signal [45]. In contrast, the untreated control was negligibly fluorescent, indicating continual efflux of calcein (Fig. 7L).

BCRP activity was assessed using the substrate LY [29]. As touched on above, accumulation of LY within the 4BD central cavity was observed (see section 3.4.1) despite a lesser maturation of tight junctions in this model (see section 3.6.1). Within this context, it became clear that the 4BD tight junctions were nonetheless sufficiently leak-proof and that apical efflux of LY was at play. Being a hydrophilic small molecule, LY has a low membrane permeability [29]. Still, especially in the presence of a concentration gradient, small quantities can reasonably be expected to enter cells. Comparing LY behavior in 4AD with that in 4BD, significant accumulation was only observed in non-permeabilized 4BD colonoids (Fig. 7M). In short, 4AD effluxed absorbed LY back into the surrounding medium while 4BD did so into the central cavity.

#### 3.7.2 Uptake transporter MCT1 localizes to the organoid exterior irrespective of configuration

Other transporters were quantified, but not further characterized, save for the uptake-mediating Monocarboxylate Transporter 1 (SLC16A1/MCT1). This transporter was highly expressed in 4C (12.40 ± 0.90 fmol/µg) and to a lesser extent in both colonoid types (2.22 ± 0.52 fmol/µg in 4AD and 3.96 ± 0.81 fmol/µg in 4BD) (S1:3G). Though the colonoid values were significantly different from 4C, they were still much higher than in 4T (0.66 ± 0.15 fmol/µg) (S1:3G). This 4T value is in the same order of magnitude as a reported median value of ≈ 0.14 fmol/µg from a targeted, tissue-based approach [19]. While this alignment is a testament to the quality of our global method, it also highlights how tissue-based proteomics approaches (see section 3.5.4) have underestimated the abundance of MCT1 in healthy colon.

Thus, uptake via MCT1 likely plays a greater role in the colon than previously published colon proteomes would suggest, in line with its biological function. Though its core functionality is the absorption of SCFAs [104], small xenobiotics resembling SCFAs are also transported (see section 3.8.2) [105]. As such, its interchangeable orientation [106] was of interest. MCT1 appeared to localize to the exposed side of the plasma membrane irrespective of colonoid type (i.e., apically in 4AD and basally in 4BD) (S1:3B), presumably driven by nutrient availability (i.e., access to the surrounding medium).

#### 3.7.3 Minimal phase I CYP metabolism in suspended differentiated colonoids

After transporter characterization, we investigated drug-metabolizing enzymes (DMEs). As a main perpetrator of phase I metabolism, cytochrome P450 (CYP) 3A4 was only quantified in 4AD, irrespective of the proteomic approach chosen (global, 0.095 ± 0.037 fmol/µg and targeted, 2.22 fmol/µg, respectively) (Fig. 1D). Failure to quantify this protein in 4C does not rule out its presence in this sample, but its level is unlikely to be high [107]. Recent studies on human jejunum, where CYP3A4 activity is known to play a greater role, suggest that primary cells and differentiated apical-out organoids yield approximately comparable CYP3A4 levels. Compared to these jejunal values (21.37 fmol/µg in primary cells) [29], we found negligibly low levels.

Interestingly, 4AD CYP3A4 expression seemed to be enriched in (a) certain cell type(s) (Fig. 7I), likely absorptive colonocytes due to their relative abundance and role in processing absorbed molecules. Such enrichment was less pronounced in 4BD, where we only detected CYP3A4 via ICC (S1:3H). Consistent with low CYP3A4 expression levels, no appreciable depletion of parent compound midazolam was observed in 4AD, and only traces of the primary (1-OH-midazolam) and secondary (4-OH-midazolam) metabolites were formed in functional assays (Fig. 7N). All in all, CYP3A4-mediated metabolism appears to be negligibly low (yet measurable) in the colonic epithelium.

#### 3.7.4 CES2 is highly expressed in suspended differentiated colonoids

Expression levels of additional phase I DMEs (alcohol dehydrogenases, aldehyde dehydrogenases, carboxylesterases, and other CYPs) are documented in S1:3J-M. Of these DMEs, most were expressed at comparable or lower levels relative to CYP3A4 in 4C and both colonoid types, implying a low impact on metabolism. Higher levels of 10 fmol/µg or greater were only reached by a small subset of enzymes, such as carboxylesterase 2 (CES2). CES2 was present at ≈ 19 fmol/µg in 4C and both colonoid types (S1:3K).

Though this expression is only about a third of that of the small intestine [29], it is still of clinical relevance. While the activation of esterified prodrugs can be exploited as a drug design feature, caution must be exercised where it can lead to toxic effects. An example of these toxic effects is the conversion of irinotecan to SN-38, primarily by CES2 [108], known to lead to local damage in the colonic epithelium [109]. With their native-like expression of this and other DMEs, the present colonoid models could be used in the preclinical assessment of such local toxic effects, improving patient safety and therapeutic outcomes.

#### 3.7.5 SULT1A exhibits low activity, but UGT1A-mediated phase II metabolism is rapid in suspended differentiated colonoids

Subsequently, we explored the sulfotransferase (SULT) 1A and UDP-glucuronosyltransferase (UGT) 1A protein families, both major effectors of phase II metabolism [110]. All SULT1A members save SULT1A4 were detected at low to moderate levels, though not in all samples (Fig. 7E). At 16.20 ± 1.29 fmol/µg in 4C, SULT1A3 was most highly expressed, with both colonoid cultures yielding comparable values. Staining revealed SULT1A localization predominantly to cytosolic vesicles in 4AD (Fig. 7J). Conversely, this protein family appeared to be distributed throughout the cell in 4BD (S1:3N).

Despite clear SULT1A presence and high permeability of the selected probe substrate acetaminophen [111], only scant metabolism was observed (Fig. 7O). Possible explanations are acetaminophen’s lack of SULT specificity as well as its reliance on microbial crosstalk for colonic phase II metabolite generation [112]. Additionally, though acetaminophen is known to exert toxic effects on the intestinal epithelium [113], it is unclear if this toxicity is observable over such short timescales. As SULT substrates with higher enzyme specificity emerge, it will be possible to resolve remaining issues around 3D-optimized SULT functional assays.

UGT1A family members also manifested at moderate levels (up to 14.90 ± 1.67 fmol/µg) in 4C, 4AD, and 4BD in global protein analysis (Fig. 7F), aligning with targeted values (S1:3P). UGT1A appeared to localize throughout the cell interior in 4AD (Fig. 7K), whereas the 4BD stain indicates enrichment within (a) specific cell type(s) (S1:3O). In 4AD, rapid metabolic transformation of the parent compound 4-methyl-umbelliferone was observed, resulting in its complete depletion within 30-60 min (Fig. 7P).

Several other phase II enzymes were quantified by global and targeted methods, though some only in trace amounts (S1:3P-Q). High expression and activity of phase II enzymes in the colon partially counteracts the microbially catalyzed deconjugation of phase II metabolites. These metabolites, formed in the liver and secreted into the bile, diminish enterohepatic recirculation and the bioavailability of molecules undergoing phase II metabolism [114]. Our colonoids could thus be suitable model systems to study the interplay of xenobiotic metabolism in liver, colon, and the microbiota.

### 3.8 Intracellular accumulation and bioavailability reflect active uptake, trapping, and epithelial geometry

#### 3.8.1 An 11-drug panel covers diverse molecular properties ADME-T functions

Having confirmed ADME-T suitability and functionality, we measured intracellular bioavailability (F_ic_), the product of total cellular accumulation (Kp) and the intracellular unbound fraction (f_u_). F_ic_ estimates intracellular unbound drug exposure relative to the extracellular concentration and thereby drug availability to intracellular drug targets. Thus, F_ic_ is a measure of the actual drug potency at target [38] and off target sites [115]. Alongside 4AD and 4BD, we included HEK293 cells, an established model for F_ic_ measurements [38,40,43,116,117], as a methodological comparator. Since intracellular binding depends on intracellular lipid content and composition [39,43], we determined f_u_ separately for each model.

The panel of 11 small molecule drugs selected here covered diverse molecular properties, particularly lipophilicity (S1:4B), within the core of chemical drug space of approved drugs (Fig. 8A, S1:4D-G). Collectively, the drugs were reported substrates of all efflux transporters and phase I and phase II DMEs explored in section 3.7. Most of the compounds in the panel were reported substrates of P-gp and/or of BCRP, and of CYP3A4 (S1:4H), consistent with the broad involvement of these ADME-T proteins in small-molecule drug disposition [118]. Following individual functional assays in section 3.7, we used cassette dosing [38] for characterizing drug accumulation and intracellular bioavailability. Where available, we compared our findings to colonic absorption data (sections 3.8.2-3.9.3).

**Fig. 8:**
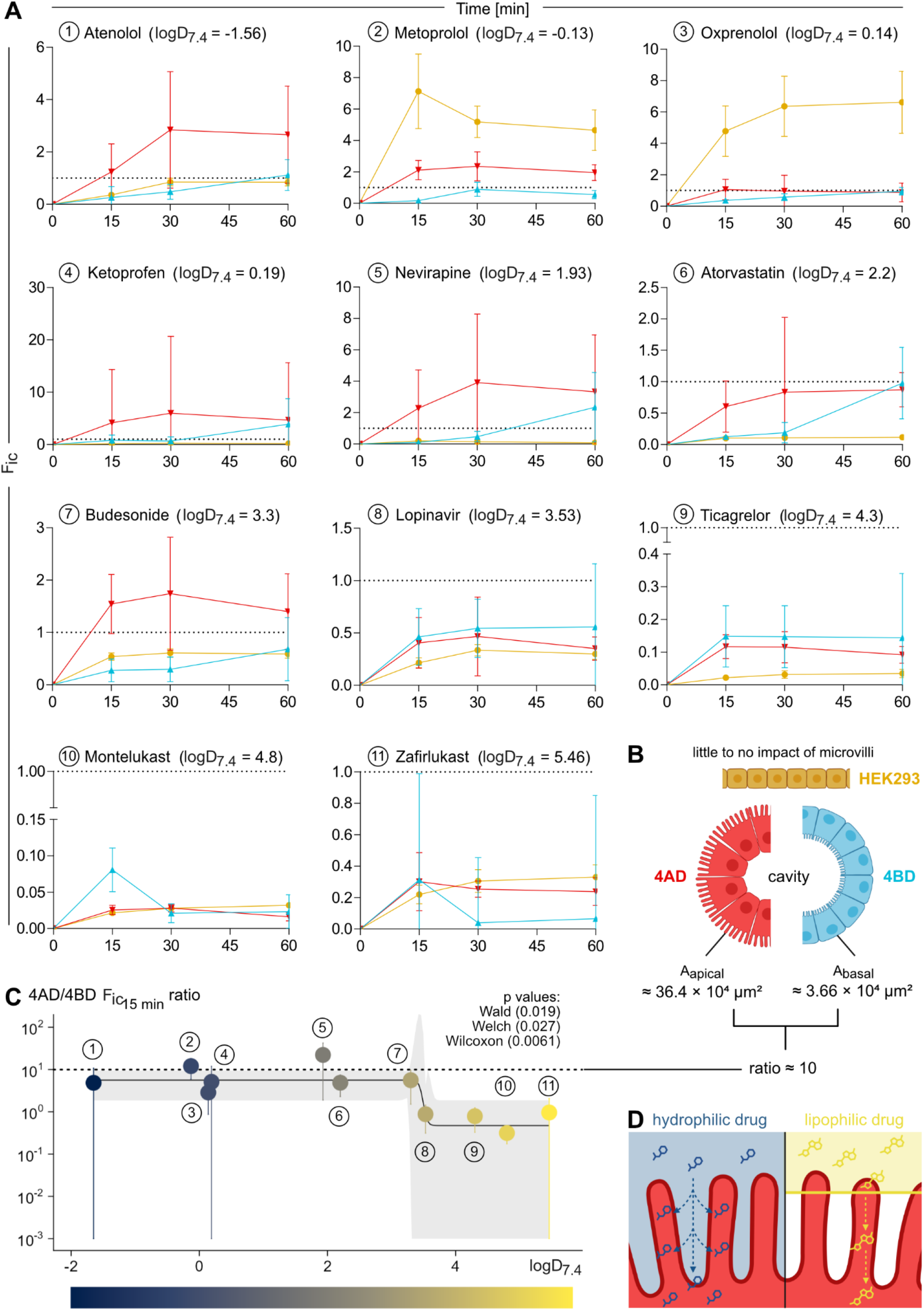
Intracellular accumulation is driven by active uptake, trapping, and model geometry. (A) F_ic_ over time (0-60 min, geometric mean ± SEM) for the studied 11-drug panel. Refer to panel B for color legend. (B) Schematic depiction of the sample types studied (HEK293, 4AD, and 4BD) and the impact of microvillus presence and orientation. (C) 4AD to 4BD F_ic_ ratio at 15 min (geometric mean ± SEM) by compound, colored by relative lipophilicity. Refer to panel A for compound numbering. Dotted line at 9.94 corresponds to ratio indicated in panel B, 95% confidence interval for curve fit shown as a grey ribbon. (D) Schematic of presumable lipophilicity-dependent use of the available microvillus surface. Graphics in B and D produced in BioRender.

#### 3.8.2 Compound-specific accumulation and bioavailability suggest contributions from active uptake and trapping

In all three models (4AD, 4BD and HEK), F_ic_ values were generally lower for the more lipophilic compounds, Fig. 8A). The accompanying reduction in f_u_ values is consistent with the established relationship between lipophilicity and cellular lipid binding [38,39,43,119], examined further in section 3.9. When individual compounds were compared between models, colonoids generally showed higher or comparable F_ic_ values relative those in HEK293 (Fig. 8A), reflecting greater total accumulation (Kp) (S1:6B-C).

Metoprolol and oxprenolol were exceptions, with approximately 2- to 8-fold and 7- to 8-fold higher F_ic_ values, respectively, in HEK293 cells than in the colonoids (Fig. 8A, S1:6C). These beta blockers, being weak bases, are susceptible to lysosomal trapping [120]. However, oxprenolol reached comparable Kp values across all samples (S1:6B-C), indicating that its higher HEK293 F_ic_ (Fig. 8A, S1:6C) reflected a higher f_u_ rather than greater total accumulation (see section 3.9.1). Comparable Kp values do not, however, exclude differences in the process contributing to accumulation.

Ketoprofen reached the highest average F_ic_ values (≈ 5) in 4AD (Fig. 8A). These high F_ic_ values align with ketoprofen’s high P_eff_ value (3.37 × 10^-4^ cm/s) [3] (S1:4C) and F_ic_ > 1 indicates the involvement of active uptake. Although ketoprofen is a substrate of organic anion transporters 1 to 3 (OAT1, OAT2, and OAT3) (S1:4H), none of these OATs were quantified in our dataset (S1:3G). In contrast, MCT1, which also transports ketoprofen [121,122] (S1:4H), was relatively abundant and accessible at the exterior of both our colonoids (see section 3.7.2) and could therefore contribute to the accumulation. The slower convergence to ketoprofen equilibrium in 4BD vs. 4AD (Fig. 8A) despite comparable, outward-facing levels of MCT1 (see section 3.7.2) suggests additional drivers of compound enrichment, including epithelial geometry (see section 3.8.3). Kp values were most variable in 4BD particularly for, e.g., zafirlukast (S1:6B-C). Mucus associated (see section 3.6.1) retention in the closed internal lumen may possibly contribute to this variability, consistent with suggestions that aggregated glycans can act as drug binding sinks [43].

#### 3.8.3 Epithelial geometry is associated with differences in accumulation kinetics

Consistent with lipid-driven accumulation [39], Kp generally increased in relation to logD_7.4_ across all three models. HEK293 monolayers had markedly lower Kp than colonoids for most compounds, with oxprenolol and zafirlukast as exceptions (S1:6B-C). These differences may reflect both cellular composition and culture geometry. Both colonoid configurations enclose a cavity (Fig. 8B), which is a lumen in the classical sense only in 4BD. Still, in both configurations, the presence of a cavity introduces an additional compartment that may influence accumulation and equilibration. Overall, 4AD and 4BD reached comparable K_p_ (S1:6B-C) and F_ic_ values (Fig. 8A, S1:6C) for many compounds, although equilibrium may not have been reached within 60 min for some compounds in 4BD, including atenolol and budesonide.

Microvilli, a defining feature of colonic epithelium that is essentially absent in kidney-derived HEK293 cells [123] (Fig 8B), provide further geometric distinction. Compounds with a logD_7.4_ ≤ 3.3 accumulated more rapidly in 4AD, where microvilli face the incubation medium, than in 4BD, where the exposed surface area is basolateral and hence comparatively flat (Fig. 8A-B). At 15 min, i.e., prior to steady-state, the 4AD/4BD F_ic_ ratios for these less lipophilic compounds were broadly consistent with the approximately 10-fold difference in exposed membrane area [124] (Fig. 8B-C, S1:6D-J). However, this ratio incorporates differences in both accumulation and f_u_ and therefore does not directly measure relative uptake rates. In contrast, compounds with logD_7.4_ > 3.3 generally accumulated rapidly in both configurations, with little apparent benefit from the enlarged microvillar surface (Fig. 8A, C, S1:6B-C).

These findings are consistent with differential use of the microvilli surface: less lipophilic compounds seem to diffuse between the microvilli and access a larger membrane surface area, whereas rapid partitioning of more lipophilic compounds into the first available membrane may favor entry near microvillus tips (Fig. 8D). This interpretation extends the mechanism previously described for small intestinal villi [125] and applied in PBPK modeling [12] to the microvillar scale.

### 3.9 Lipidomic analysis identifies potential contributors to intracellular binding

To investigate the binding component of F_ic_, we measured f_u_ by equilibrium dialysis of homogenates from 4AD, 4BD, and HEK293 cells and characterized the same matrices by targeted lipidomics. Phospholipids are established major determinants of cellular binding, whereas the contribution of neutral lipids is more compound-dependent [39,43]. Lysosomal trapping, which depends on intact intracellular pH gradients, is distinct from binding measured in homogenates [39].

The close molecular and functional correspondence between colonoids and isolated colonic epithelial cells, particularly for 4AD, supported the use of these models to investigate epithelial drug binding. Their lipidomes were not, however, directly benchmarked against native primary epithelial cells.

#### 3.9.1 Phospholipid differences are associated with increased binding to colonoids

HEK293 cells had approximately an order of magnitude less choline-containing phospholipid per cell than colonoids (Fig. 9A) and up to more than 20-fold higher f_u_ values (Fig. 9B, S1:6C). This association is consistent with previous findings linking cellular phospholipid content to drug binding [36,37,41]. In our 11-drug panel, f_u_ decreased markedly around logD_7.4_ of approximately 2, with the clearest HEK293-colonoid differences among the less lipophilic compounds (Fig. 9B).

**Fig. 9:**
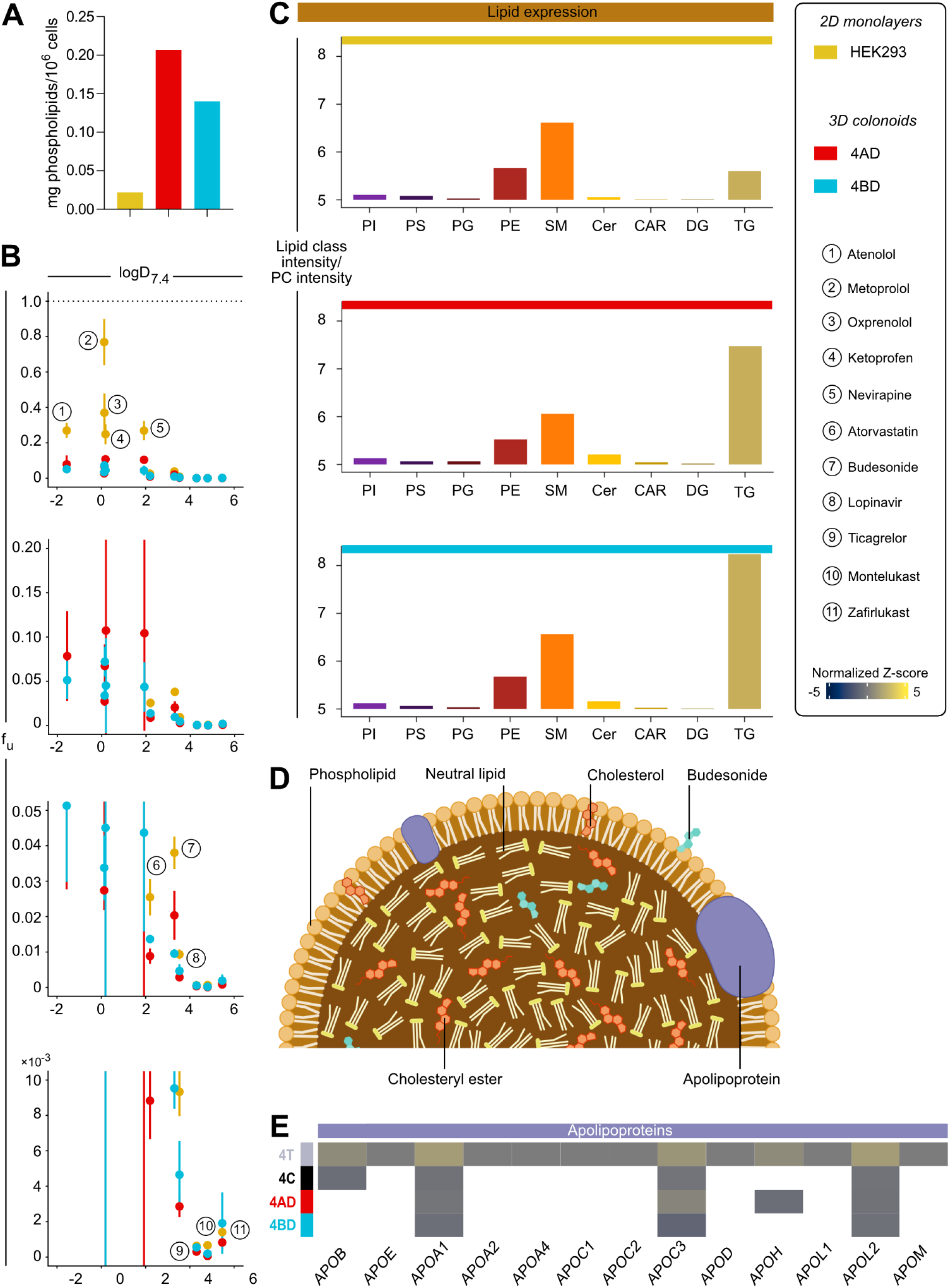
Lipidomic analysis reveals lipid classes driving intracellular binding. (A) Mean total choline-containing phospholipid content in each sample. (B) Cellular f_u_ values (geometric mean ± SEM) shown at different y axis scalings to highlight logD_7.4_-dependent differences. (C) Mean lipid expression of phospholipids (phosphatidylinositol = PI, phosphatidylserine = PS, phosphatidylglycerol = PG, phosphatidylethanolamine = PE, phosphatidylcholine = PC, and sphingomyelins = SM) and neutral lipids (ceramides = Cer, acetylcarnitines = CAR, diglycerides = DG, triglycerides = TG). (D) Schematic of the structure of a chylomicron, with budesonide added. Graphic produced in BioRender. (E) Heatmap of apolipoprotein expression based on global proteomics.

The enzymatic assay indicated more phospholipid in 4AD than in 4BD (Fig. 9A), whereas binding of less lipophilic compounds was generally stronger in 4BD (Fig. 9B). Targeted lipidomics also gave generally higher phospholipid intensities in 4BD (S1:7B). These measurements differ in analytical scope and normalization and therefore do not establish a single ranking of total phospholipid abundance between the two colonoid configurations. Possible contributors include matrix interference, extraction efficiency, and normalization to cellular protein. Although colonoid protein values were corrected for extracellular matrix proteins and mucins, quantifying intracellular protein remains challenging because extracellular proteins are also recovered. The assumption of comparable protein per cell volume intracellularly [126] does not remove this potential contribution.

Phospholipid composition also differed. Relative to PC, both colonoid configurations were enriched in phosphatidylinositol (PI) and phosphatidylglycerol (PG) compared to HEK293 (Fig. 9C). Together, the binding and lipid measurements support determining f_u_ directly in colonoids rather than substituting HEK293-derived values.

Stronger intracellular binding should not be equated with poor epithelial permeability. For example, metoprolol has an effective permeability (P_eff_) value of 1 × 10^-4^ cm/s and relative bioavailability of 100% after colonic administration (S1:4C) [3,127]. The corresponding relative bioavailability of oxprenolol is 82% (S1:4C) [3].

#### 3.9.2 Neutral lipid differences may contribute to binding of highly lipophilic compounds

A contribution from neutral lipids has been suggested for highly lipophilic compounds, particularly above logD_7.4_ ≈ 3 [39]. In this range, HEK293 generally had higher f_u_ values than colonoids (Fig. 9B), alongside lower relative triglyceride signals (Fig. 9C). Relative to PC, triglycerides were most abundant in 4BD, whereas 4AD showed greater ceramide enrichment (Fig. 9C). These compositional differences may contribute to the binding patterns, but the present comparisons do not resolve the contribution of individual lipid classes. Cholesterol, which is enriched in apical membranes [168], may also contribute, but was not quantified.

The low reported colonic P_eff_ of ticagrelor, 0.021 × 10^−4^ cm/s, and its relative bioavailability of 18% after colonic administration [3] (S1:4C) illustrate that high lipophilicity (logD_7.4_ = 4.3) does not ensure efficient colonic absorption. These clinical findings do not, however, establish neutral lipid binding as the limiting mechanism.

#### 3.9.3 Budesonide retains high intracellular bioavailability despite high affinity for lipid sinks

Budesonide had f_u_ values of approximately 2% in 4AD and 1% in 4BD, relatively high compared with neighboring lipophilic compounds in the panel (Fig. 9B, S1:6C). Nevertheless, these values indicate extensive intracellular binding. Its high accumulation, reaching an apparent steady state within 15 min in 4AD, yielded F_ic_ > 1 (Fig. 8A, S1:6B-C).

As a corticosteroid derivative (S1:4A), budesonide shares structural features with endogenous steroids. Proteins associated with lipid assembly and transport, including chylomicron-related proteins, were enriched in colonoids (Fig. 9D-E, S1:7C-E). Although cholesterol is an abundant cellular lipid [128,129] neither its abundance nor its turnover were measured here. Chylomicron production, typically associated with the small intestine [129], has been reported in colon-derived cells [130], and is therefore a possible contributor to lipid-associated drug retention [1].

Culture conditions may also influence this phenotype. OGM and ODM media formulations, although intended for colonic cultures [131], support the greater nutritional requirements in the small intestine [29,131]. They also derive from the original culture approaches established for the small intestine [13,131]. Nutrient exposure without physiological modulation by the small intestine and microbiota may contribute to the expression of some small intestinal features in colonoids, including SGLT1 (section 3.5.2) and sucrase isomaltase [88,89,132] (Fig. 6, S1:10D).

Retention in lipid aggregates could contribute to budesonide accumulation (Fig. 9D). If such aggregates are disrupted during homogenization, measured f_u_ may differ from that in intact cells. Ultrasonication can remove chylomicron-associated apolipoproteins [133] and nanoemulsify lipids [134], but whether this occurred here, and how it affected budesonide binding, remains unresolved.

The reported relative systemic bioavailability of 53% following colonic budesonide administration [3] is compatible within complete systemic delivery but does not identify a specific retention mechanism. In this study, recovery of unchanged budesonide remained within the accepted mass balance limits of 75–125%, indicating no substantial loss of parent compound under assay conditions. CYP3A4, its major metabolic pathway (S1:4H), showed low activity in colonoids (section 3.7.3), suggesting a limited contribution from metabolism. Local esterification, previously described for budesonide in the lung [135], cannot be excluded but was not associated with a detectable loss of unchanged budesonide beyond these limits.

Thus, substantial total accumulation sustained intracellular unbound budesonide exposure despite extensive binding. This finding is consistent with its established local efficacy after colonic delivery [5,6] and illustrates the value of combining accumulation and binding measurements in a physiologically relevant colonic epithelial model [5,6].

## 4 Conclusion

In conclusion, suspended, suitably differentiated 3D colonoids reproduce the structural, molecular, and functional signatures of native colonic epithelium. Donor-matched, non-enzymatically isolated epithelial cells provide a more representative native reference proteome by reducing the contribution of non-epithelial cell types present in bulk tissue. Overall, apical-out differentiated colonoids have the most native-like signatures and are most amenable to 3D-adapted assays. These models demonstrate native-like ADME-T activity mediated by clinically relevant transporters and drug-metabolizing enzymes. Studies of drug distribution and intracellular bioavailability (F_ic_), reflecting drug availability to intracellular targets, revealed how 3D epithelial organization and microvillar architecture shape absorption in a lipophilicity-dependent manner. Complementary binding and lipid analyses showed higher phospholipid content and generally stronger intracellular drug binding in colonoids than in HEK293 cells, supporting the use of colonoid-specific binding measurements to assess colonic intracellular drug exposure.

Together, these findings establish suspended differentiated 3D colonoids and F_ic_ measurements as tools for elucidating colonic drug behavior. Their use as physiologically relevant NAMs can support mechanistic studies of colonic drug exposure and inform next generation risk assessment (NGRA).

## Acknowledgements

The authors acknowledge the facilities and technical assistance of the Umeå Center for Electron Microscopy (UCEM) at the Chemical Biological Center (KBC), Umeå University, a part of the National Microscopy Infrastructure NMI (VR-RFI 2019-00217), as well as the Swedish Metabolomics Center located at the same premises. We thank Rasmus Hammar for data analysis support.

## Contributors

Rebekkah Hammar: Conceptualization, methodology, validation, formal analysis, writing - original draft preparation, investigation, visualization, validation, project administration; Patrik Lundquist: Methodology, supervision, funding acquisition; Dinh Son Vo.: Methodology, investigation, validation, Foteini Tzioufa.: Methodology, formal analysis; Jens Eriksson: Methodology; Ana C.C. Lopes.: Resources, investigation; Helen S. Hammer.: Investigation, formal analysis; Oliver Pötz: Methodology, Funding acquisition; Maria Letizia di Martino: Methodology; Wilhelm Graf: Resources, project administration; Madlen Hubert: Formal analysis, funding acquisition, visualization; Magnus Sundbom.: Resources, project administration; Mikael E. Sellin.: Resources, supervision, funding acquisition, project administration; Per Artursson.: Conceptualization, supervision, funding acquisition, project administration. All authors: Writing – review and editing.

