## Supplementary Information 1 for "Native-like 3D colonoids reveal mechanisms of colonic drug disposition"

### S1:1 Generation of matched colonic samples

#### Medical and demographic information for colon biopsy donors

**A**

| Donor ID | Sex | Age group | Colon segment | Material used for proteomics |
|---|---|---|---|---|
| Donor 1 | F | Seniors | Sigmoideum | Tissue |
| Donor 2 | M | Old-aged adults | Sigmoideum | Tissue, cells, colonoids |
| Donor 3 | M | Old-aged adults | Transversum | Tissue |
| Donor 4 | M | Old-aged adults | Transversum | Tissue, cells, colonoids |
| Donor 5 | M | Old-aged adults | Transversum | Tissue |
| Donor 6 | F | Old-aged adults | Transversum | Tissue |
| Donor 7 | M | Old-aged adults | Transversum | Tissue, colonoids |
| Donor 8 | F | Seniors | Sigmoideum | Tissue |
| Donor 9 | M | Middle-aged adults | Ascendens | Tissue, cells |
| Donor 10 | M | Middle-aged adults | Sigmoideum | Tissue, cells |
| Donor 11 | M | Seniors | Sigmoideum | Tissue |

**S1:1.** (A) Age groups: middle-aged adults (31-45), old-aged adults (46-75), seniors (>75). F = female, M = male.

#### Primary cell isolations

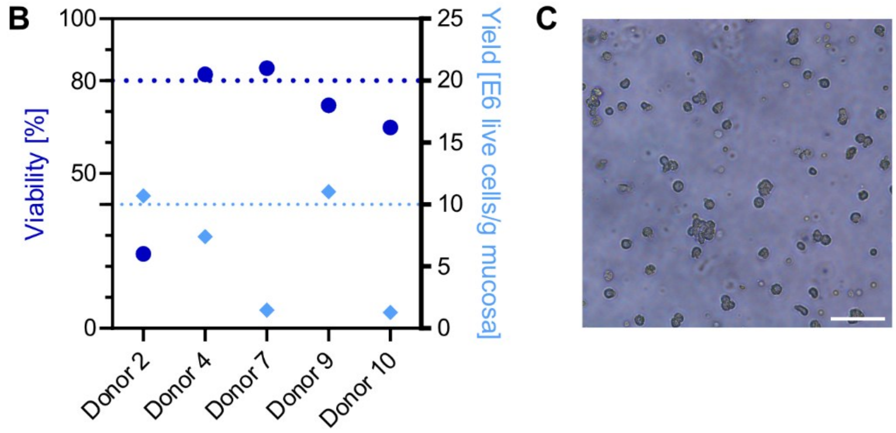

**S1:1 (continued).** (B) Primary cell viabilities and yields by donor. Cells from donor 7 were used for apoptosis assay. (C) Brightfield micrograph of primary cells post-isolation (scale bar 100 µm).

#### Colonoid establishment

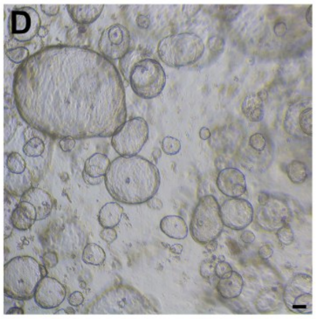

**S1:1 (continued).** (D) Growth-medium maintained colonoids at passage 8 in Matrigel at day 6 post-passaging (scale bar 100 µm).

### S1:2 Morphology and viability

#### Antibodies used for immunocytochemistry staining

**A**

| Antibody | Source | Identifier |
|---|---|---|
| Invitrogen™ Alexa Fluor™ 660 Phalloidin | Thermo Fisher Scientific | 10402322 |
| Invitrogen™ ZO-1 Monoclonal Antibody (ZO1-1A12), Alexa Fluor™ 555 | Thermo Fisher Scientific | MA3-39100-A555 |
| Invitrogen™ P-Glycoprotein Monoclonal Antibody (C219) | Thermo Fisher Scientific | MA1-26528 |
| Invitrogen™ SULT1A3 Polyclonal Antibody | Thermo Fisher Scientific | PA5-97760 |
| Anti-UGT antibody [7H2AF11] | Abcam | ab129729 |
| Invitrogen™ CYP3A4 Monoclonal Antibody (3H8) | Thermo Fisher Scientific | MA5-17064 |
| Anti-MUC2 antibody | Abcam | ab90007 |
| Anti-BCRP/ABCG2 antibody [EPR21122] | Abcam | ab229193 |
| Invitrogen™ FABP1 Recombinant Rabbit Monoclonal Antibody (ARC0545) | Thermo Fisher Scientific | MA5-35080 |
| Anti-MCT1/Monocarboxylic acid transporter 1 antibody | Abcam | ab85021 |
| Goat Anti-Mouse IgG H&L (Alexa Fluor™ 488) | Abcam | ab150113 |
| Invitrogen™ Goat anti-Rabbit IgG (H+L) Cross-Adsorbed Secondary Antibody, Alexa Fluor™ 488 | Thermo Fisher Scientific | A-11008 |

**S1:2.** (A) Tabulated sourcing information for antibodies used in ICC staining.

#### Viability

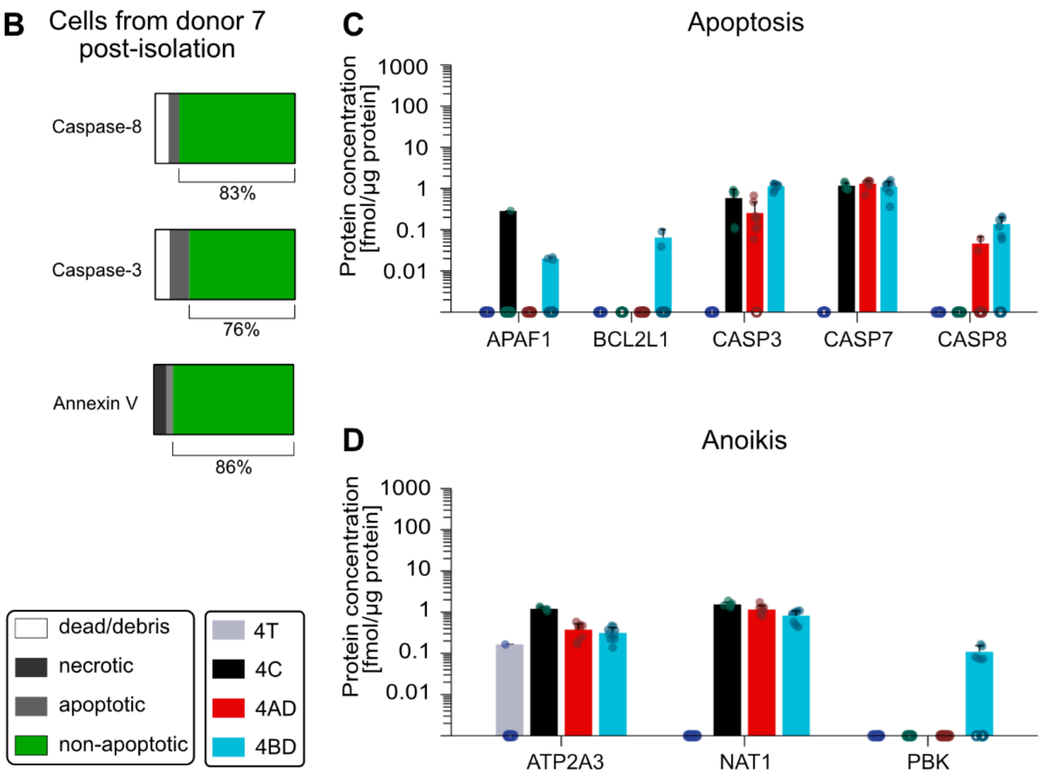

**S1:2 (continued).** (B) Proportion of cells expressing cell death markers in cells from donor 7 (Hu22-7col). (C) and (D) protein expression levels of cell death markers (apoptosis and anoikis, respectively), based on global proteomics. Hollow circles on the x axis indicate replicates where the protein of interest was not quantifiable.

### S1:3 Additional ADME information

#### Transporters

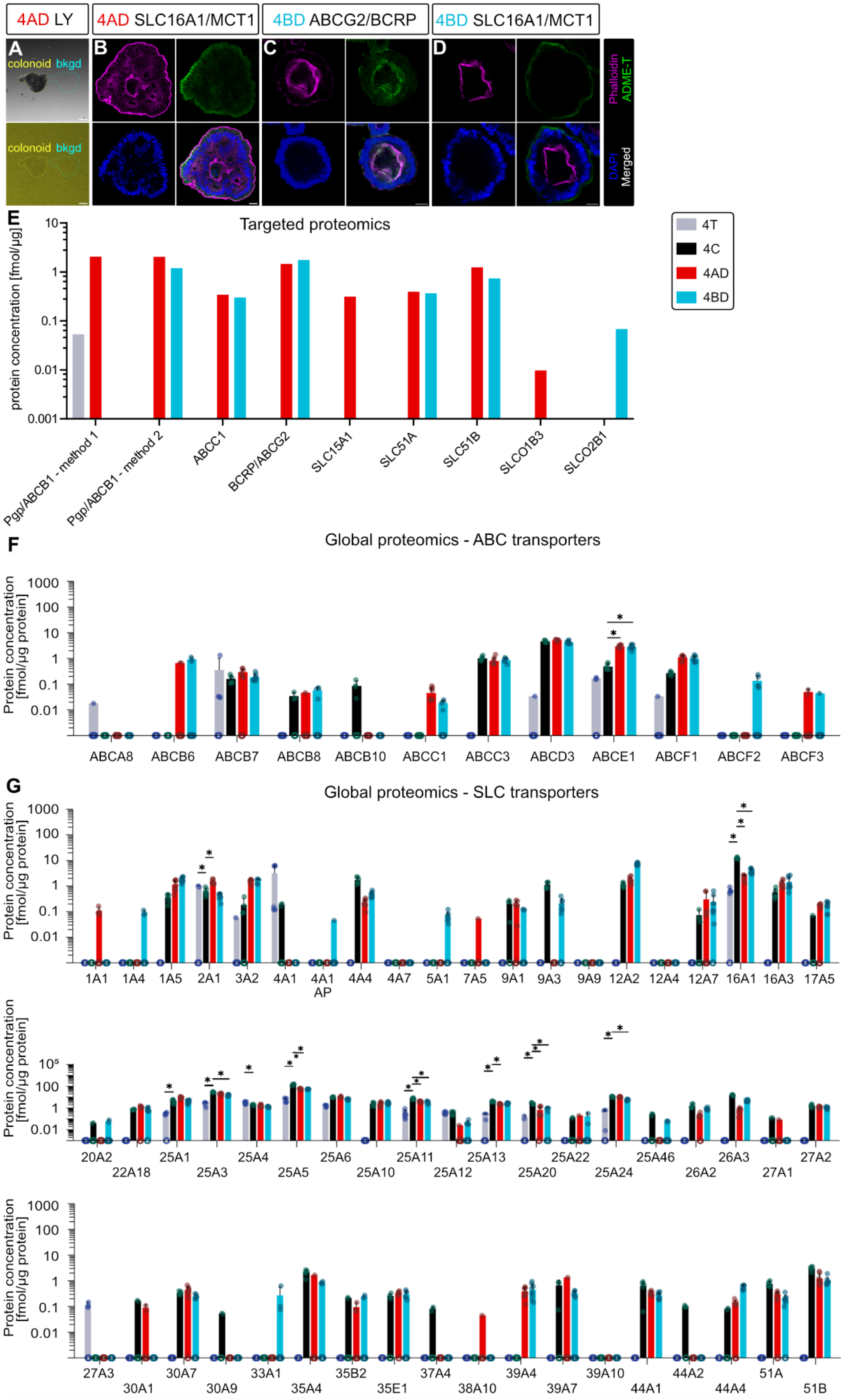

**S1:3 (continued from previous page)**. (A), Example micrograph of colonoid and matched background during live-cell LY assay. (B) ICC image of MCT1 in 4AD. (C) and (D) ICC images of 4BD, showing BCRP and MCT1, respectively. (E) Targeted proteomics of transporters. (F) Global proteomics of additional ABC transporters. (G) Global proteomics of SLC transporters. For global proteomics data, hollow circles on the x axis indicate replicates where the protein of interest was not quantifiable. *p-value < 0.05 (DEqMS with Benjamini-Hochberg multiple testing correction).

In Fig. 7, the flurorescence intensity ratio shown is that of

$$\frac{(Mean\ intensity\ in\ colonoid - Mean\ intensity\ in\ background)}{Mean\ intensity\ in\ background}$$

### Phase I metabolism

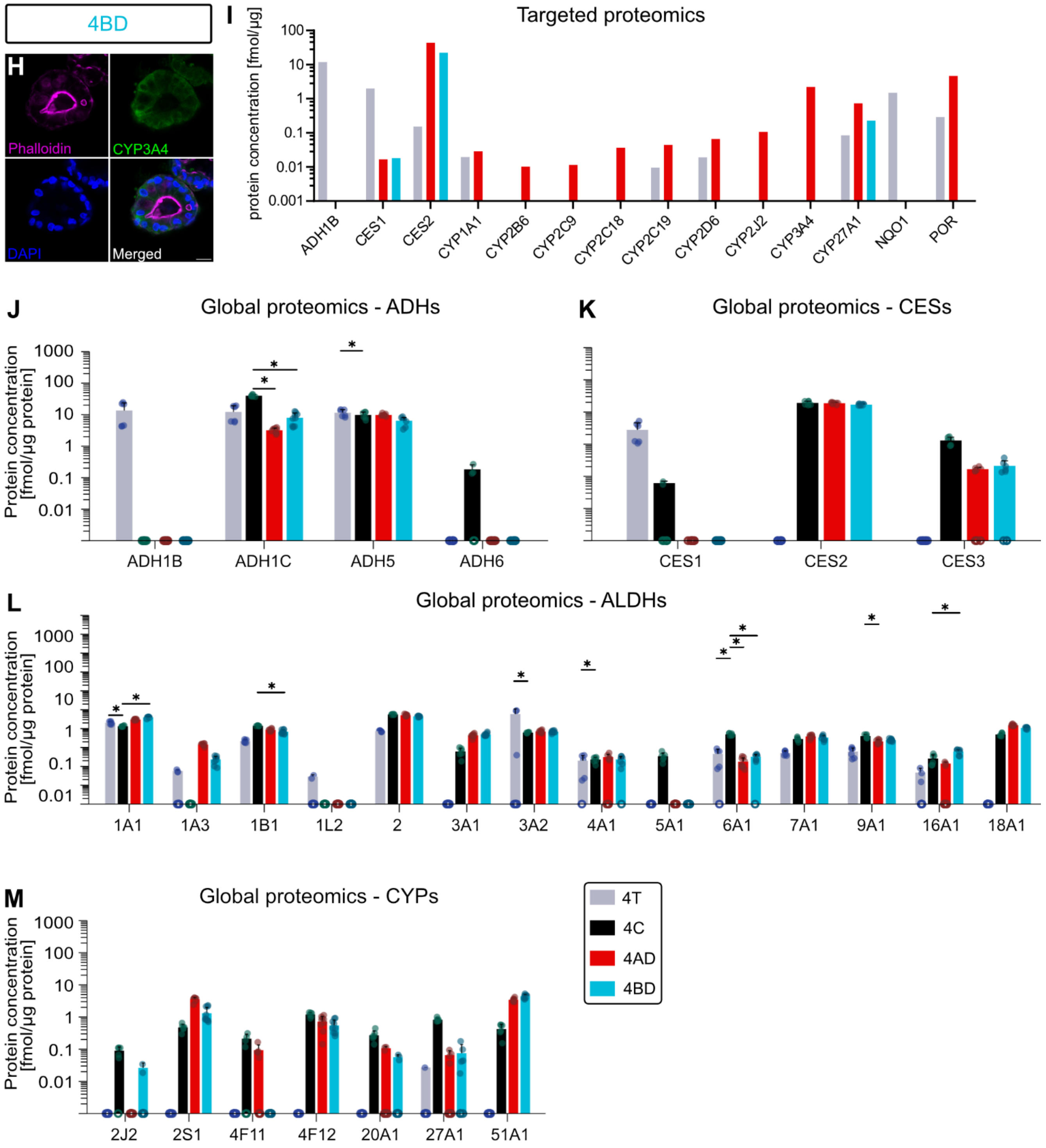

**S1:3 (continued).** (H) ICC image of CYP3A4 in 4BD. (I) Targeted proteomics of phase I DMEs. (J-M) Global proteomics of phase I DMEs. For global proteomics data, hollow circles on the x axis indicate replicates where the protein of interest was not quantifiable.

#### Phase II metabolism

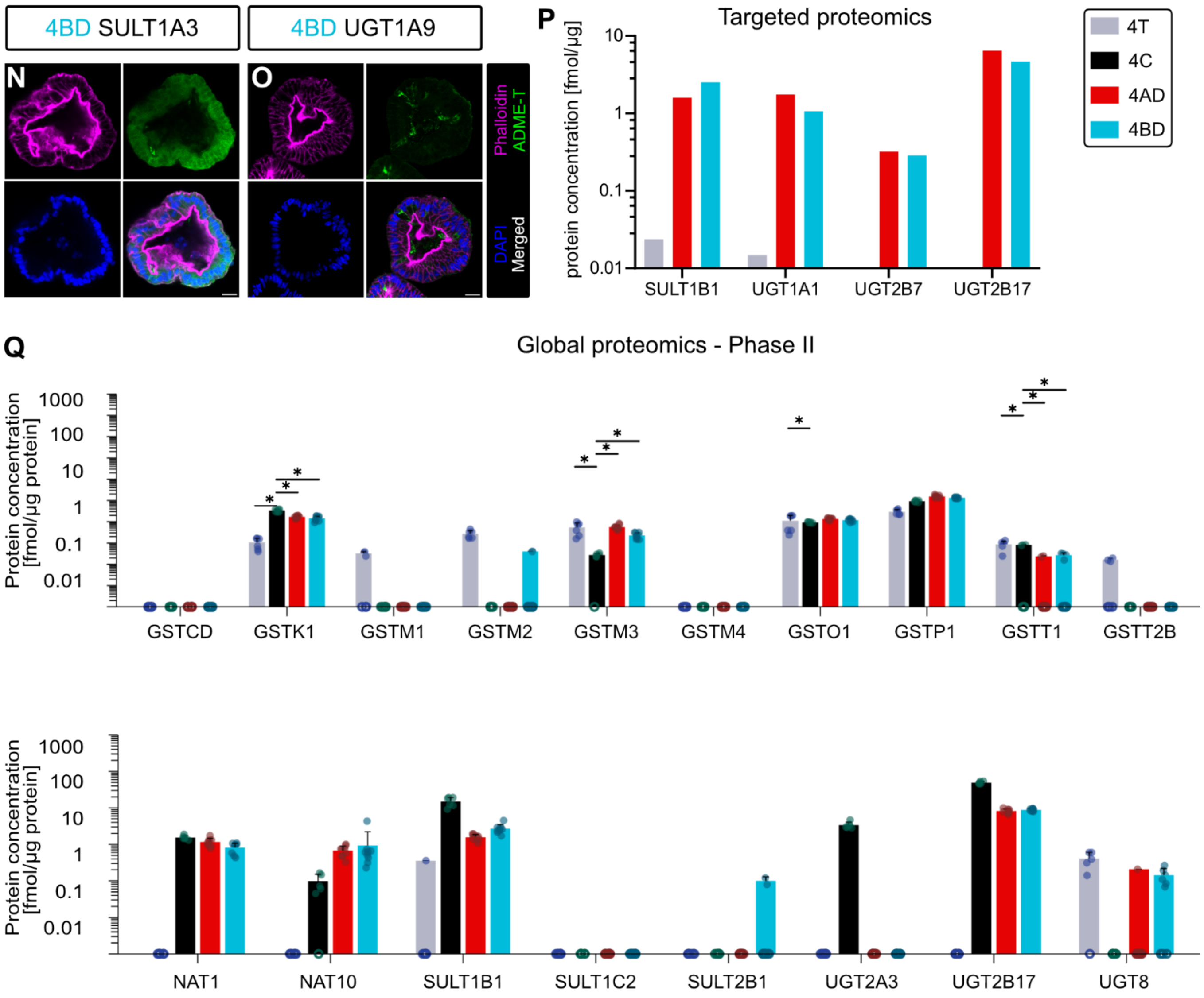

**S1:3 (continued).** (N) and (O), ICC images of SULT1A3 and UGT1A9, respectively, in 4BD. (P) Targeted proteomics of phase II DMEs. (Q) Global proteomics of phase II DMEs. For global proteomics data, hollow circles on the x axis indicate replicates where the protein of interest was not quantifiable.

### S1:4 Drug panel information

#### Compound structures

**A**

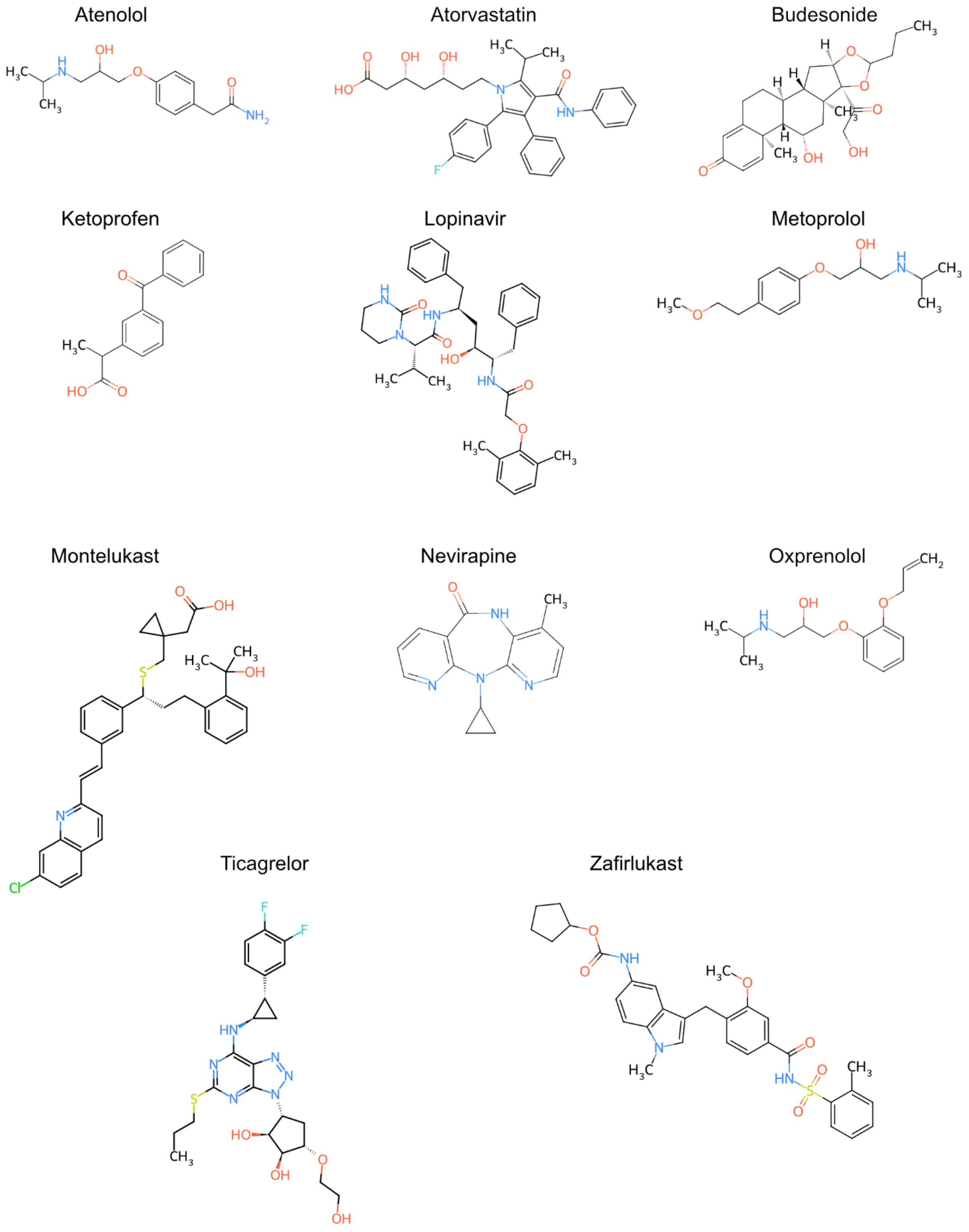

**S1:4.** (A) Compound structures of the 11-drug panel used in this study.

#### Physicochemical and pharmacokinetic properties

**B**

| Compound | Mw | PC | TPSA | $logD_{7.4}$ | $pK_a$ | CL | Vd | B:P | $f_{u,plasma}$ | FPE | BCS |
|---|---|---|---|---|---|---|---|---|---|---|---|
| Atenolol | 266.3 | 1 | 84.6 | -1.56 | 9.21[b] | 10.5 | 18 | 1.07 | 0.91 | 0 | III |
| Atorvastatin | 558.64 | -1 | 112 | 2.2 | 4.46a | 37.5 | 381 | 1 | 0.02 | 86 | II |
| Budesonide | 430.54 | 0 | 93.1 | 3.3 | 13.75a | 78.4 | 28.5 | 0.8 | 0.15 | 87 | II |
| Ketoprofen | 254.3 | -1 | 54.4 | 0.19 | 4.45[a] | 4.43 | 2.9 | 0.56 | 0.017 | 6 | II |
| Lopinavir | 628.8 | 0 | 120 | 3.53 | 13.39a | 6.5 | 16.9 | 0.5 | 0.0145 | 75 | IV |
| Metoprolol | 267.4 | 1 | 50.7 | -0.13 | 9.18[b] | 48 | 67.2 | 1.07 | 0.9 | 53 | I |
| Montelukast | 586.2 | -1 | 95.7 | 4.8 | 5.7[a] | 2.7 | 9.5 | 0.62 | 0.01 | 2.9 | II |
| Nevirapine | 266.3 | 0 | 58.1 | 1.93 | 3.28[b] | 2.67 | 213 | 1[c] | 0.4 | 16 | II |
| Oxprenolol | 265.4 | 1 | 50.7 | 0.14 | 9.5[b] | 32.2 | 16.8 | 0.9 | 0.119 | 36 | I |
| Ticagrelor | 522.6 | 0 | 164 | 4.3 | 3.05[b] | 10.99 | 12.2 | 0.6 | 0.0001 | 64 (oral)<br>30 (colon) | II |
| Zafirlukast | 575.69 | -1 | 124 | 5.46 | 4.29[a] | 20 | 70 | 0.66 | 0.01 | 32 | II |

**S1:4 (continued).** (B) Physicochemical and human pharmacokinetic properties of selected drug panel [1–15] (see reference list at end of Supplementary Information). Mw = molecular weight [g/mol]. PC = physiological charge. TPSA = topological polar surface area [$Å^2$]. CL = total systemic clearance [L/h]. Vd = volume of distribution [L]. B:P = blood/plasma ratio. $f_{u,plasma}$ = fraction unbound in plasma. FPE = first pass extraction [%]. BCS = biopharmaceutical classification system. For $pK_a$ values, [a] and [b] indicate acid and base, respectively.

#### Human colonic absorption data

**C**

| Compound | Colon dose | Dose volume | Colon formulation | $AUC_{0\text{-}t}$ | $Frel_{colon}$ | $Fabs_{colon}$ | $P_{eff}$ |
|---|---|---|---|---|---|---|---|
| Atenolol | 10 | 12.65 | Solution | 0.010 | 5 | 3 | 0.013[#] |
| Budesonide | 3 | 6.2 | Solution | 0.004 | 53 | 53 | -- |
| Ketoprofen | 5 | 12.65 | Solution | 0.736 | 88 | 88 | 3.37[#] |
| Metoprolol | 19.5 | 6 | Solution | 0.108 | 100 | 95 | 1.30[#] |
| | | 6 | Powder | 0.139 | 94 | -- | |
| Oxprenolol | 80 | 5 | Solution | 1.10 | 82 | 74 | -- |
| | | 5 | Powder | 1.92 | 100 | -- | |
| Ticagrelor | 100 | 0.7 | Suspension | 1.53 | 18 | -- | -- |

**S1:4 (continued).** (C) Colon dose [mg]. Dose volume [mL]. $AUC_{0\text{-}t}$, Area under the curve between time zero and the last observed time point [μg × h/mL]. $Frel_{colon}$, Relative systemic bioavailability after administration to colon as compared to oral/duodenal administration [%]. $Fabs_{colon}$ = Observed fraction absorbed after colon administration [%], estimated by $Fabs_{oral}$ × $Frel_{colon}$. $Fabs_{oral}$ = Observed fraction absorbed after oral administration [%]. $P_{eff}$ = Effective permeability in the colon [$\times 10^{-4}$ cm/s]. All values are from [9], except those marked with a pound sign ($^{\#}$), which denotes values from *in vivo* intraluminal colonic administration of a drug cocktail (10 mL containing 10 mg atenolol 10 mg [37.5 µmol], 25 mg metoprolol [93.5 µmol], and 5 mg ketoprofen [19.7 µmol]) as reported in [16] (see reference list at end of Supplementary Information).

#### Chemical representativeness within approved drug space

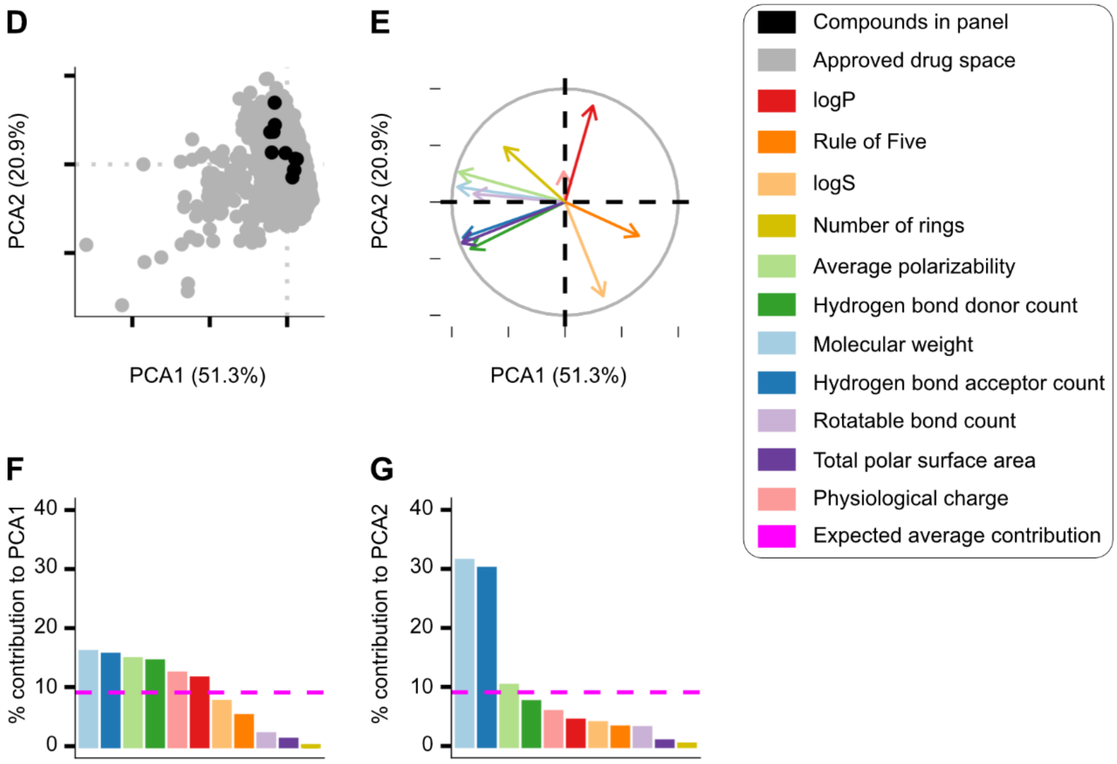

**S1:4 (continued).** (D) Principal component analysis of approved drugs (based on DrugBank database) with compounds in 11-drug panel highlighted. (E) Loadings and contributions post-scaling and centering. (F) Contribution of variables to Dimension 1 (51.3% of variance explained). (G) Contribution of variables to Dimension 2 (20.9% of variance explained).

### ADME information

## H

| Compound | Efflux transporters | | Phase I metabolism | Phase II metabolism | | Other |
|---|---|---|---|---|---|---|
| | P-gp (ABCB1) | BCRP (ABCG2) | CYP3A4 | SULTs | UGTs | |
| Atenolol | X | | | | UGT1A6 | CYP2D6<br>ABCB11, SLCO1A2, SLCO2B1 |
| Atorvastatin | X | X | X | | UGT1A1, UGT1A3, UGT2B7 | CYP2C8/9/19, CYP2D6<br>ABCB11, ABCC1/2/4/5, SLCO1A2, SLCO1B1/3, SLCO2B1 |
| Budesonide | X | | X | SULT2A1 | | CYP2A6, CYP1B1, CYP2B6, CYP2C8/9/19<br>ABCB11, SLC22A8, SLCO1A2 |
| Ketoprofen | | | | | UGT1A1 | CYP2C8/9<br>ABCC4, SLCO1A2, SLC16A1, SLC22A6/7/8/11 |
| Lopinavir | X | (X) | X | | UGT1A1, UGT1A3, UGT1A4 | CYP1A2, CYP2B6, CYP2C9/19, CYP2D6<br>ABCB11, SLCO1B1/3 |
| Metoprolol | | | X | | UGT1A1 | CYP2B6, CYP2C9, CYP2D6<br>(ABCB11), SLC22A2 |
| Montelukast | | | X | | UGT1A3 | CYP2A6, CYP2C8/9, PTGS1<br>SLCO2B1 |
| Nevirapine | | (X) | X | X | X | CYP1A2/6, CYP2B6, CYP2C9/19, CYP2D6<br>(ABCB11), ABCC10, SLC22A1 |
| Oxprenolol | | | | | X | CYP2D6<br>SLC22A2 |
| Ticagrelor | X | (X) | X | | UGT1A9 | CYP2B6, CYP2C9<br>SLCO1B1 |
| Zafirlukast | | X | X | | UGT1A, UGT2B | CYP1A2, CYP2C8/9/19, CYP2D6, PTGS1<br>SLC10A1 |

**S1:4 (continued).** (H) Data compiled from [17–32] (see reference list at end of Supplementary Information). X = substrate. For protein families, the subtype is listed (where known). Substrates with inhibitory function listed in parentheses (X).

### S1:5 UHPLC-MS/MS parameters

#### UHPLC parameters

**A**

| Time [min] | %A (95% water + 4.9% acetonitrile + 0.1% formic acid) | %B (acetonitrile + 0.1% formic acid) | Flow [mL/min] | Diverter valve |
|---|---|---|---|---|
| 0 | 98 | 2 | 0.5 | To waste |
| 0.4 | 98 | 2 | 0.5 | To MS |
| 0.5 | 98 | 2 | 0.5 | To MS |
| 1.2 | 5 | 95 | 0.5 | To MS |
| 1.6 | 5 | 95 | 0.5 | To MS |
| 1.9 | 5 | 95 | 0.5 | To waste |
| 1.95 | 98 | 2 | 0.5 | To waste |
| 2.00 | 98 | 2 | 0.5 | To waste |

**S1:5.** (A) Gradient parameters for analysis of the compounds listed below.

#### MS/MS parameters

**B**

| Compound | Parent [m/z] | Daughter [m/z] | Dwell [s] | Cone [V] | Collision [V] | $R_t$ [min] |
|---|---|---|---|---|---|---|
| 1-OH-Midazolam | 342.1000 | 203.0000 | 0.02 | 34 | 28 | 1.20 |
| 4-Methyl-umbelliferone | 175.0000 | 133.0000 | 0.02 | 47 | 30 | 1.22 |
| 4-OH-Midazolam | 342.1000 | 234.0300 | 0.02 | 36 | 22 | 1.20 |
| Acetaminophen | 152.0958 | 109.9100 | 0.02 | 40 | 20 | 0.80 |
| Atenolol | 267.1000 | 145.2000 | 0.02 | 28 | 26 | 1.22 |
| Atorvastatin | 559.2000 | 440.2000 | 0.02 | 30 | 22 | 1.45 |
| Budesonide | 431.3000 | 147.0250 | 0.02 | 18 | 36 | 1.44 |
| Ketoprofen | 254.9000 | 77.000 | 0.02 | 56 | 46 | 1.40 |
| Lopinavir | 629.5000 | 155.1000 | 0.02 | 22 | 46 | 1.51 |
| Metoprolol | 268.2000 | 116.0000 | 0.02 | 26 | 18 | 1.11 |
| Montelukast | 586.3380 | 278.1370 | 0.02 | 20 | 30 | 1.69 |
| Midazolam | 326.0300 | 291.0400 | 0.02 | 44 | 28 | 1.19 |
| Nevirapine | 267.1000 | 226.2000 | 0.02 | 34 | 23 | 1.23 |
| Oxprenolol | 266.2543 | 72.0316 | 0.02 | 14 | 14 | 1.16 |
| Ticagrelor | 523.1000 | 153.1030 | 0.02 | 8 | 44 | 1.50 |
| Warfarin | 309.1662 | 163.1476 | 0.02 | 34 | 16 | 1.43 |
| Zafirlukast | 576.3380 | 337.1590 | 0.02 | 18 | 16 | 1.58 |

**S1:5 (continued).** (B) Mass analysis parameters for the small molecules used in this study.

### S1:6 Intracellular bioavailability

#### Formulae relating to intracellular bioavailability

**A**

##### Total accumulation (Kp)

$$Kp = \frac{A_{cell}}{V_{cell}} \times \frac{1}{C_{buffer}}$$

where $A_{cell}$ is the amount of compound in the cells, $V_{cell}$ is the cell volume (calculated from a scaling factor of 6.5 μL/mg protein), and $c_{buffer}$ is the compound concentration in the buffer.

##### Intracellular unbound fraction ($f_u$) via unbound fraction in cell homogenate ($f_{u,hom}$)

$$f_{u,hom} = \frac{c_{buffer}}{c_{hom}}$$

where $c_{buffer}$ is the compound concentration in the buffer and $c_{hom}$ is the compound concentration in cell homogenate.

$$f_u = \frac{1}{\frac{1}{D} \times \left(\frac{1}{f_{u,hom}} - 1\right) + 1}$$

where $f_{u,hom}$ is the unbound fraction in cell homogenate and D is the dilution factor, obtained using a scaling factor of 6.5 μL cell volume/mg protein.

##### Intracellular bioavailability ($F_{ic}$)

$$F_{ic} = Kp \times f_u$$

**S1:5.** (A) Formulae relating to intracellular bioavailability.

### Total accumulation (Kp)

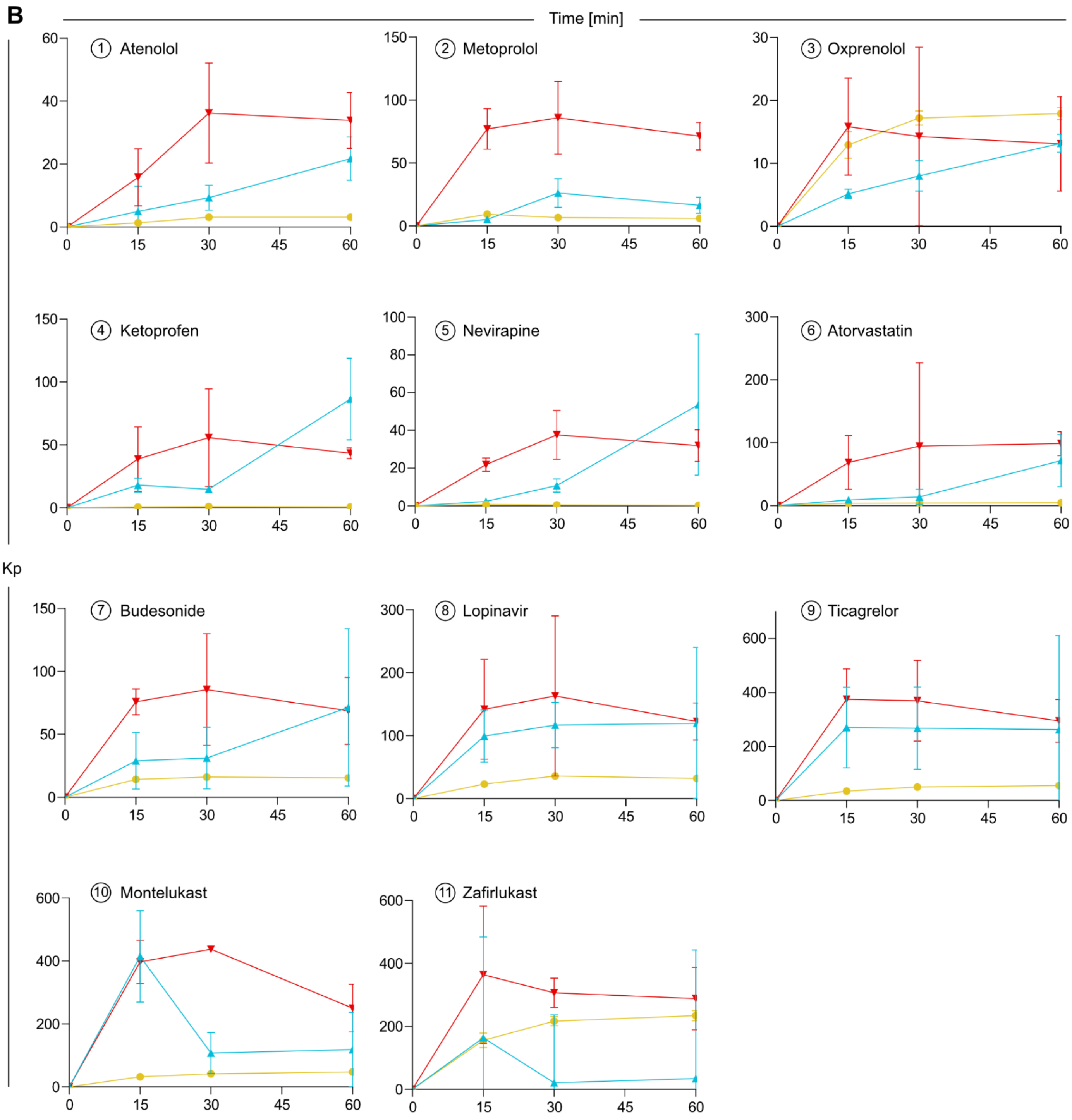

**S1:6 (continued).** (B) Total accumulation (Kp, geomean ± SEM) of the 11-drug panel used in this study.

#### Tabulated values for Kp, $f_u$, and $F_{ic}$

**C**

| Compound | Model | Kp | $f_u$ | $F_{ic}$ |
|---|---|---|---|---|
| Atenolol | HEK293 | 3.13 ± 0.15 | 0.27 ± 0.04 (n = 2) | 0.84 ± 0.14 |
| Atenolol | 4AD | 34 ± 9 | 0.078 ± 0.051 | 2.66 ± 1.86 |
| Atenolol | 4BD | 22 ± 7 (n = 5) | 0.051 ± 0.022 | 1.12 ± 0.58 |
| Metoprolol | HEK293 | 6.06 ± 1.32 | 0.77 ± 0.13 | 4.66 ± 1.29 |
| Metoprolol | 4AD | 71 ± 11 | 0.027 ± 0.0056 (n = 2) | 1.95 ± 0.50 |
| Metoprolol | 4BD | 16 ± 6 | 0.033 ± 0.0091 | 0.56 ± 0.26 |
| Oxprenolol | HEK293 | 18 ± 1 | 0.37 ± 0.11 | 6.62 ± 1.98 |
| Oxprenolol | 4AD | 13 ± 8 | 0.067 ± 0.025 | 0.88 ± 0.60 |
| Oxprenolol | 4BD | 13 ± 1 | 0.072 ± 0.017 | 0.95 ± 0.24 |
| Ketoprofen | HEK293 | 0.76 ± 0.25 | 0.25 ± 0.057 | 0.19 ± 0.076 |
| Ketoprofen | 4AD | 43 ± 4 | 0.11 ± 0.25 | 4.66 ± 10.98 |
| Ketoprofen | 4BD | 86 ± 32 (n = 5) | 0.045 ± 0.054 | 3.89 ± 4.87 |
| Nevirapine | HEK293 | 0.31 ± 0.06 | 0.27 ± 0.054 | 0.082 ± 0.022 |
| Nevirapine | 4AD | 32 ± 8 | 0.10 ± 0.11 | 3.33 ± 3.63 |
| Nevirapine | 4BD | 54 ± 37 | 0.044 ± 0.028 | 2.34 ± 2.21 |
| Atorvastatin | HEK293 | 4.5 ± 0.32 | 0.025 ± 0.0051 | 0.12 ± 0.025 |
| Atorvastatin | 4AD | 99 ± 19 | 0.0088 ± 0.0022 | 0.87 ± 0.27 |
| Atorvastatin | 4BD | 72 ± 41 | 0.014 ± 0.0011 | 0.98 ± 0.57 |
| Budesonide | HEK293 | 15 ± 1 | 0.038 ± 0.0046 | 0.59 ± 0.080 |
| Budesonide | 4AD | 69 ± 27 | 0.020 ± 0.0069 | 1.40 ± 0.72 |
| Budesonide | 4BD | 71 ± 63 | 0.0095 ± 0.0012 | 0.68 ± 0.60 |
| Lopinavir | HEK293 | 32 ± 3 | 0.0093 ± 0.0014 | 0.30 ± 0.052 |
| Lopinavir | 4AD | 123 ± 30 | 0.0029 ± 0.00060 | 0.35 ± 0.11 |
| Lopinavir | 4BD | 120 ± 120 | 0.0047 ± 0.0019 | 0.56 ± 0.60 |
| Ticagrelor | HEK293 | 55 ± 3 | 0.00063 ± 0.00023 | 0.035 ± 0.012 |
| Ticagrelor | 4AD | 295 ± 80 | 0.00031 ± 0.000025 | 0.092 ± 0.026 |
| Ticagrelor | 4BD | 263 ± 349 | 0.00055 ± 0.00017 | 0.14 ± 0.20 |
| Montelukast | HEK293 | 48 ± 4 | 0.00067 ± 0.000021 | 0.032 ± 0.0031 |
| Montelukast | 4AD | 250 ± 76 (n = 5) | 0.000064 ± 0.000012 | 0.016 ± 0.0057 |
| Montelukast | 4BD | 119 ± 118 (n = 2) | 0.00019 ± 0.000024 | 0.023 ± 0.023 |
| Zafirlukast | HEK293 | 234 ± 16 | 0.0014 ± 0.00032 | 0.33 ± 0.079 |
| Zafirlukast | 4AD | 288 ± 99 (n = 5) | 0.00083 ± 0.00011 | 0.24 ± 0.088 |
| Zafirlukast | 4BD | 34 ± 409 (n = 4) | 0.0019 ± 0.0017 | 0.066 ± 0.78 |

**S1:6 (continued).** (C) Tabulated values for steady-state Kp, $f_u$, and $F_{ic}$ given as geometric mean ± SEM. For Kp, n = 6 unless otherwise indicated. For $f_u$, n = 3 unless otherwise indicated.

#### Cell and organoid dimension calculations

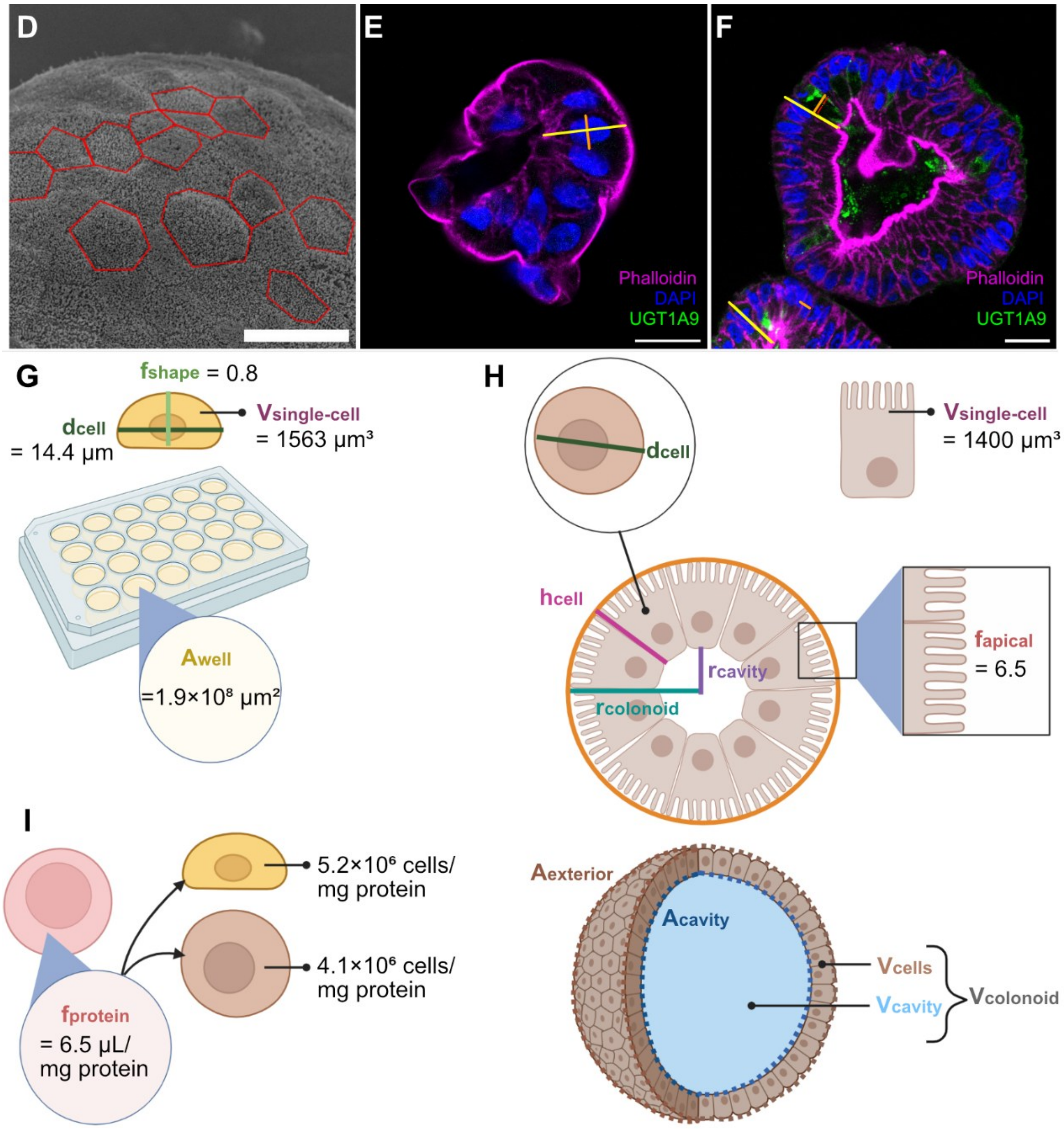

**S1:6 (continued).** (D) 13 individually discernible cells from 4AD SEM image (scale bar 10 µm) used for estimating $d_{cell}$ (via the Feret diameter). (E-F) Example ICC images from 4AD and 4BD, respectively, used to estimate $h_{cell}$ (yellow) and $d_{cell}$ (orange) (scale bars 20 µm). (G) Schematic showing relevant factors for the dimension calculation for HEK293 cells yielding the $f_{protein}$ scaling value shown in panel F; $d_{cell}$ = cellular diameter, $f_{shape}$ = shape factor to correct for flattening, $V_{single\text{-}cell}$ = literature value for the volume of a single cell. (H) Schematic showing relevant factors for the dimension calculation for 4AD and 4BD colonoids, given in the table below; $d_{cell}$ = cellular diameter (shown in inset depicting cross-sectional view from above), $h_{cell}$ = cellular height, $r_{lumen}$ = lumen radius, $r_{colonoid}$ = colonoid radius, $A_{colonoid}$ = area of the sphere defined by $r_{colonoid}$, $A_{exterior}$ = exposed surface area of colonoid without accounting for $f_{apical}$, $A_{lumen}$ = surface area of sphere defined by $r_{cavity}$, $V_{cells}$ = total cellular volume, $V_{cavity}$ = volume of internal cavity, $V_{colonoid}$ = total volume occupied by colonoid, refer to panel D figure legend for other parameters. (I) Number of cells per mg protein for HEK293 cells and colonoid cells using a literature-defined scaling factor, $f_{protein}$.

#### Tabulated organoid dimensions

**J**

| | **4AD** | **4BD** |
|---|---|---|
| $d_{cell}$ [μm]* | 8.93 | 8.41 |
| $h_{cell}$ [μm]# | 25.19 | 23.25 |
| $A_{colonoid}$ [μm$^2$]† | 14002 | 9155 |
| $r_{colonoid}$ [μm] | 67 | 54 |
| $r_{cavity}$ [μm] | 42 | 31 |
| Cells per colonoid | 600 | 417 |
| $V_{single\text{-}cell}$ [μm$^3$] | 1576 | 1290 |
| $V_{cells}$ [μm$^3$] | $9.45\times10^5$ | $5.37\times10^5$ |
| $V_{cavity}$ [μm$^3$] | $3.01\times10^5$ | $1.22\times10^5$ |
| $V_{colonoid}$ [μm$^3$] | 1.25×106 | $6.59\times10^5$ |
| $V_{cells}/V_{cavity}$ | 3.14 | 4.42 |
| $A_{exterior}$ [μm$^2$] | $5.60\times10^4$ | $3.66\times10^4$ |
| $A_{cavity}$ [μm$^2$] | $2.17\times10^4$ | $1.19\times10^4$ |
| $A_{apical}$ [μm$^2$] | $3.64\times10^5$ | $7.72\times10^4$ |
| $A_{basal}$ [μm$^2$] | $2.17\times10^4$ | $3.66\times10^4$ |
| $A_{apical}/A_{basal}$ | 16.8 | 2.1 |

**S1:6 (continued).** (J) Colonoid dimension calculations assuming spherical colonoids, cylindrical cells, and an apical surface enlargement factor ($f_{apical}$) of 6.5.

*4AD average value from n = 13 cells (SEM images) and n = 13 colonoids (ICC images), 4BD average value from n = 13 colonoids

#Average values from ICC for n = 13 colonoids each for 4AD and 4BD

†Average values from brightfield microscopy (see Fig. 1) for n = 331 colonoids and n = 349 colonoids for 4AD and 4BD, respectively

### S1:7 Lipidomics

#### Method details

**A**

Samples were extracted via a chloroform:methanol (2:1) phase extraction. The chloroform phase (= extract) was stored at -80°C until UHPLC-MS/MS analysis [33]. The extraction mix included internal standards (IS), TG(16:0/16:0/16:0)-$^{13}$C3, Cer(d18:1/16:0)-D30, and PC(18:0/18:0)-D70, added at identical concentrations to all samples prior to extraction. As a quality control (QC) sample, a mix of extract from all samples was used, generating a dilution series. A 1290 Infinity UHPLC (Agilent) was used, with a gradient optimized for plasma lipid extracts. The starting gradient was (53:13:34) (acetonitrile:isopropanol:water), ending at (11:89) (acetonitrile:isopropanol). All mobile phases included 10 mM ammonium format and 0.1% formic acid. For separation, an Acquity UPLC CSH 2.1x50 1.7µm C18 column (Waters) was used [34]. An QTOF 6546 instrument ("ODIN") (Agilent) was used for mass detection, operating in positive electrospray ionization (ESI+) mode over an m/z range of 70-1700, collecting 4 scans. MS/MS of major peaks in the QC sample was run in additional sample injections at 2 different collision energies (25 V respective 40 V).

**S1:7.** (A) Lipidomic method details.

#### Protein-scaled lipid intensities

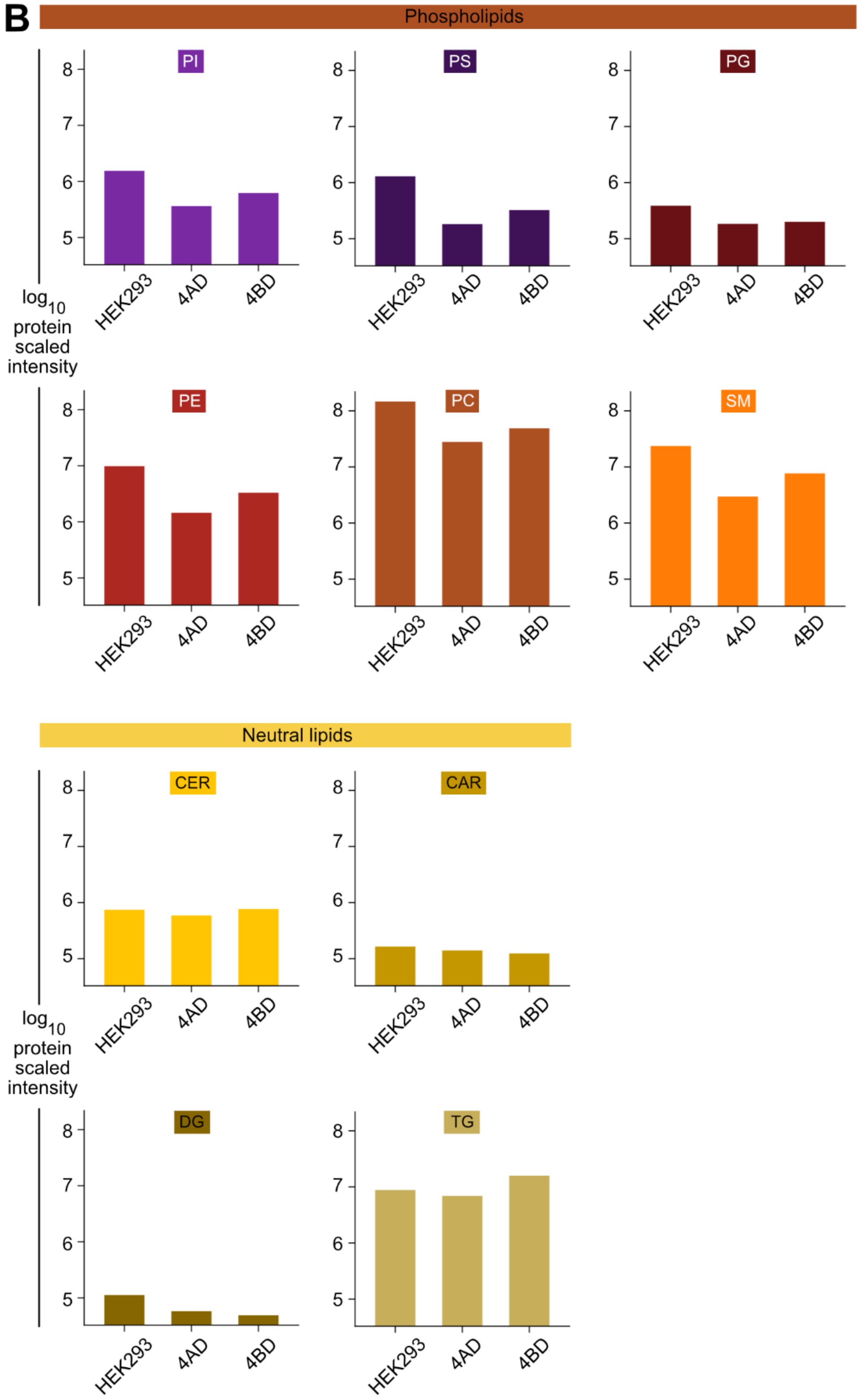

**S1:7 (continued).** (B) Mean protein-scaled intensities for individual phospholipid and neutral lipid classes (intensity/mg/mL protein in sample).

### Enrichment of proteins relating to chylomicron production

C

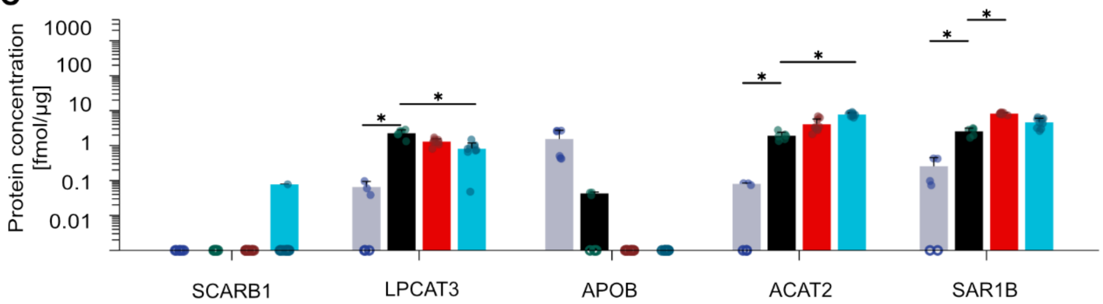

D

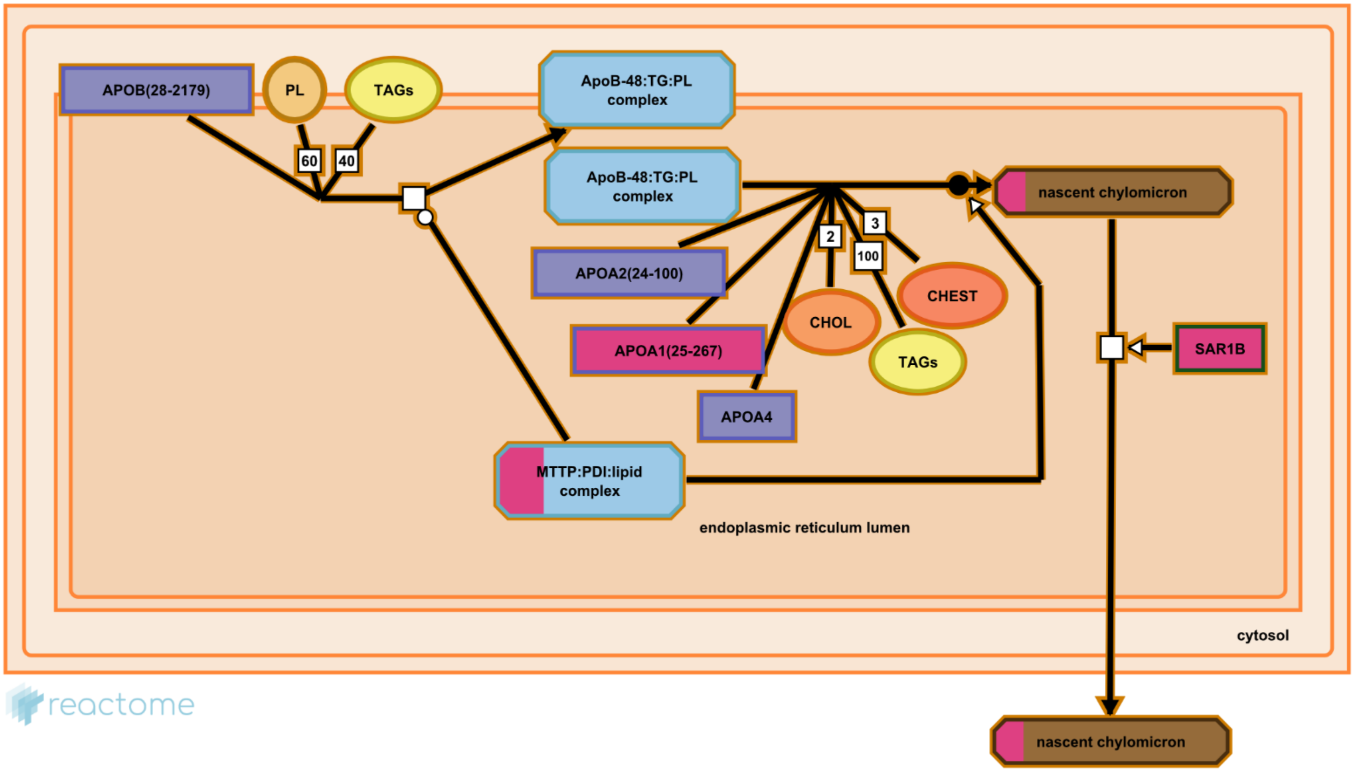

E

| GO biological process PANTHER Classification System | Associated genes | Fold enrichment | p-value | FDR |
|---|---|---|---|---|
| Chylomicron assembly (GO:0034378) | LPCAT3 MTTP* APOA4 | -- | -- | -- |
| Low-density lipoprotein particle clearance (GO:0034383) | 14 | 3.23 | $1.13 \times 10^{-4}$ | $1.51 \times 10^{-3}$ |
| Lipoprotein localization (GO:0044872) | 16 | 2.91 | $8.29 \times 10^{-4}$ | $8.54 \times 10^{-3}$ |
| Lipoprotein transport (GO:0042953) | 15 | 2.79 | $2.46 \times 10^{-3}$ | $2.12 \times 10^{-2}$ |

**S1:7 (continued).** (C) Protein expression values for chylomicron-mediating proteins, based on global proteomics. Hollow circles on the x axis indicate replicates where the protein of interest was not quantifiable. *p-value < 0.05 (DEqMS with Benjamini-Hochberg multiple testing correction). (D) Schematic showing pathways involved in chylomicron assembly, as exported from Reactome. Pink highlight indicates percentage of proteins (relative to full bar length) associated with a given pathway step that were quantified in 4AD and 4BD colonoids. Other colors in accordance with Fig. 9, with protein-lipid complexes highlighted in blue. APO = apolipoprotein. PL = phospholipids. TAGs = tri(acyl)glycerides. MTTP = microsomal triglyceride transfer protein. PDI = protein disulfide isomerase. CHOL = cholesterol. CHEST = cholesteryl esters. SAR1B = secretion associated Ras related GTPase 1B. (E) Gene enrichment analysis in PANTHER. An asterisk (*) indicates that the protein was not quantifiable in 4AD and 4BD. LPCAT3 = lysophosphatidylcholine acyltransferase 3. FDR = false discovery rate. Note that only 3 chylomicron assembly associated genes are annotated in the PANTHER database, which is too few for enrichment analysis [35–37].

### S1:8 Proteomic screening

#### Comparison by organoid donor

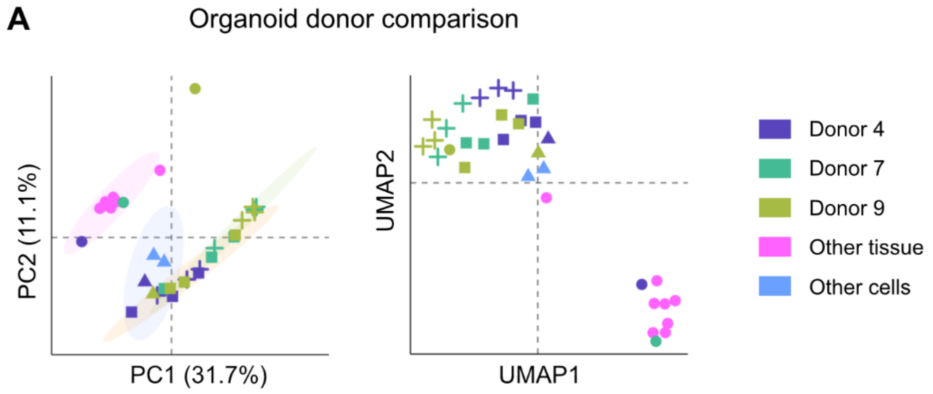

**S1:8.** (A) Dimensionality reduction techniques to assess sample variance (PCA with 95% confidence ellipses for each cluster, left) and similarity (UMAP, right), based on global proteomics.

#### Matrigel impact

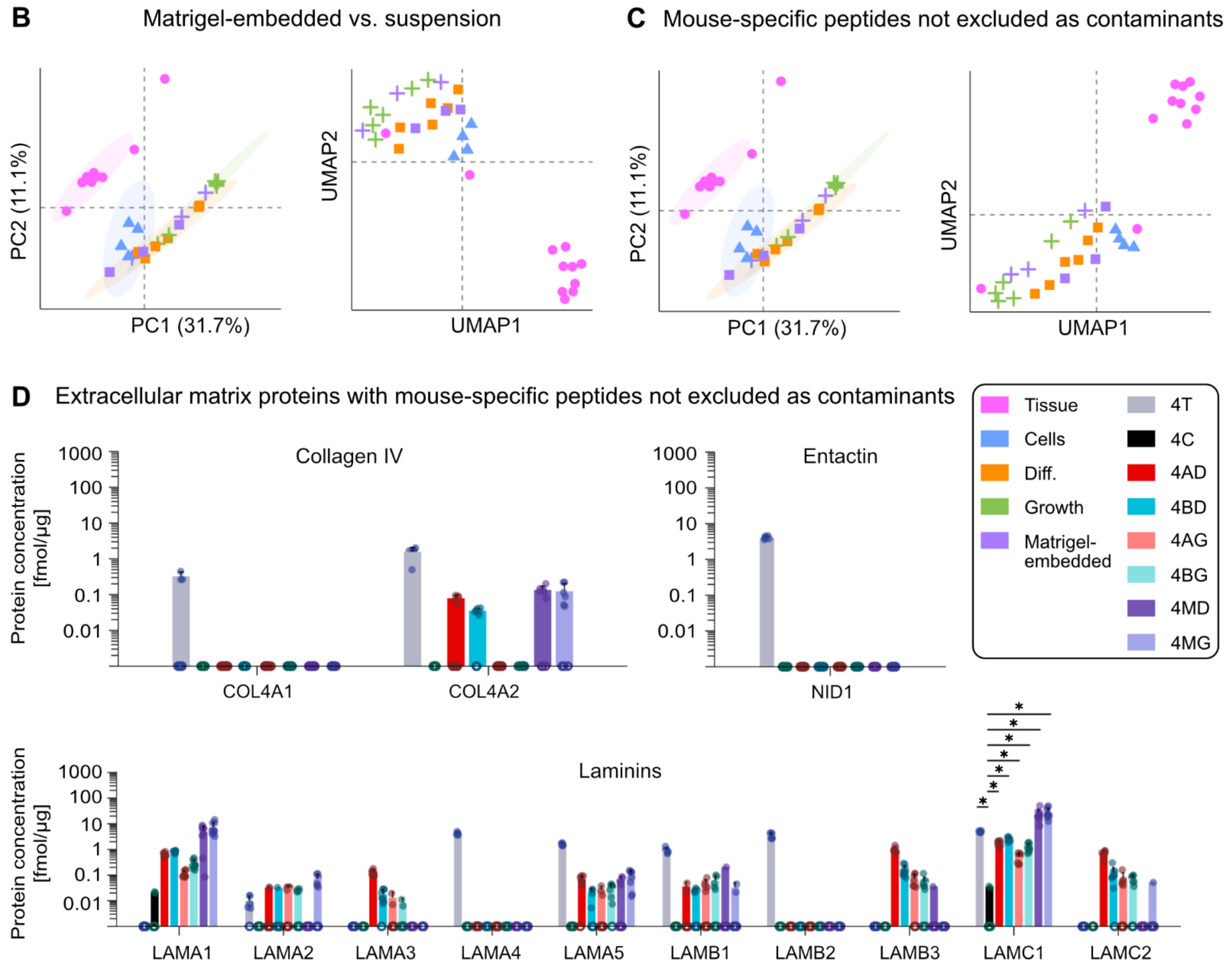

**S1:6 (continued).** (B) and (C) Dimensionality reduction techniques to assess sample variance (PCA with 95% confidence ellipses, left) and similarity (UMAP, right), based on global proteomics. (D) Barplots of protein expression based on global proteomics. Hollow circles on the x axis indicate replicates where the protein of interest was not quantifiable. *p-value < 0.05 (DEqMS with Benjamini-Hochberg multiple testing correction).

### S1:9 Additional absorption and secretion

#### Cell types barplots

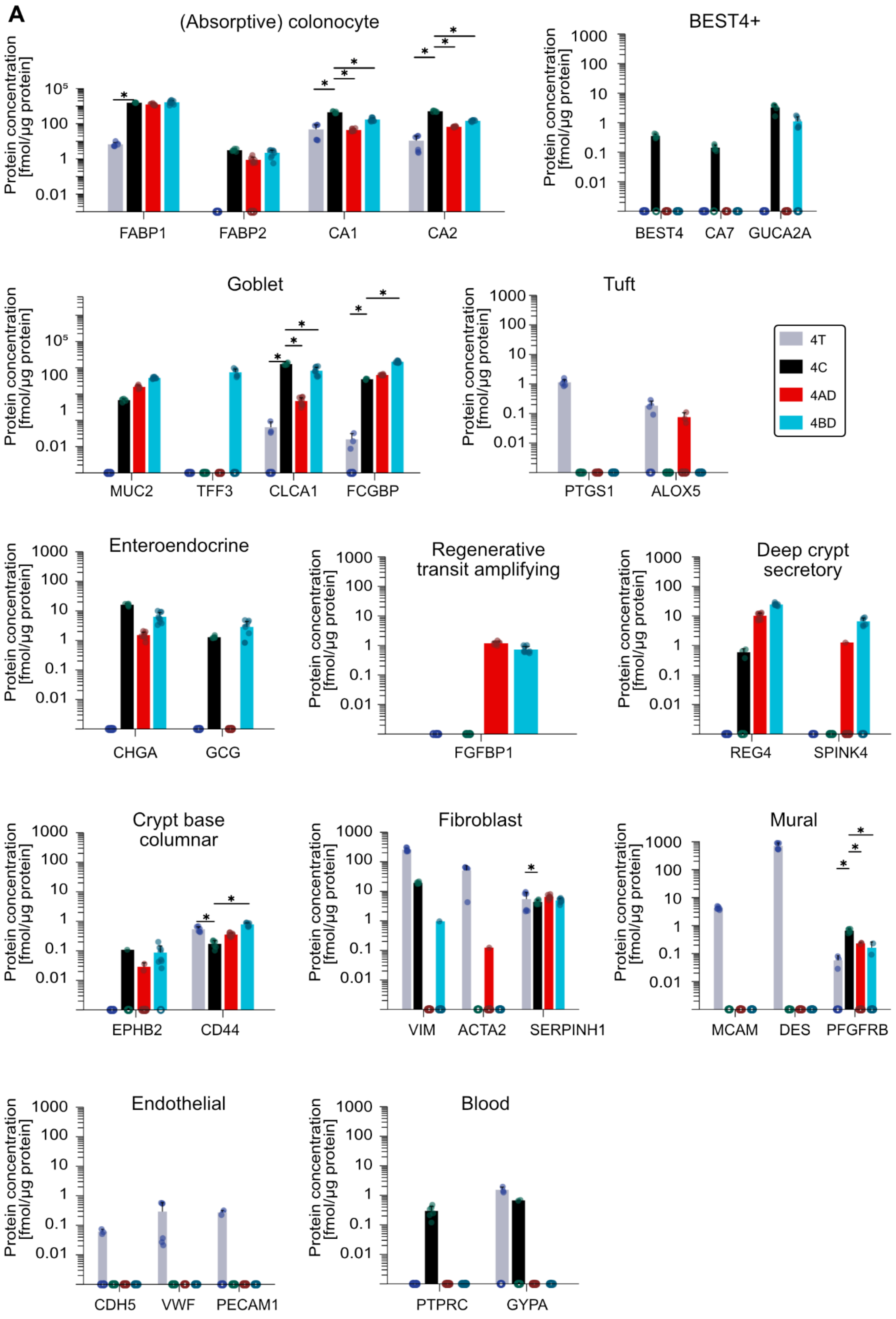

**S1:9 (continued from previous page).** (A) Protein expression levels of markers of epithelial cell types, based on global proteomics. Hollow circles on the x axis indicate replicates where the protein of interest was not quantifiable. *p-value < 0.05 (DEqMS with Benjamini-Hochberg multiple testing correction).

### Absorption and secretion

#### Nutrient processing and mucin secretion

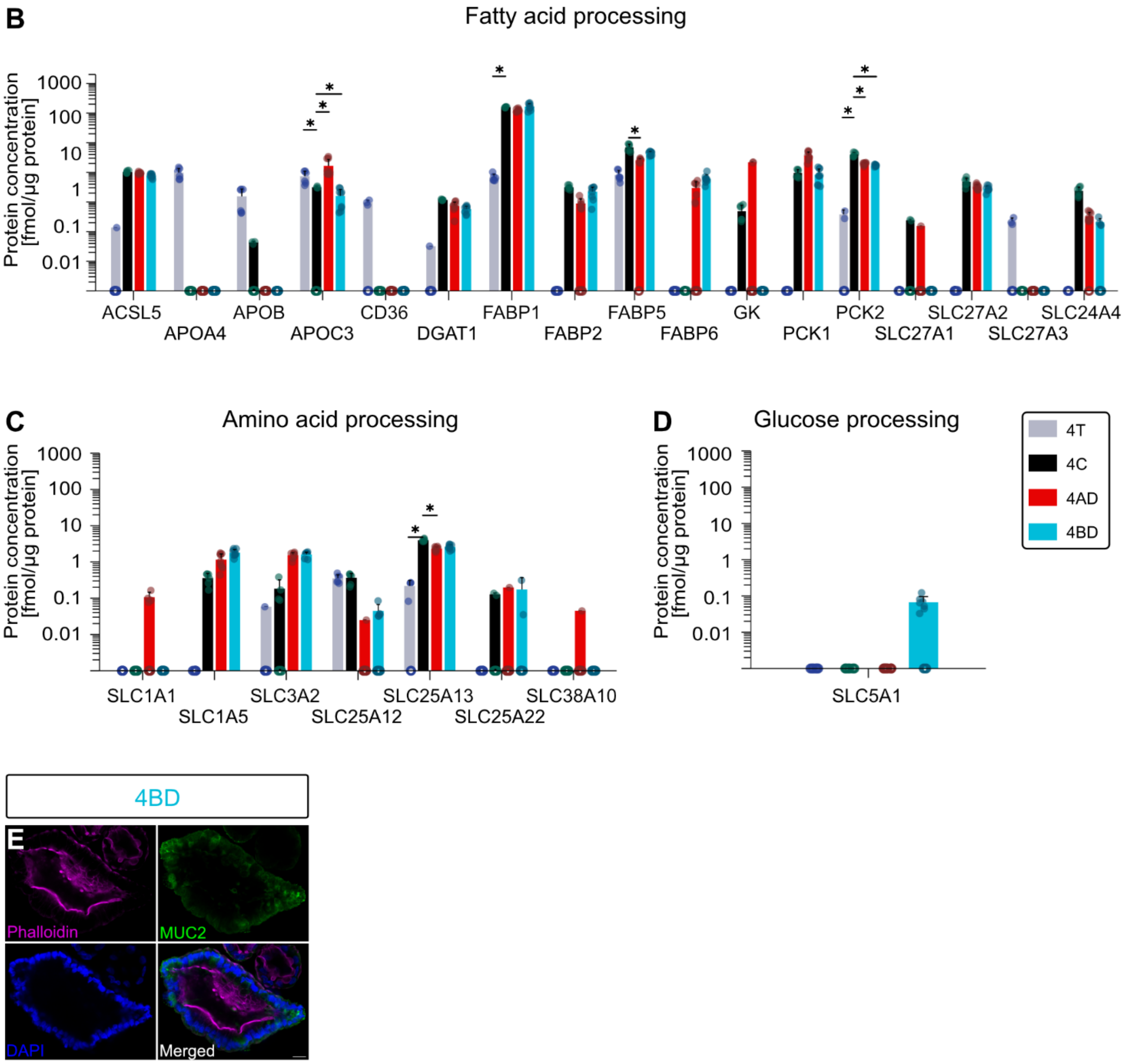

**S1:9 (continued).** (B) through (D) Protein expression of proteins relating to the processing of nutrients, based on global proteomics. Hollow circles on the x axis indicate replicates where the protein of interest was not quantifiable. (E) ICC image of MUC2 in 4BD. *p-value < 0.05 (DEqMS with Benjamini-Hochberg multiple testing correction).

#### Specialized goblet cell subpopulations

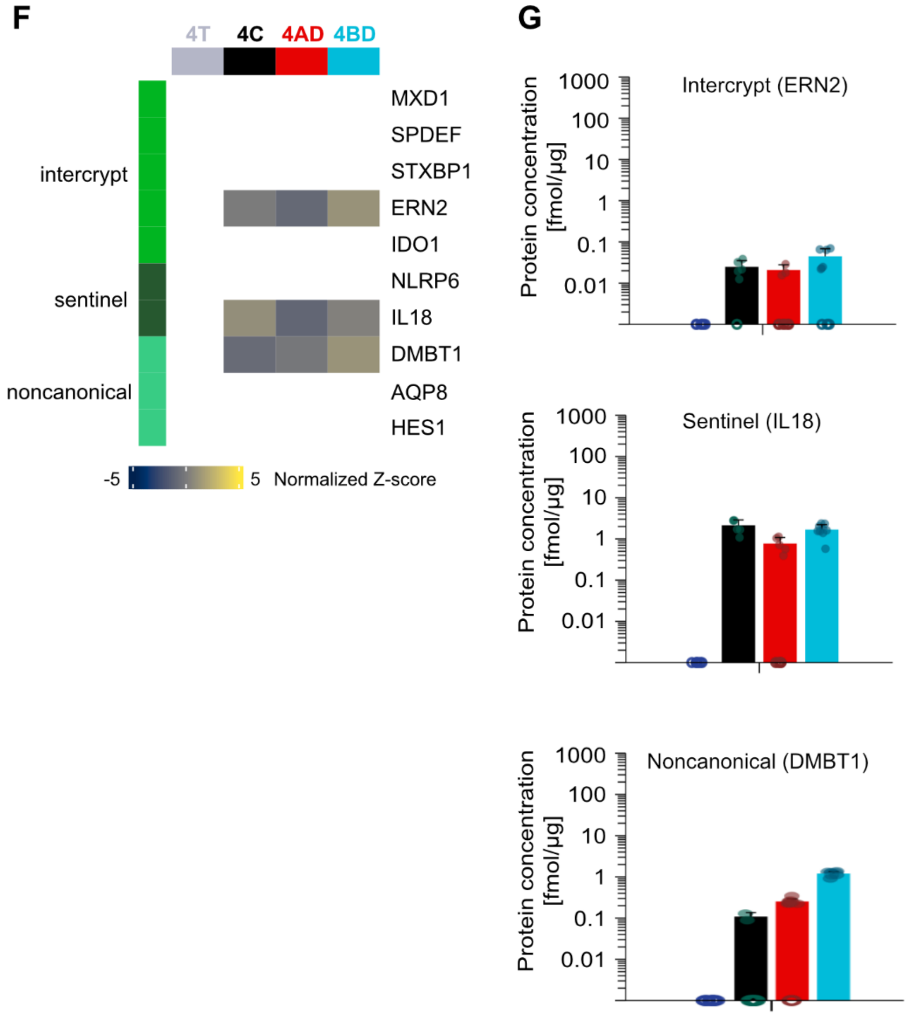

**S1:9 (continued).** (F) Heatmap of marker expression for specialized goblet cell subpopulations, many of which were not quantifiable, based on global proteomics. (G) Barplots detailing the expression levels of the markers that were quantifiable. Hollow circles on the x axis indicate replicates where the protein of interest was not quantifiable.

#### S1:10 Barriers

##### Epithelial barriers (barplots)

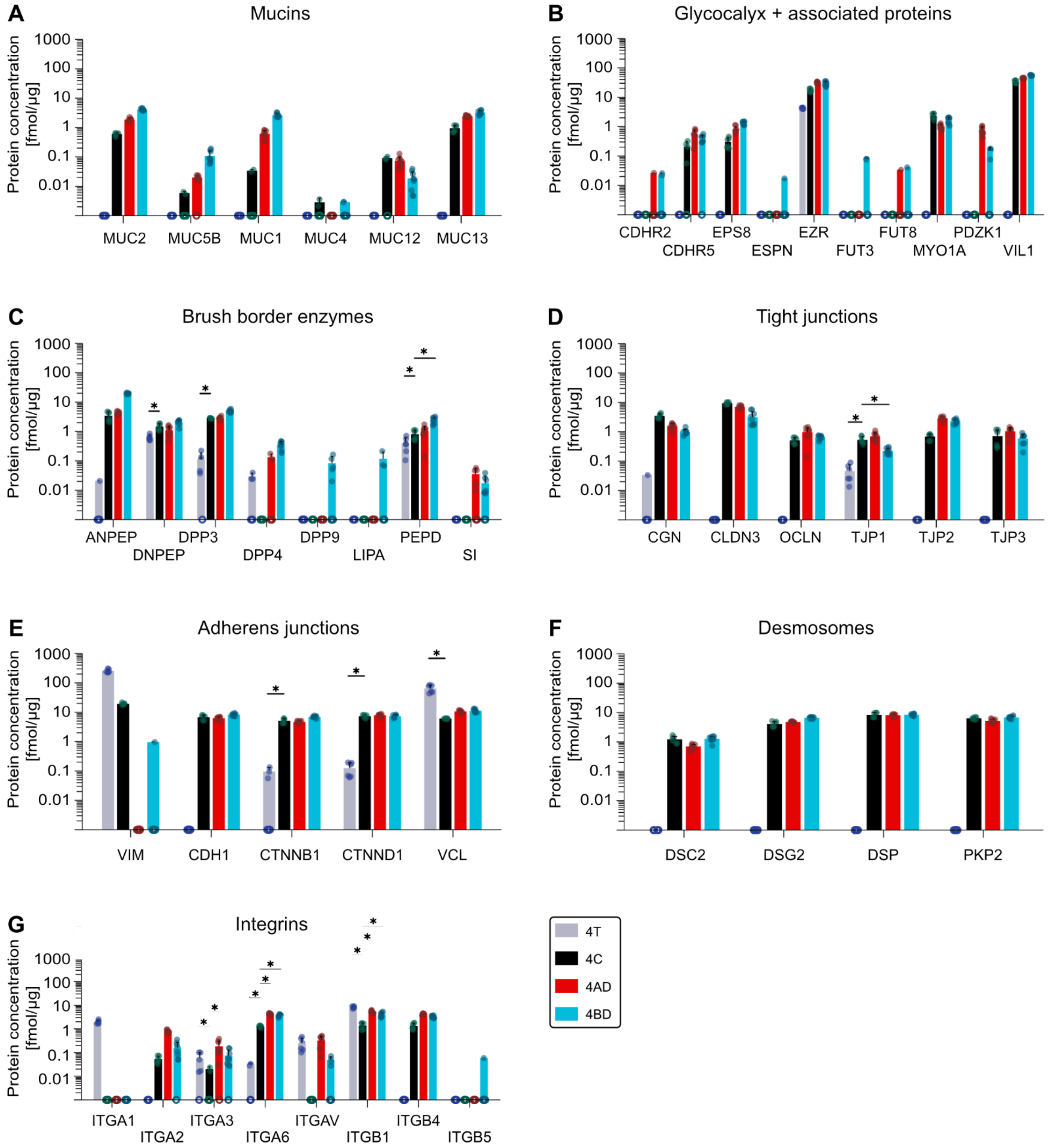

**S1:10.** (A) through (G) Protein expression of relevant barrier proteins, based on global proteomics. Hollow circles on the x axis indicate replicates where the protein of interest was not quantifiable. *p-value < 0.05 (DEqMS with Benjamini-Hochberg multiple testing correction).

### Endocytosis machinery

H

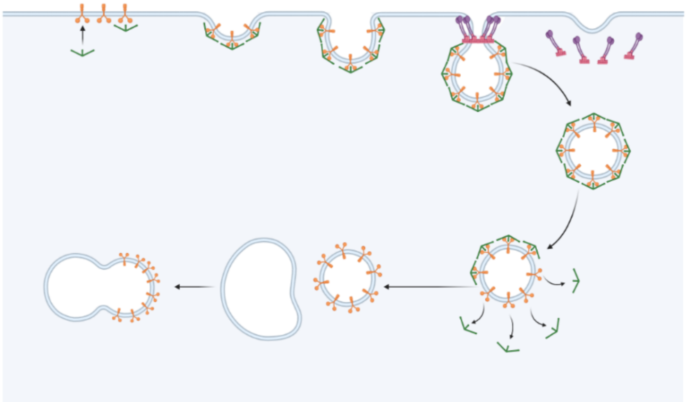

I

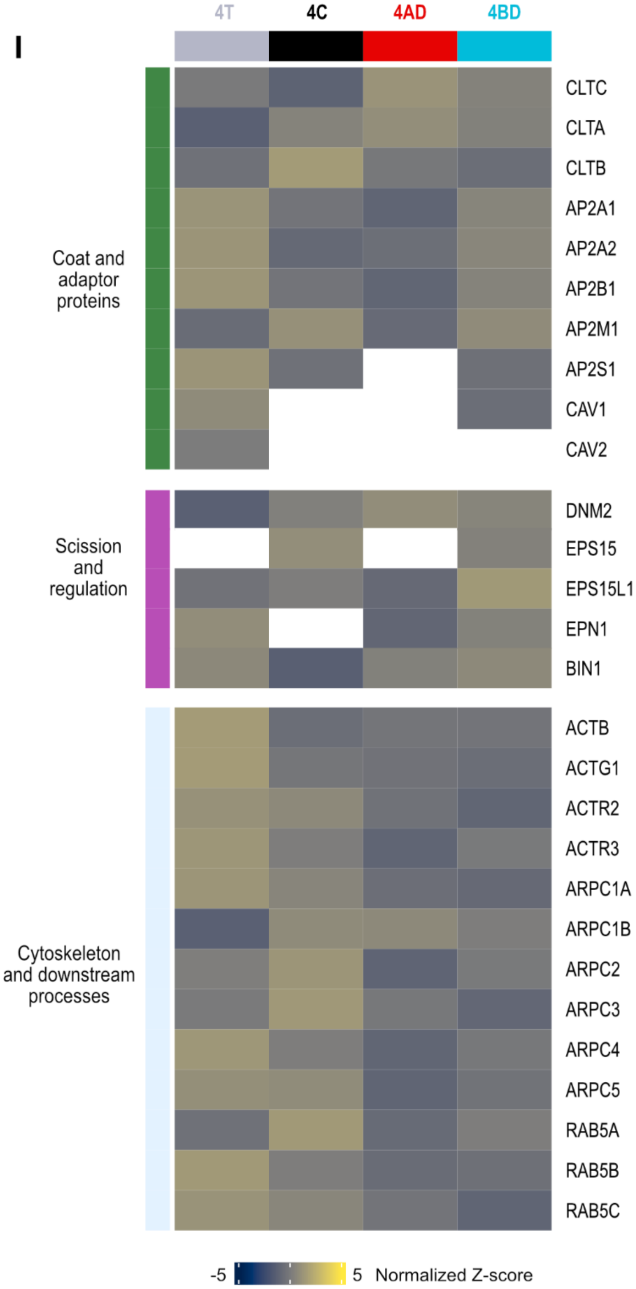

**S1:10 (continued).** (H) Schematic representation of receptor-mediated endocytosis. (I) Heatmap of endocytosis-associated proteins, based on global proteomics.
