## Supplementary Information 2 for "Native-like 3D colonoids reveal mechanisms of colonic drug disposition"

# 4T

### Gene Ontology - Biological Process

*Barplot*

**A**

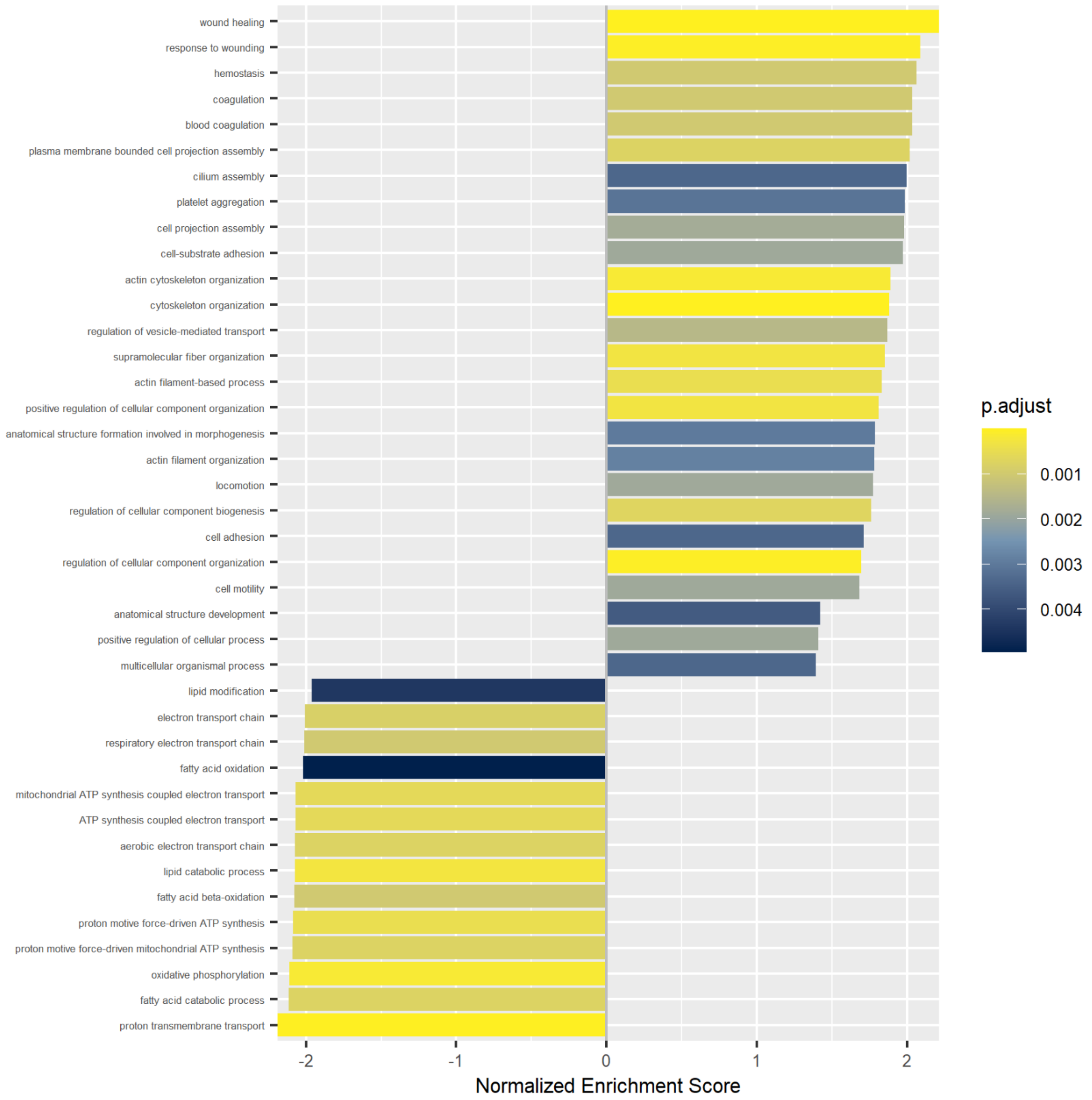

**S2.** (A) 4T vs. 4C barplot showing normalized enrichment score and p-value (adjusted according to Benjamini-Hochberg) for enriched processes based on Gene Ontology Biological Process. Based on global proteomics.

*Dotplot*

**B**

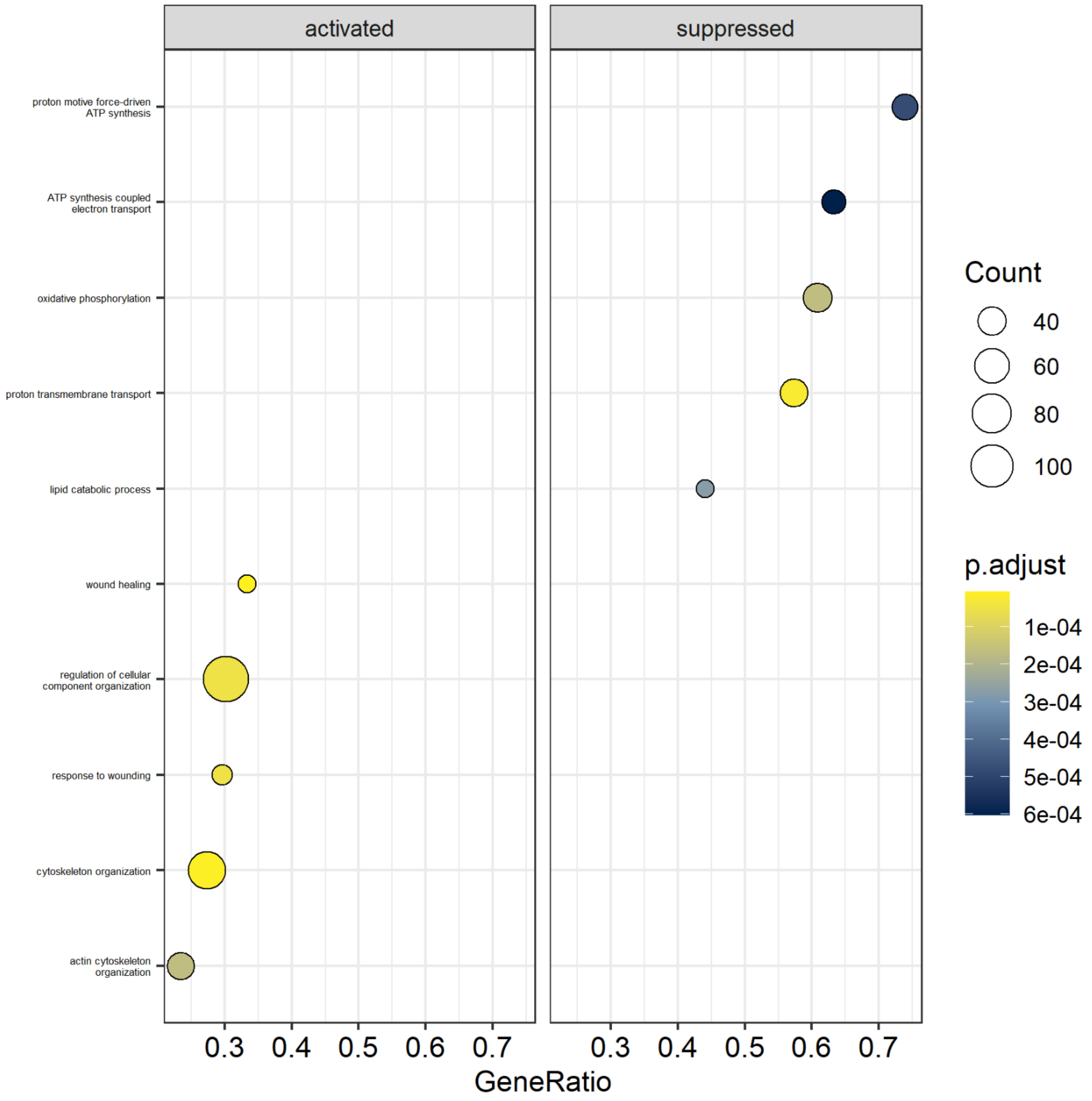

**S2 (continued).** (B) 4T vs. 4C dotplot showing activated and suppressed processes, including gene count and p-value (adjusted according to Benjamini-Hochberg) per process, based on Gene Ontology Biological Process. Based on global proteomics.

### Gene Ontology – Cellular Component

*Barplot*

**C**

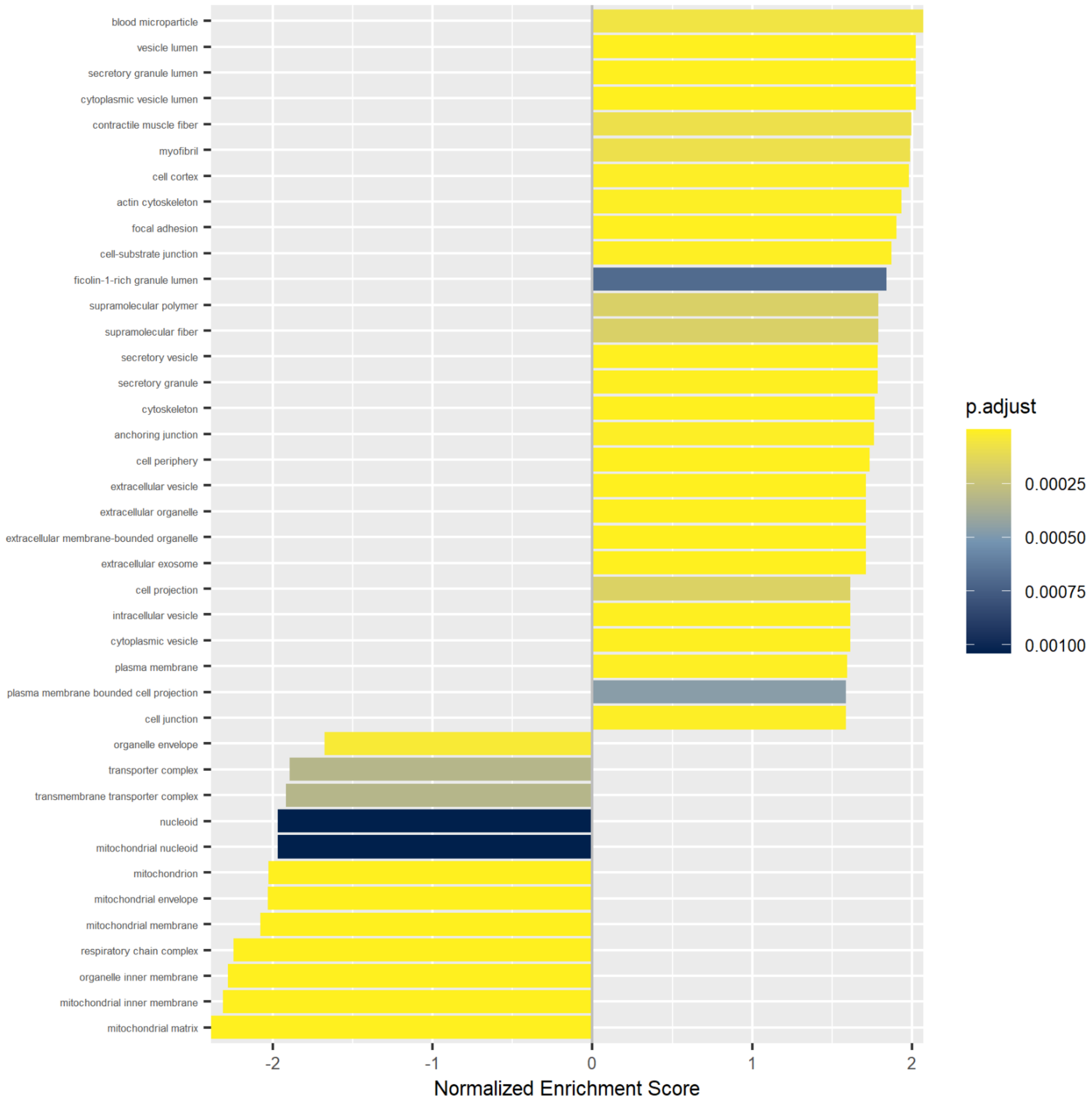

**S2 (continued).** (C) 4T vs. 4C barplot showing normalized enrichment score and p-value (adjusted according to Benjamini-Hochberg) for enriched processes based on Gene Ontology Cellular Component. Based on global proteomics.

*Dotplot*

**D**

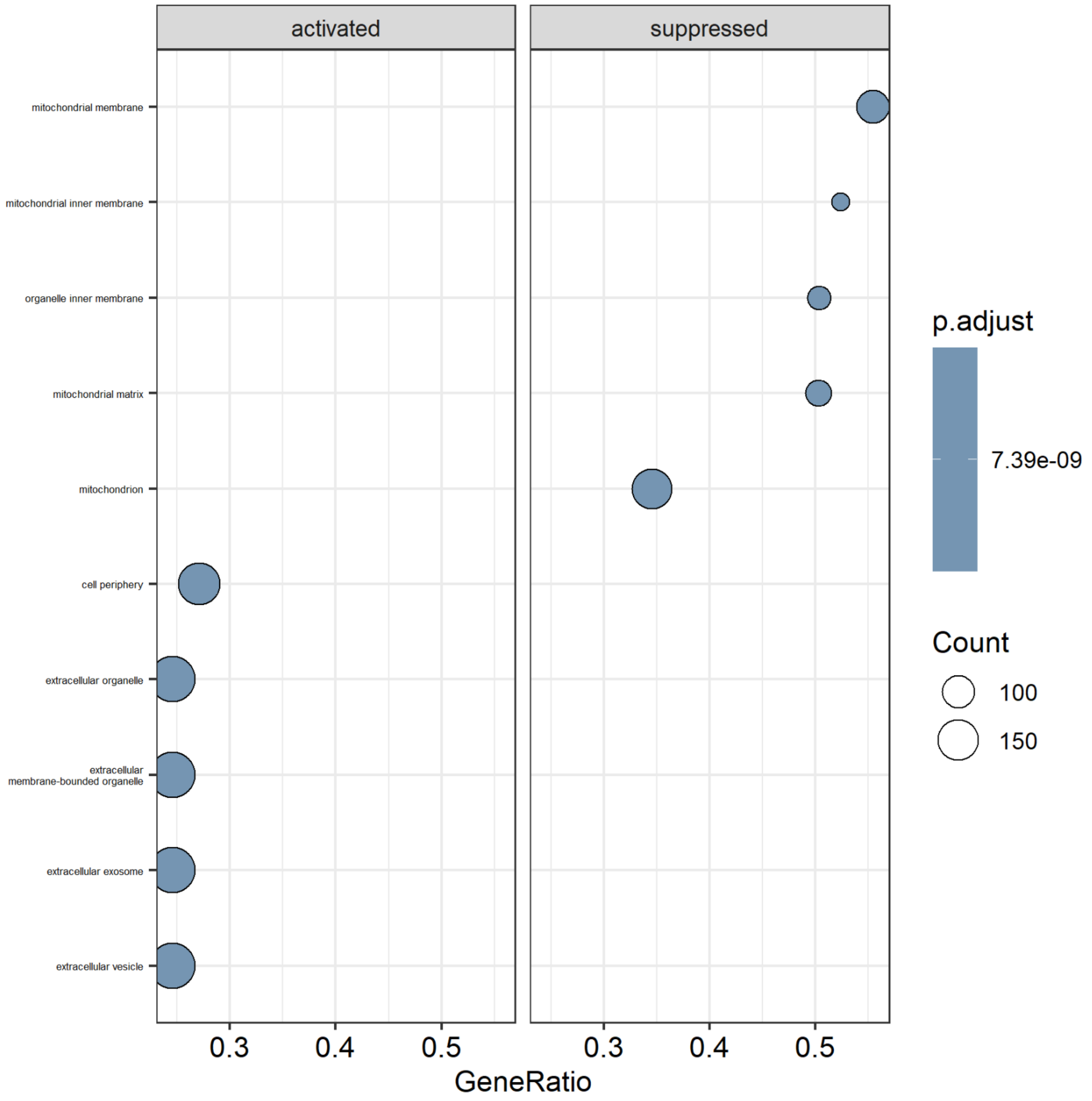

**S2 (continued).** (D) 4T vs. 4C dotplot showing activated and suppressed processes, including gene count and p-value (adjusted according to Benjamini-Hochberg) per process based on Gene Ontology Cellular Component. Based on global proteomics.

### Gene Ontology – Molecular Function

*Barplot*

**E**

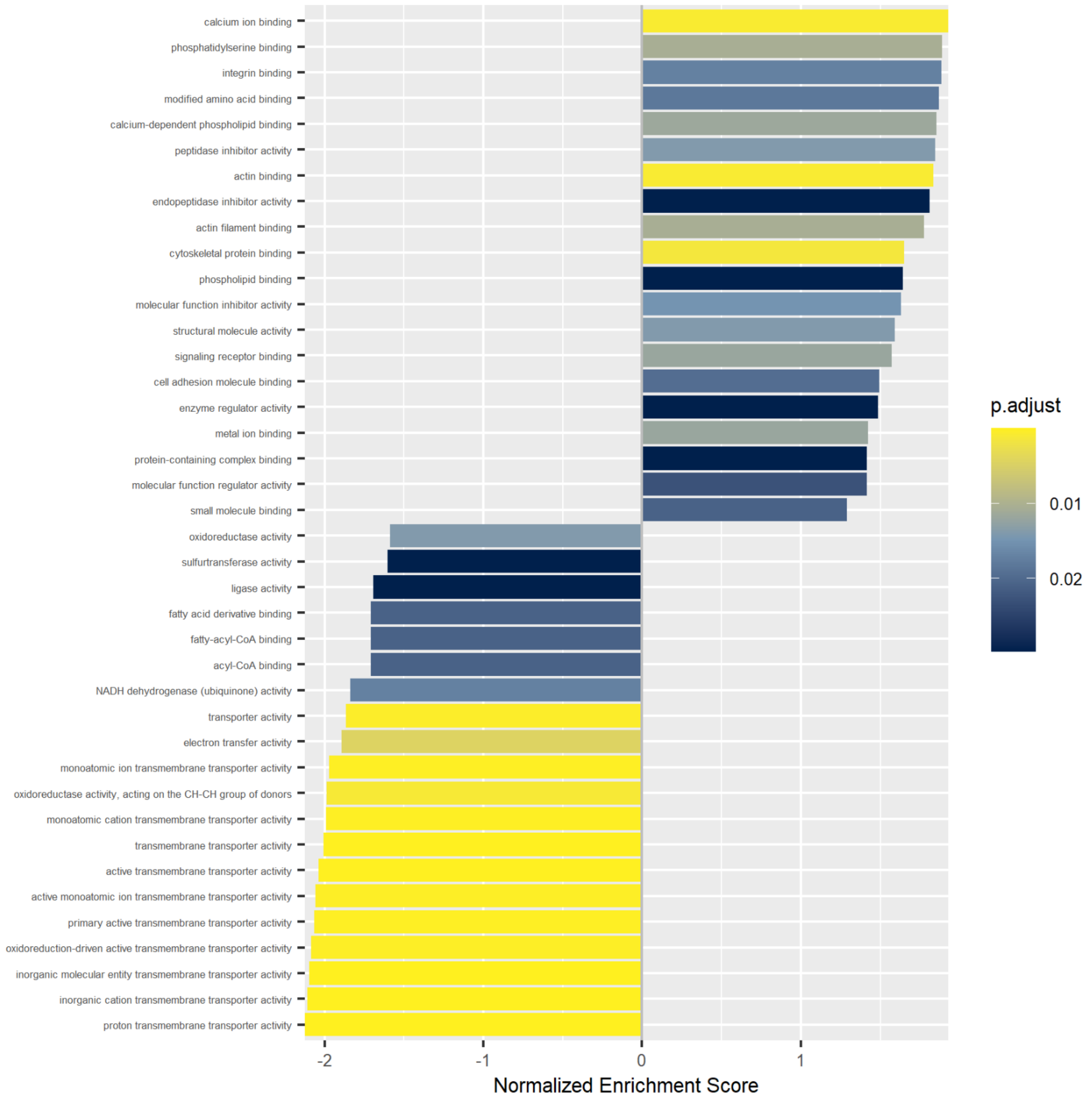

**S2 (continued).** (E) 4T vs. 4C barplot showing normalized enrichment score and p-value (adjusted according to Benjamini-Hochberg) for enriched processes based on Gene Ontology Molecular Function. Based on global proteomics.

*Dotplot*

**F**

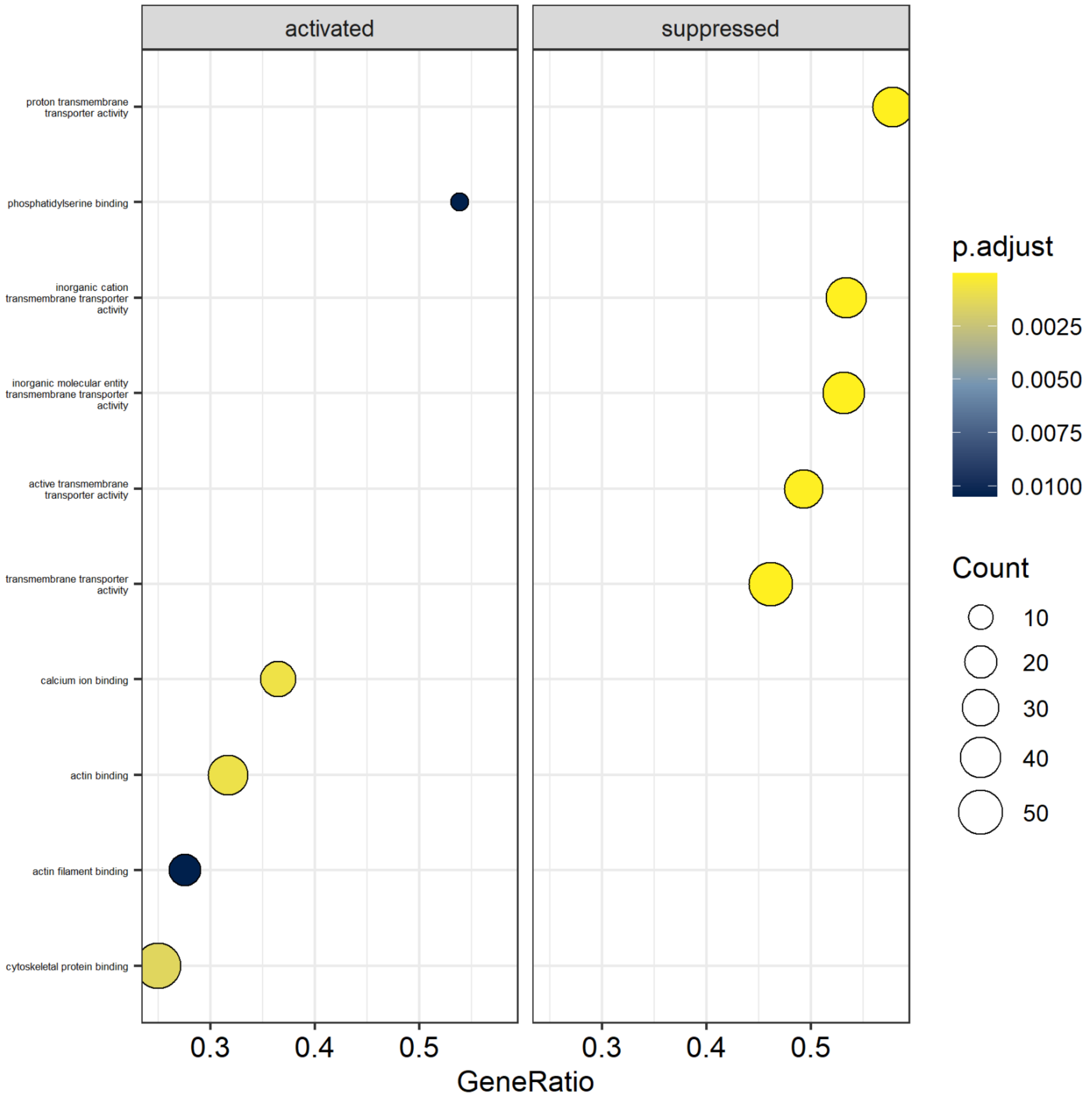

**S2 (continued).** (F) 4T vs. 4C dotplot showing activated and suppressed processes, including gene count and p-value (adjusted according to Benjamini-Hochberg) per process, based on Gene Ontology Molecular Function. Based on global proteomics.

### Reactome – Pathway Analysis

*Barplot*

**G**

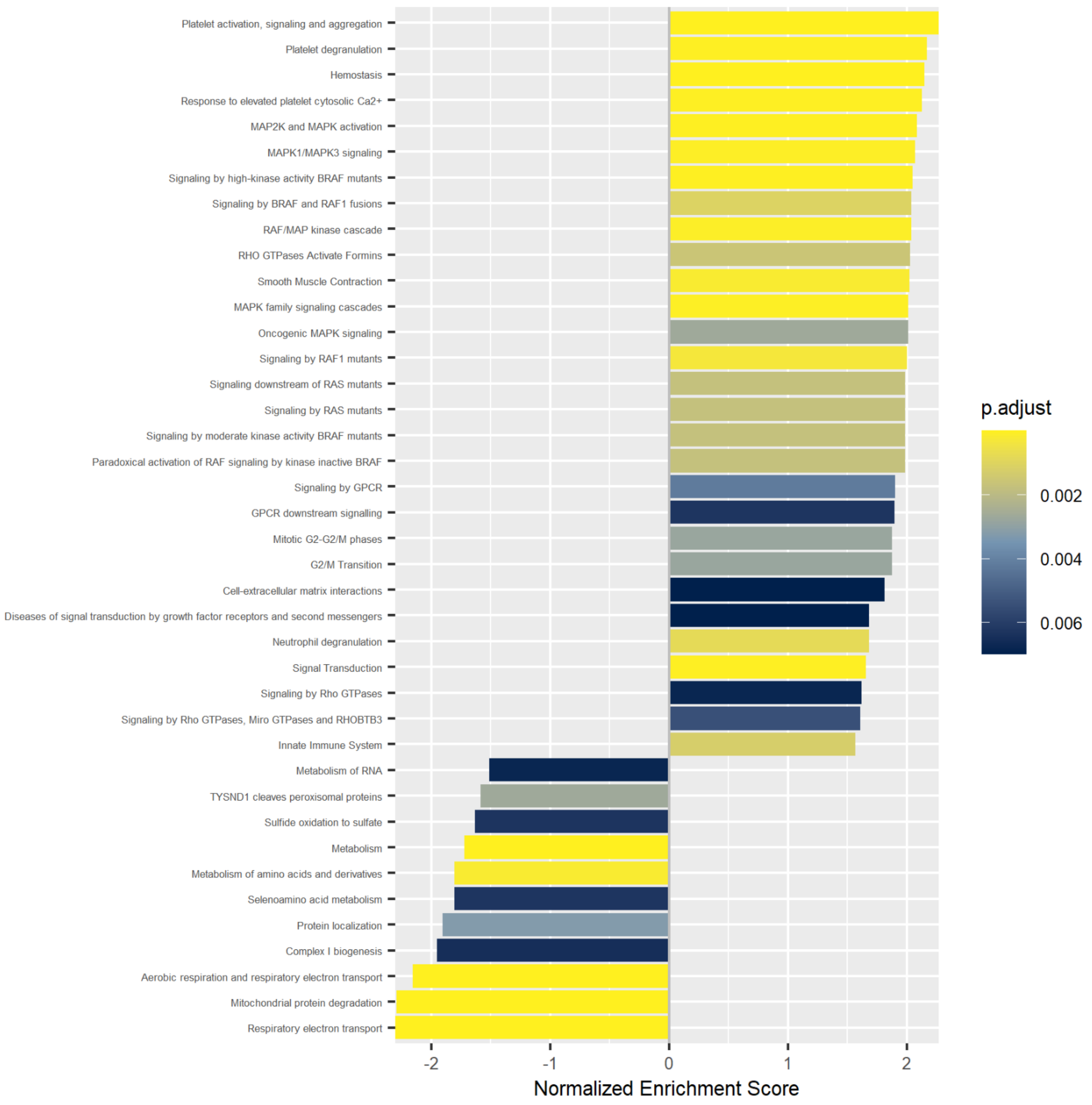

**S2 (continued).** (G) 4T vs. 4 barplot showing normalized enrichment score and p-value (adjusted according to Benjamini-Hochberg) for enriched processes based on Reactome Pathway Analysis. Based on global proteomics.

*Dotplot*

**H**

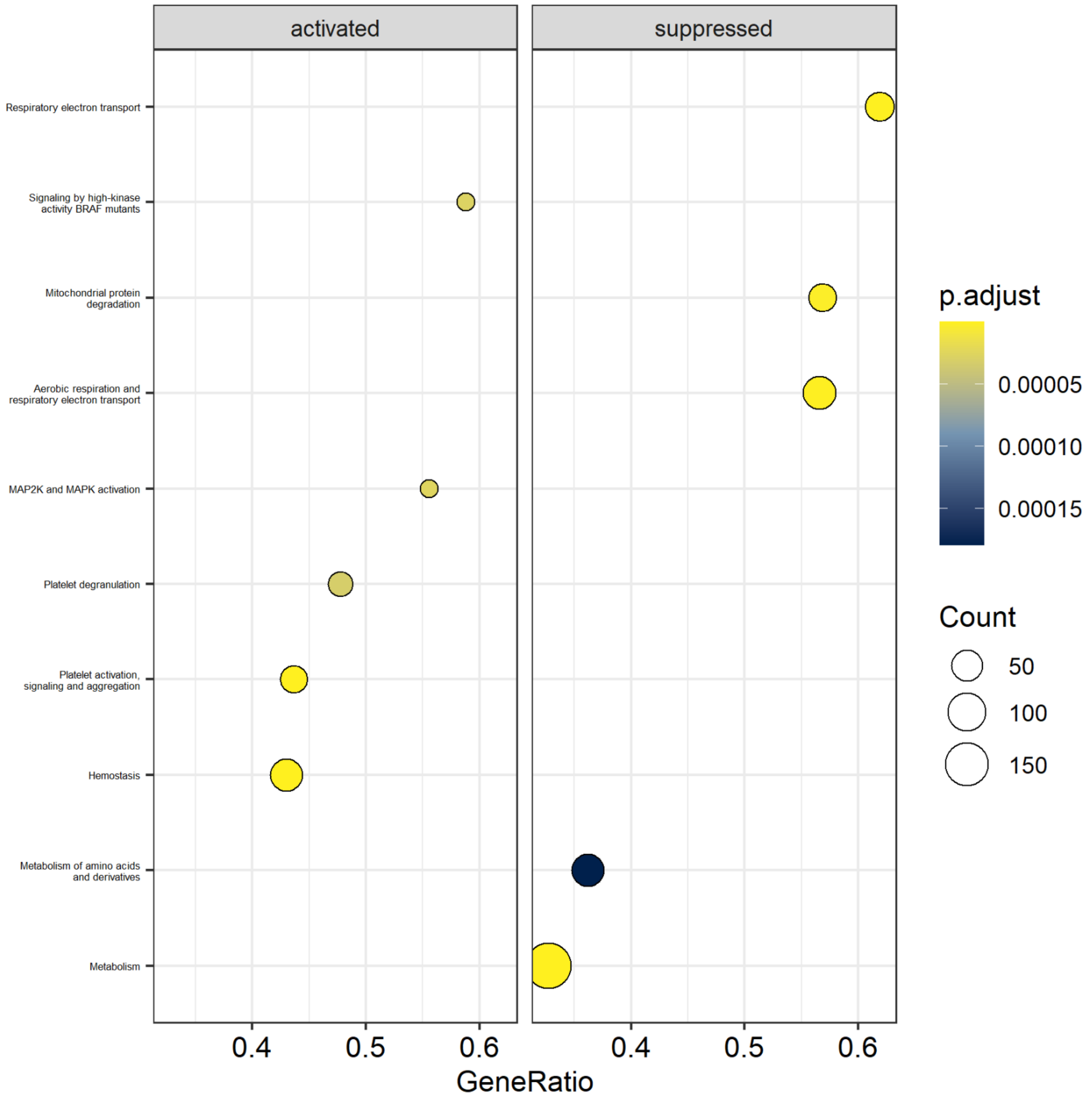

**S2 (continued).** (H) 4T vs. 4C dotplot showing activated and suppressed processes, including gene count and p-value (adjusted according to Benjamini-Hochberg) per process, based on Reactome Pathway Analysis. Based on global proteomics.

## 4AD

#### Gene Ontology - Biological Process

*Barplot*

**I**

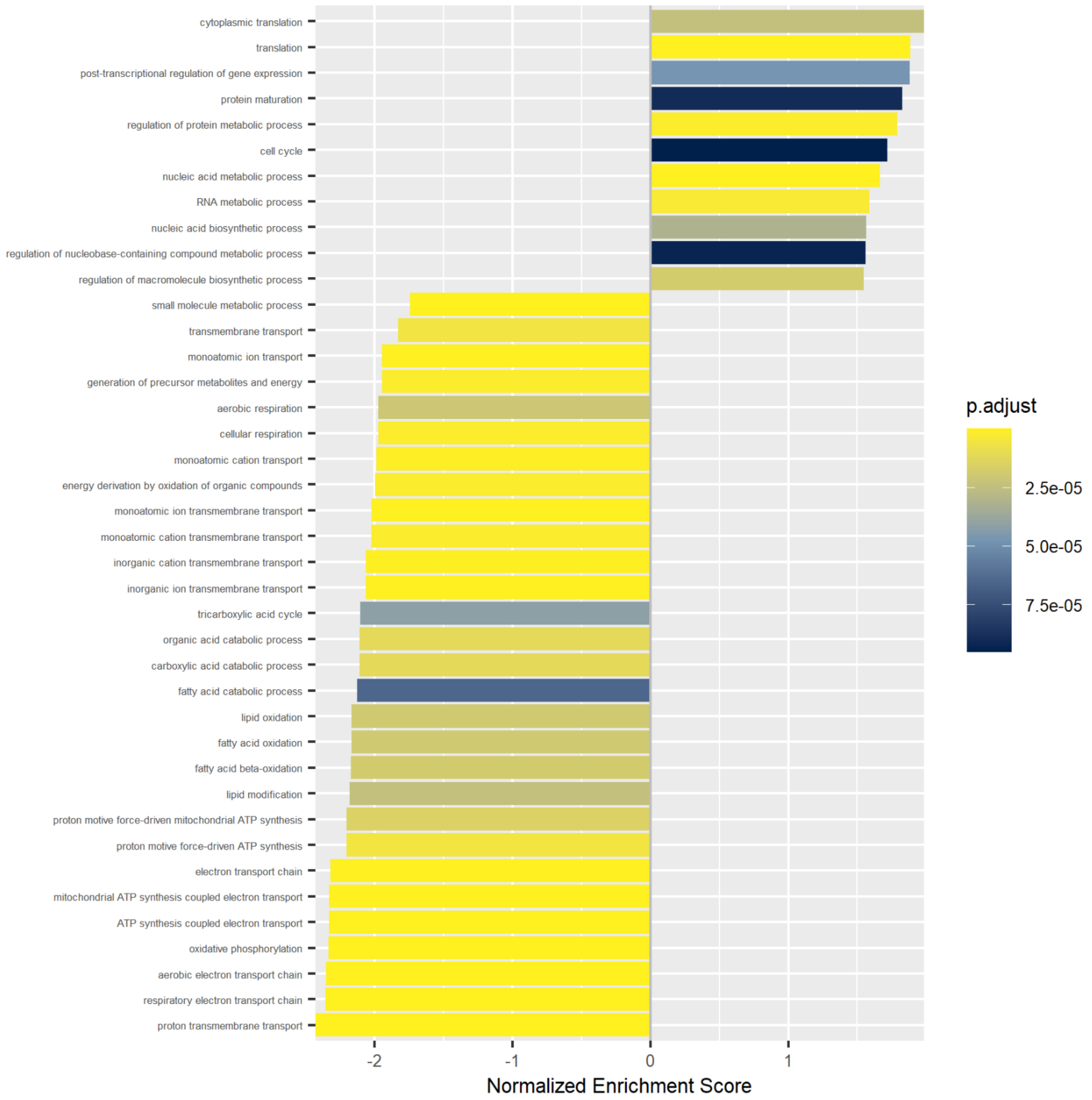

**S2 (continued).** (I) 4AD vs. 4C barplot showing normalized enrichment score and p-value (adjusted according to Benjamini-Hochberg) for enriched processes based on Gene Ontology Biological Process. Based on global proteomics.

*Dotplot*

**J**

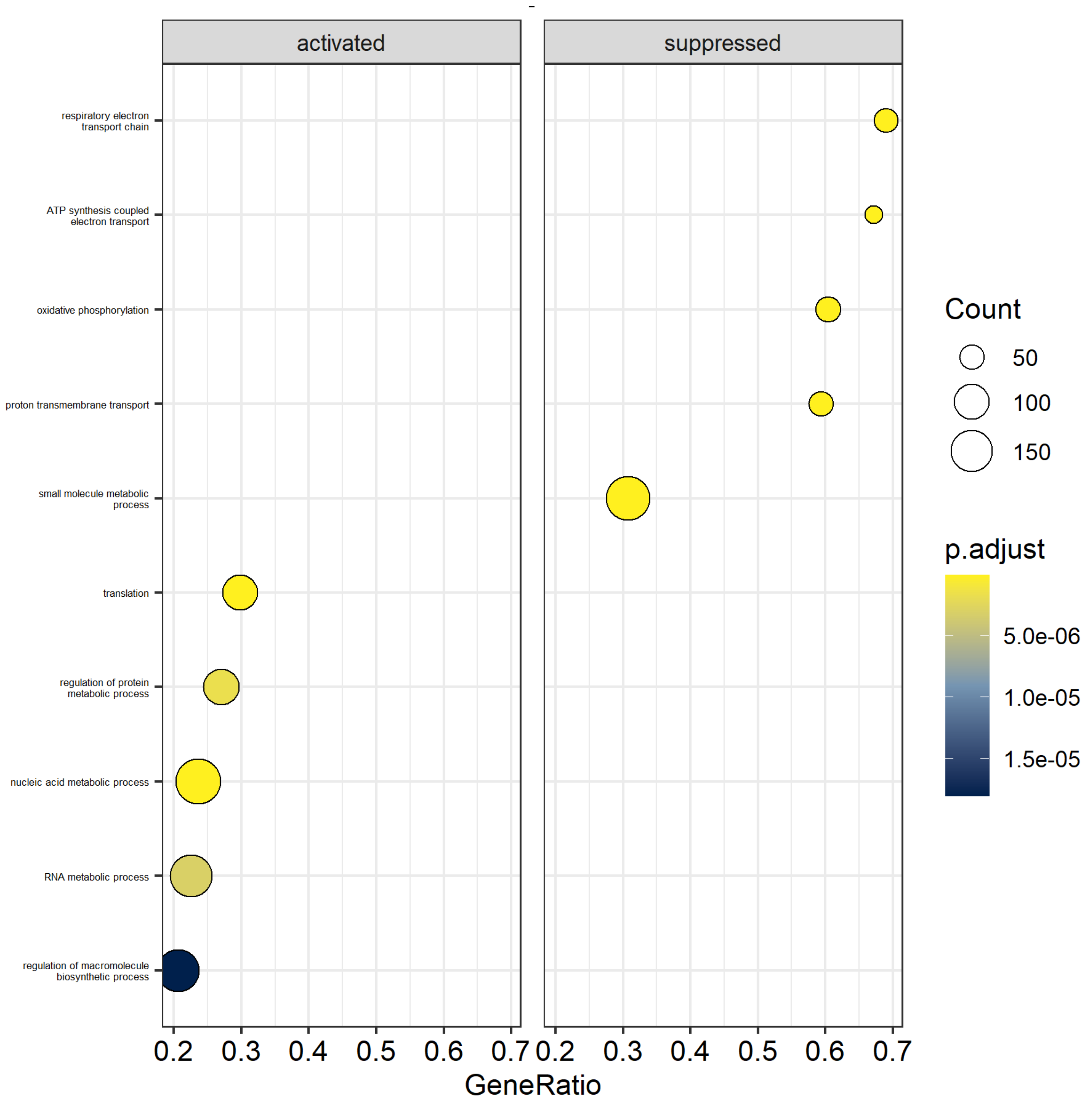

**S2 (continued).** (J) 4AD vs. 4C dotplot showing activated and suppressed processes, including gene count and p-value (adjusted according to Benjamini-Hochberg) per process, based on Gene Ontology Biological Process. Based on global proteomics.

### Gene Ontology – Cellular Component

*Barplot*

**K**

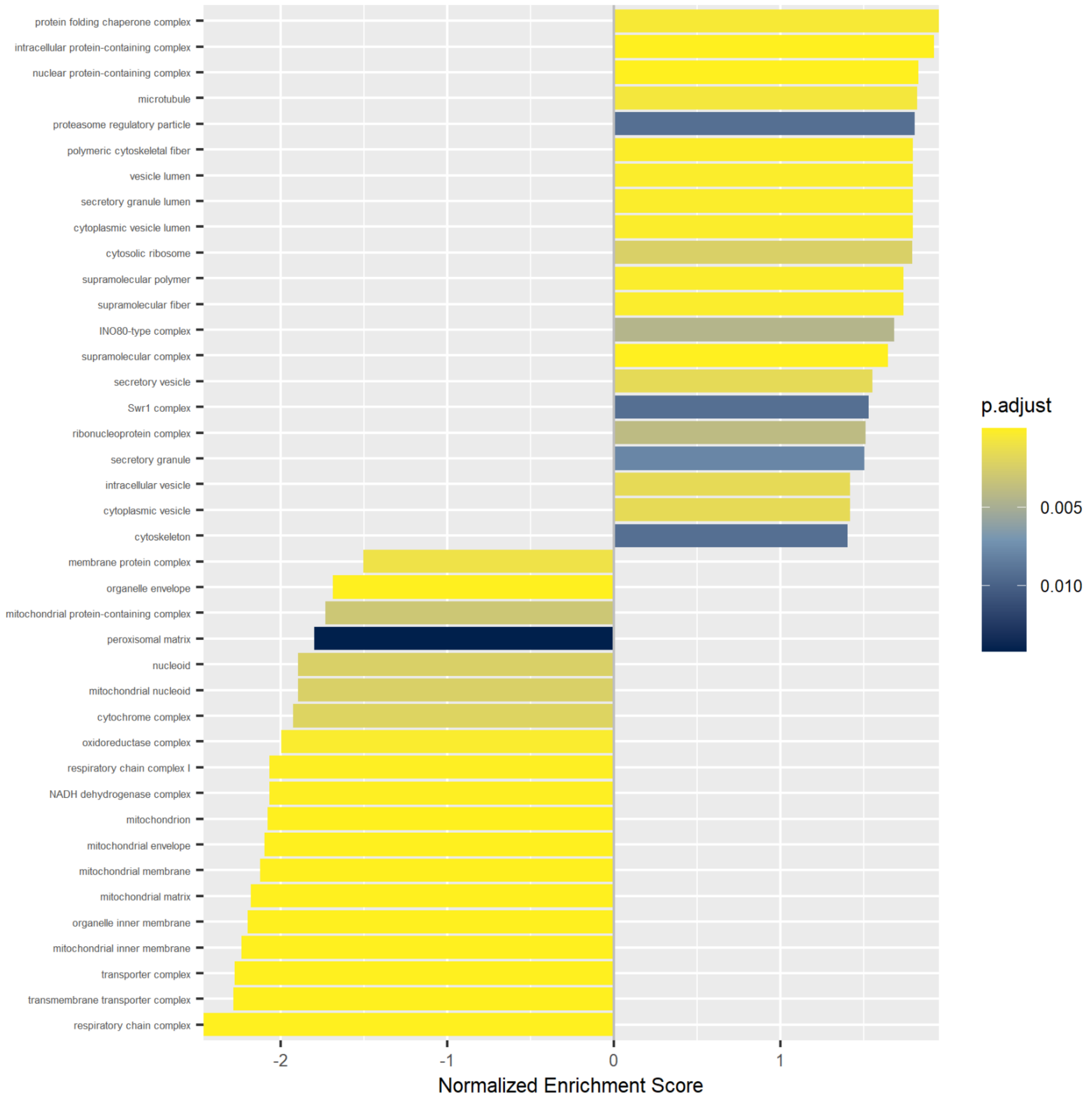

**S2 (continued).** (K) 4AD vs. 4C barplot showing normalized enrichment score and p-value (adjusted according to Benjamini-Hochberg) for enriched processes based on Gene Ontology Cellular Component. Based on global proteomics.

*Dotplot*

**L**

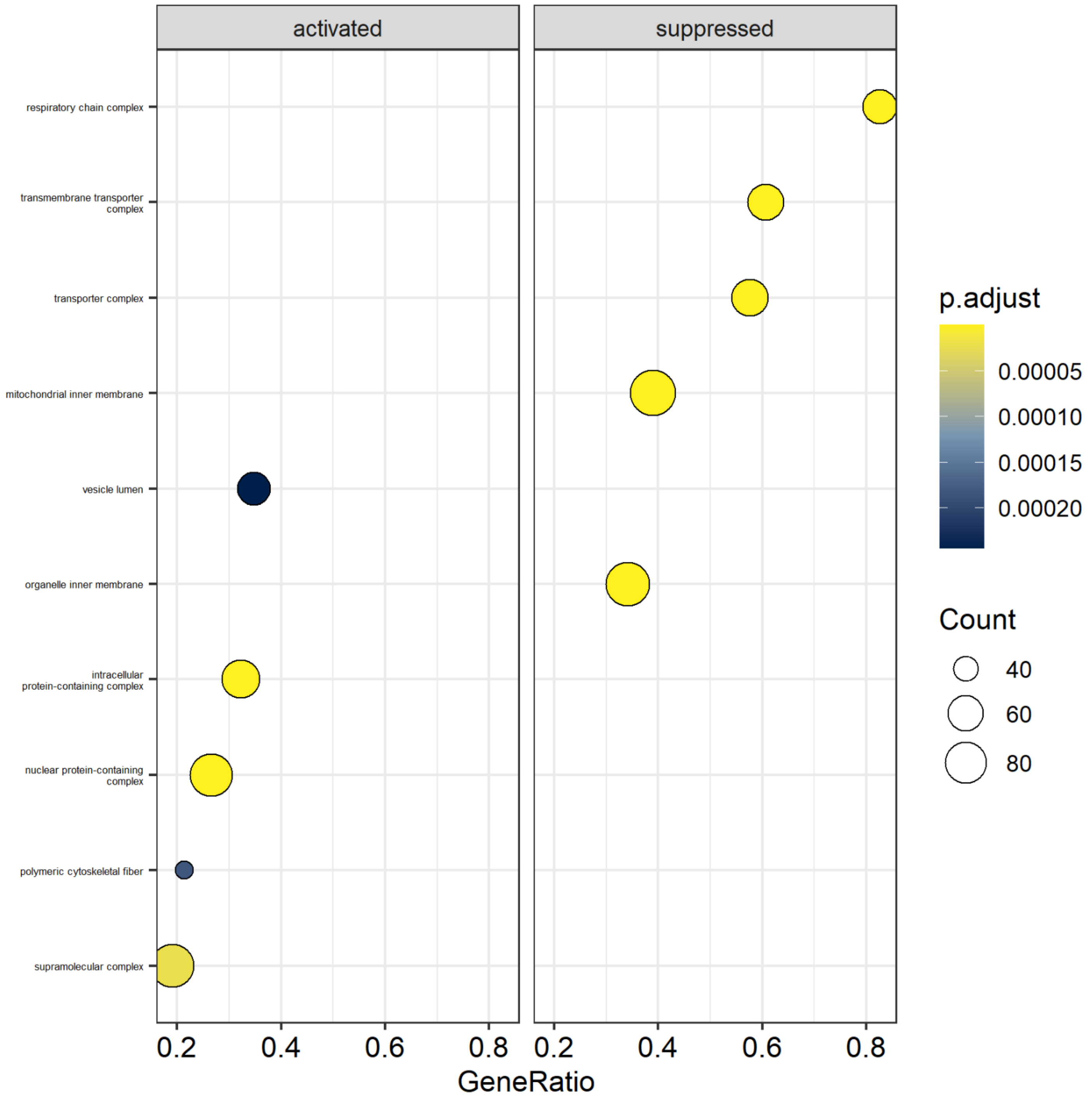

**S2 (continued).** (L) 4AD vs. 4C dotplot showing activated and suppressed processes, including gene count and p-value (adjusted according to Benjamini-Hochberg) per process based on Gene Ontology Cellular Component. Based on global proteomics.

### Gene Ontology – Molecular Function

*Barplot*

**M**

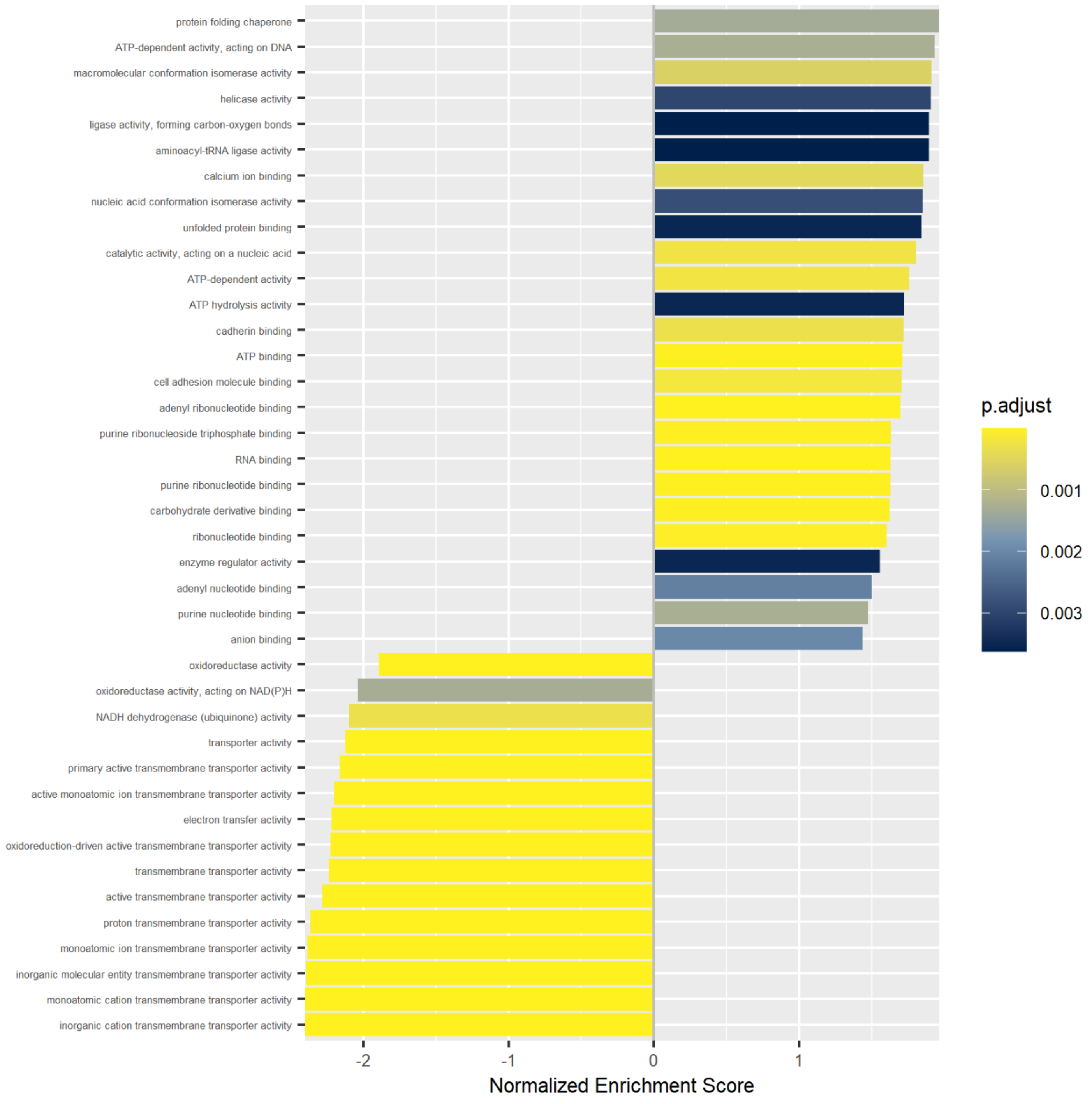

**S2 (continued).** (M) 4AD vs. 4C barplot showing normalized enrichment score and p-value (adjusted according to Benjamini-Hochberg) for enriched processes based on Gene Ontology Molecular Function. Based on global proteomics.

*Dotplot*

**N**

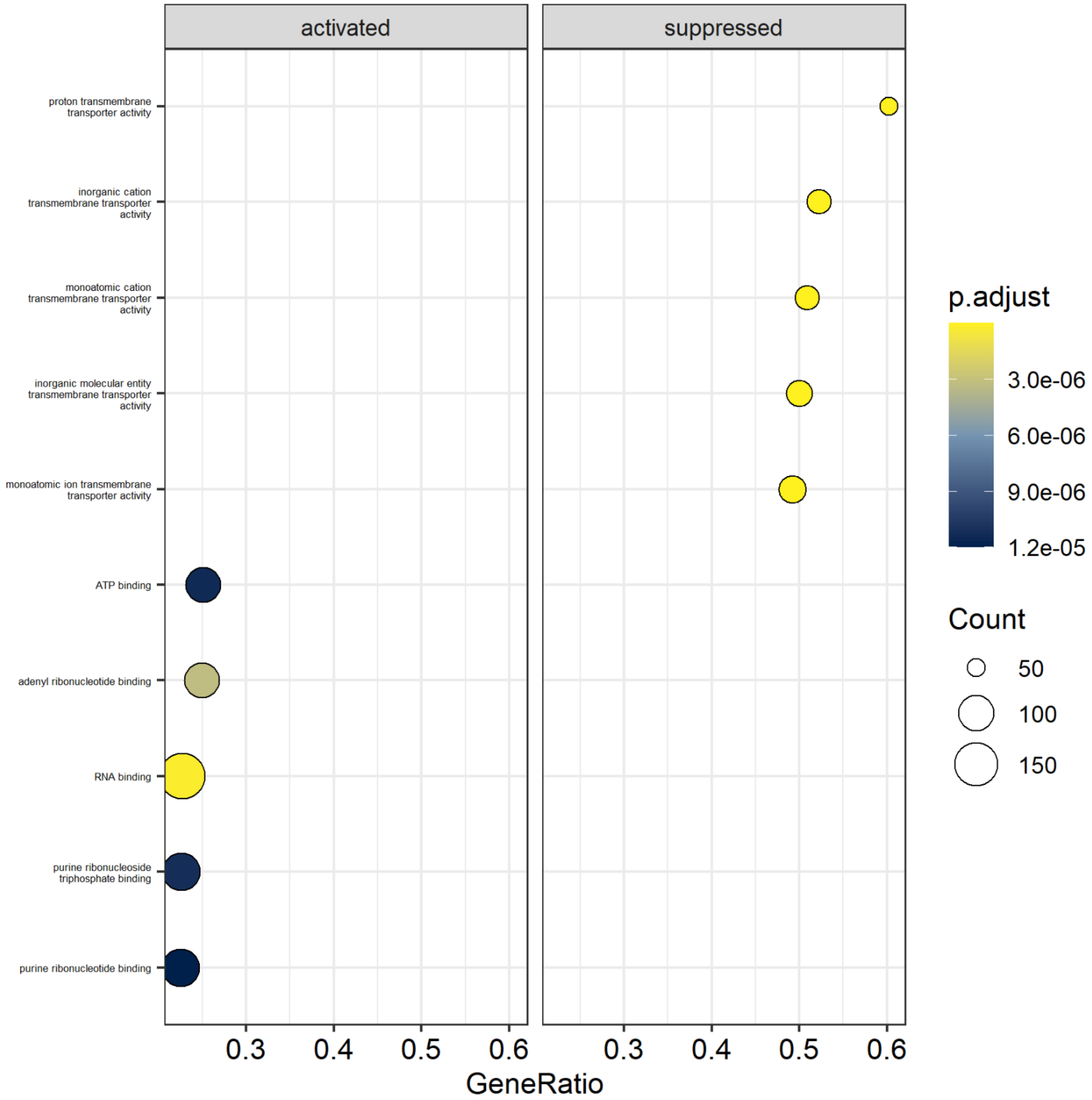

**S2 (continued).** (N) 4AD vs. 4C dotplot showing activated and suppressed processes, including gene count and p-value (adjusted according to Benjamini-Hochberg) per process based on Gene Ontology Molecular Function. Based on global proteomics.

### Reactome – Pathway Analysis

#### *Barplot*

**O**

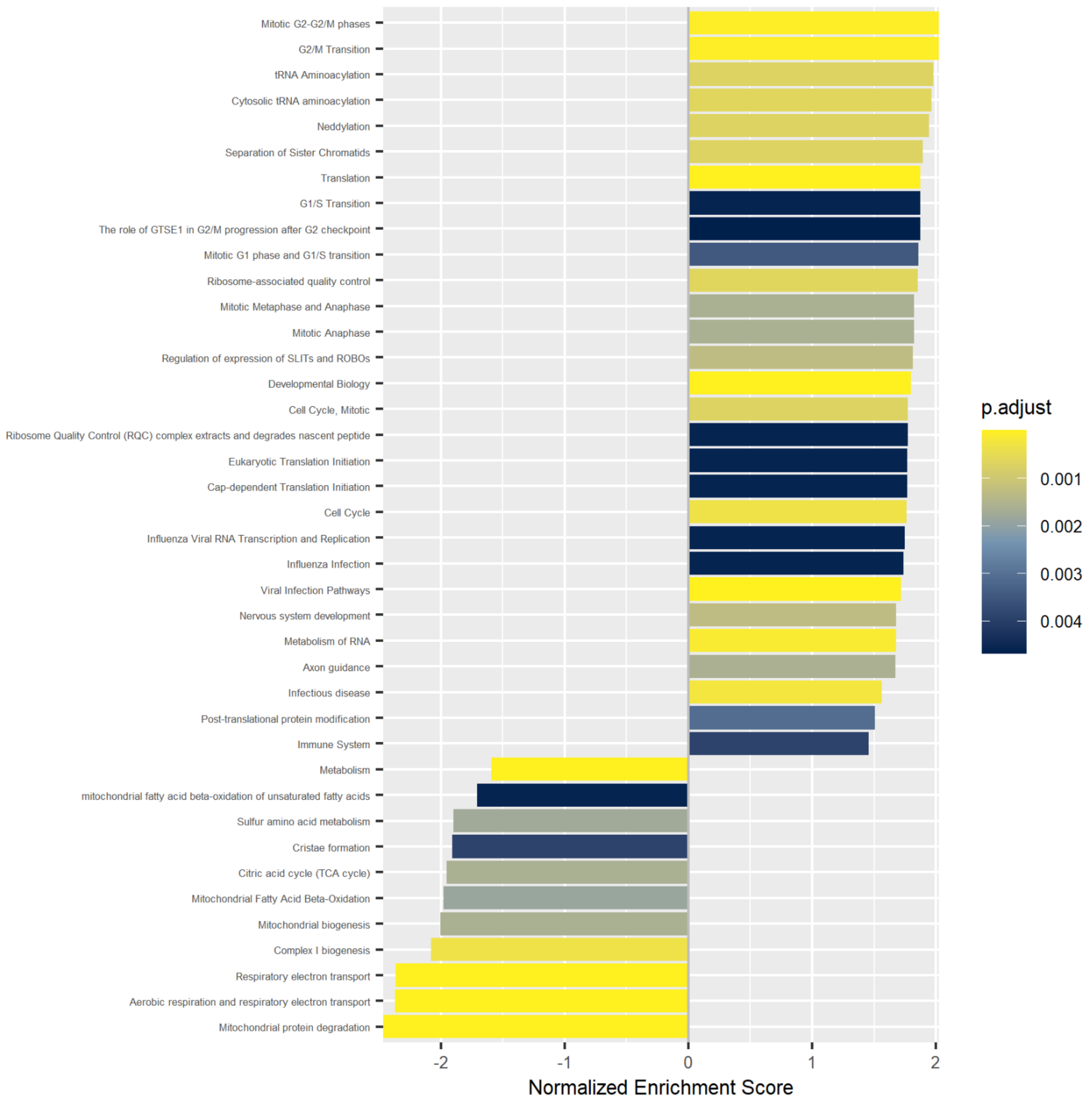

**S2 (continued).** (O) 4AD vs. 4C barplot showing normalized enrichment score and p-value (adjusted according to Benjamini-Hochberg) for enriched processes based on Reactome Pathway Analysis. Based on global proteomics.

*Dotplot*

**P**

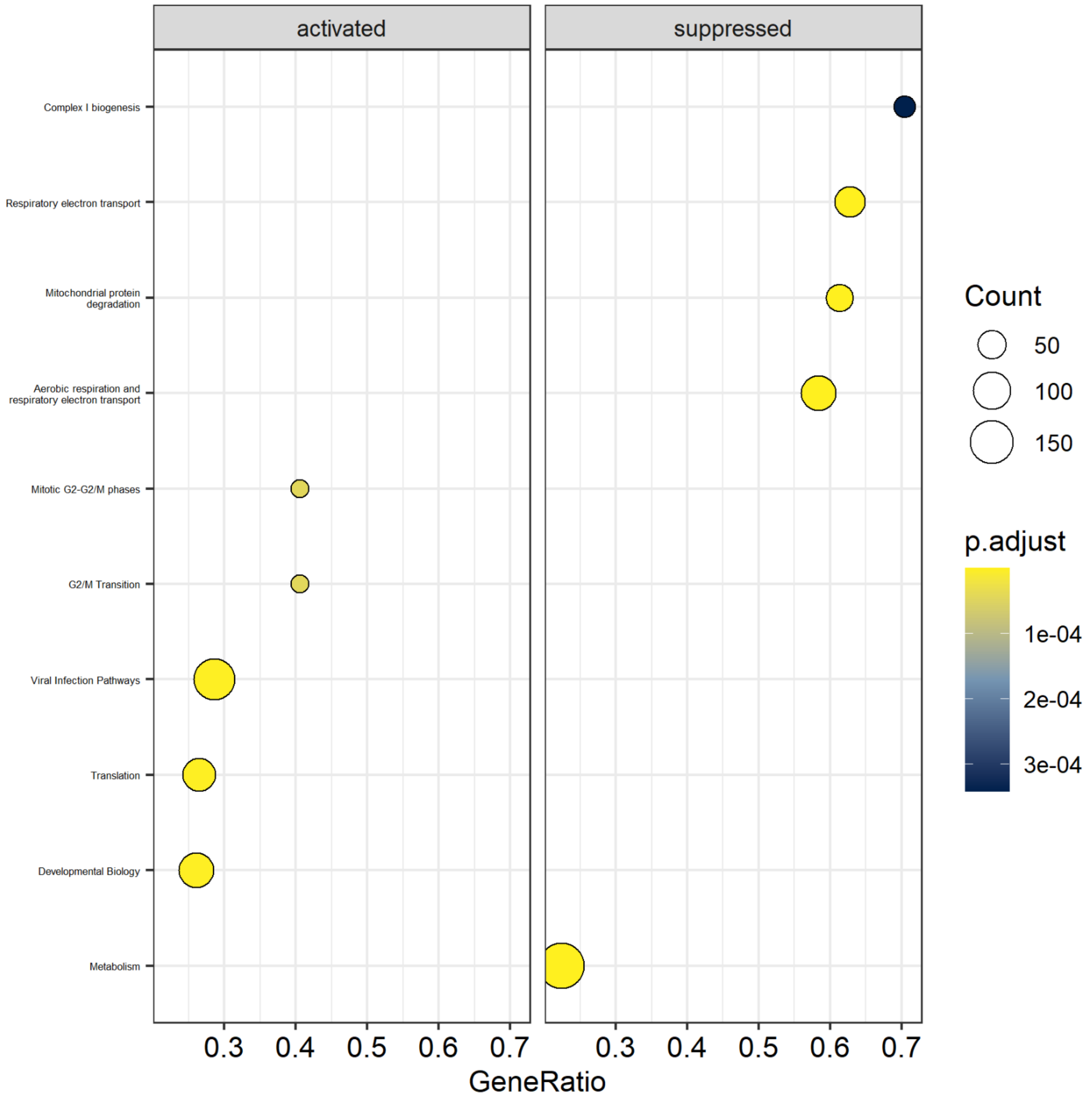

**S2 (continued).** (P) 4AD vs. 4C dotplot showing activated and suppressed processes, including gene count and p-value (adjusted according to Benjamini-Hochberg) per process, based on Reactome Pathway Analysis. Based on global proteomics.

# 4BD

### Gene Ontology - Biological Process

*Barplot*

**Q**

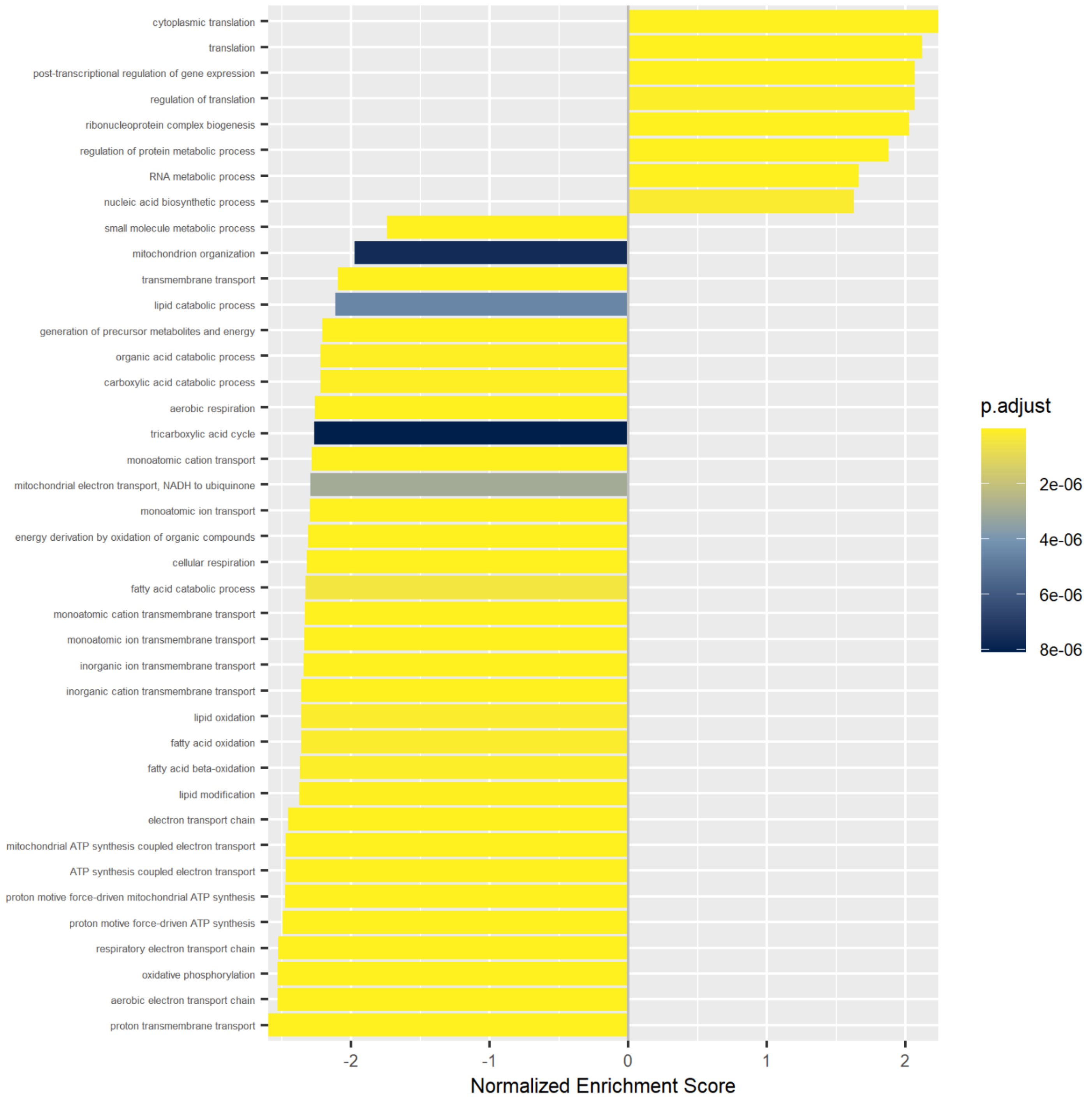

**S2 (continued).** (Q) 4BD vs. 4C barplot showing normalized enrichment score and p-value (adjusted according to Benjamini-Hochberg) for enriched processes based on Gene Ontology Biological Process. Based on global proteomics.

*Dotplot*

**R**

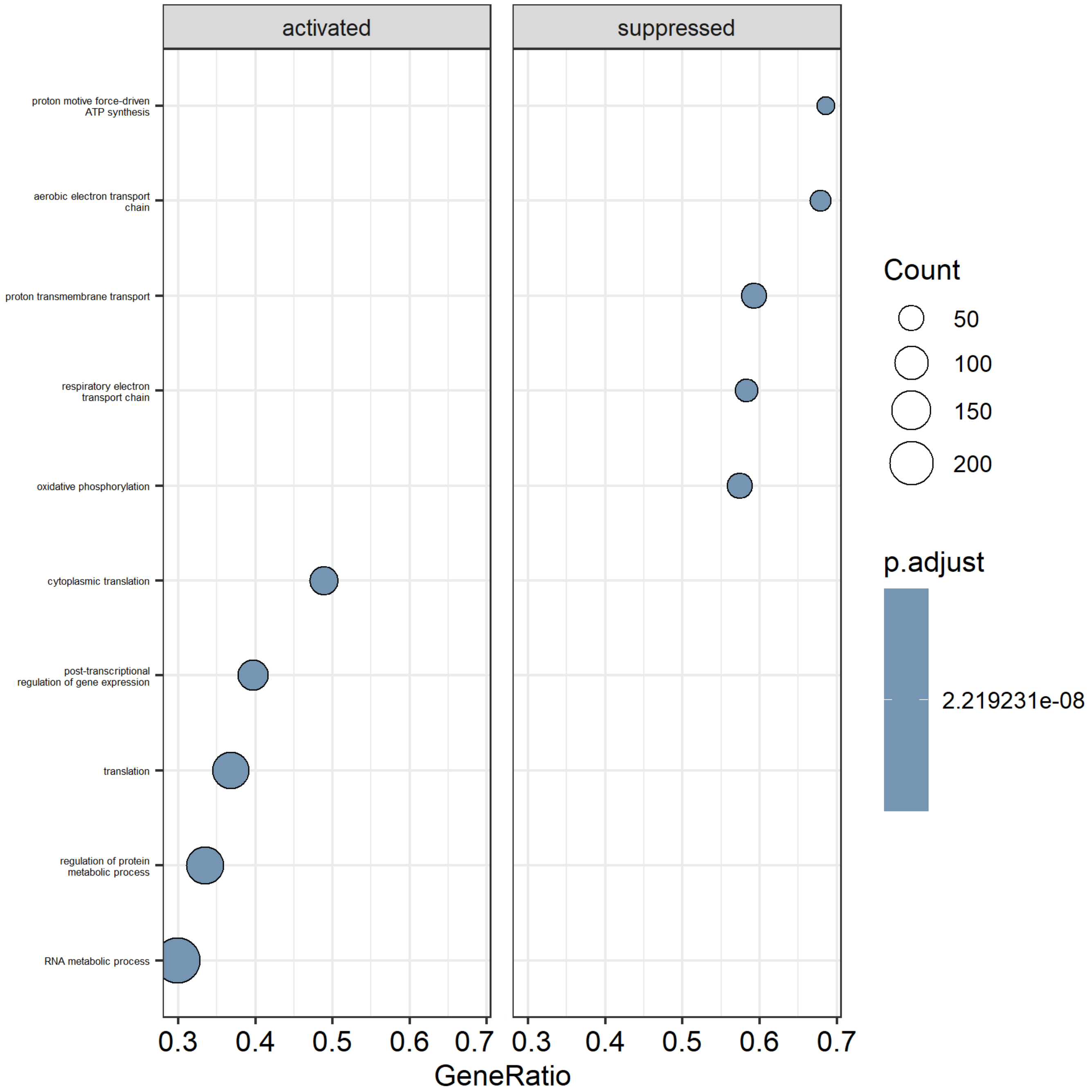

**S2 (continued).** (R) 4BD vs. 4C dotplot showing activated and suppressed processes, including gene count and p-value (adjusted according to Benjamini-Hochberg) per process based on Gene Ontology Biological Process. Based on global proteomics.

### Gene Ontology – Cellular Component

*Barplot*

**S**

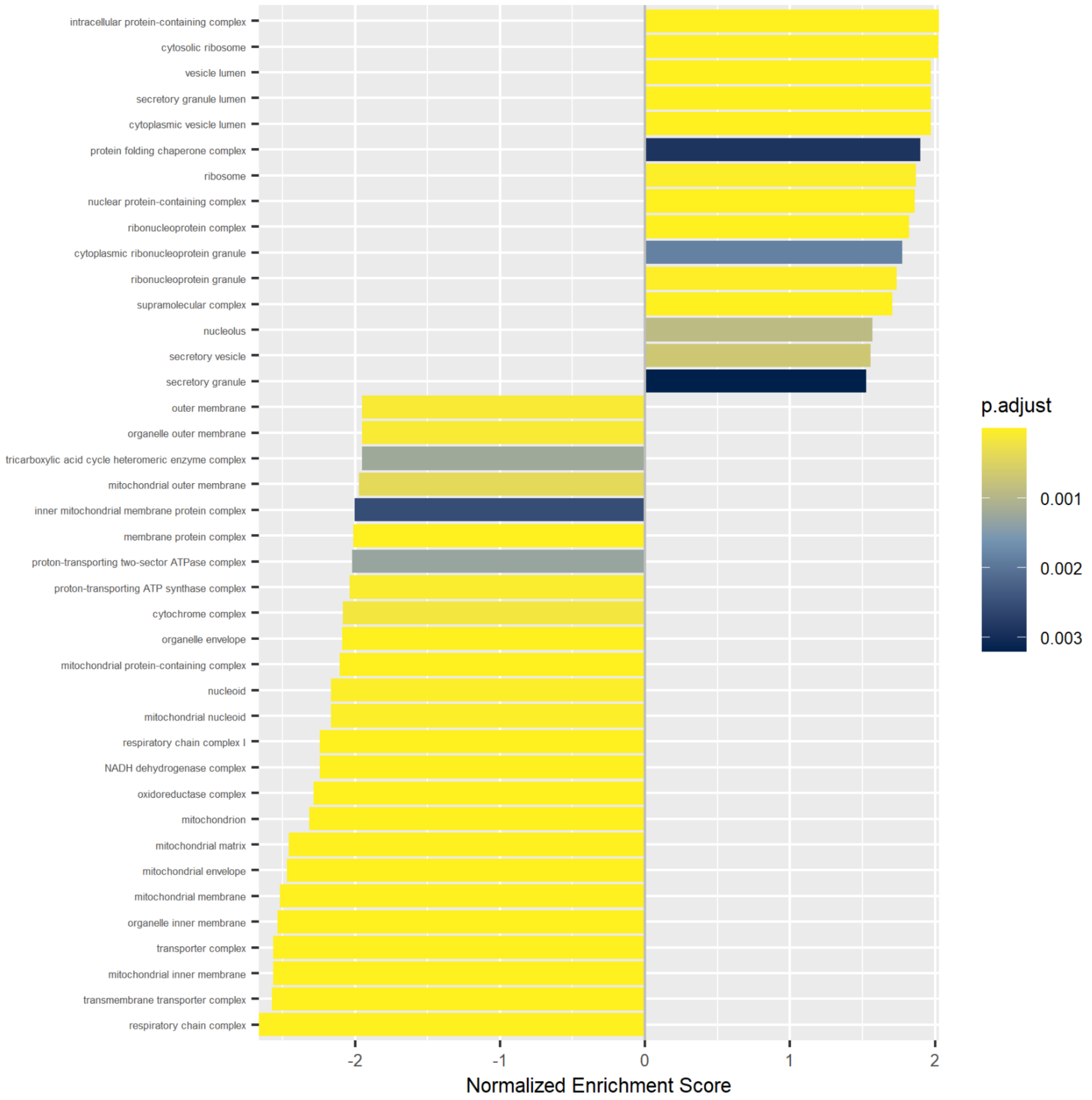

**S2 (continued).** (S) 4BD vs. 4C barplot showing normalized enrichment score and p-value (adjusted according to Benjamini-Hochberg) for enriched processes based on Gene Ontology Cellular Component. Based on global proteomics.

*Dotplot*

**T**

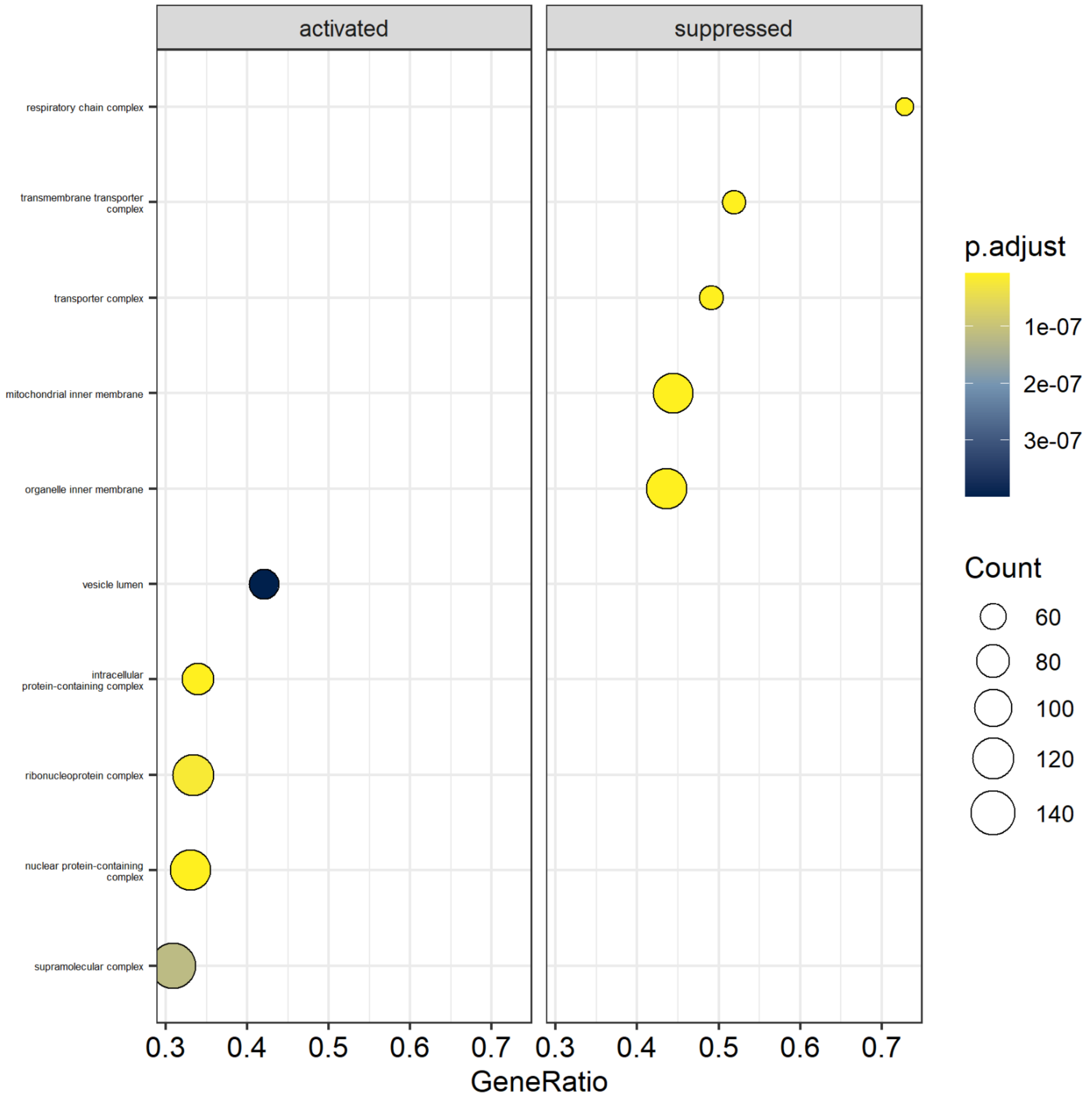

**S2 (continued).** (T) 4BD vs. 4C dotplot showing activated and suppressed processes, including gene count and p-value (adjusted according to Benjamini-Hochberg) per process based on Gene Ontology Cellular Component. Based on global proteomics.

### Gene Ontology – Molecular Function

*Barplot*

**U**

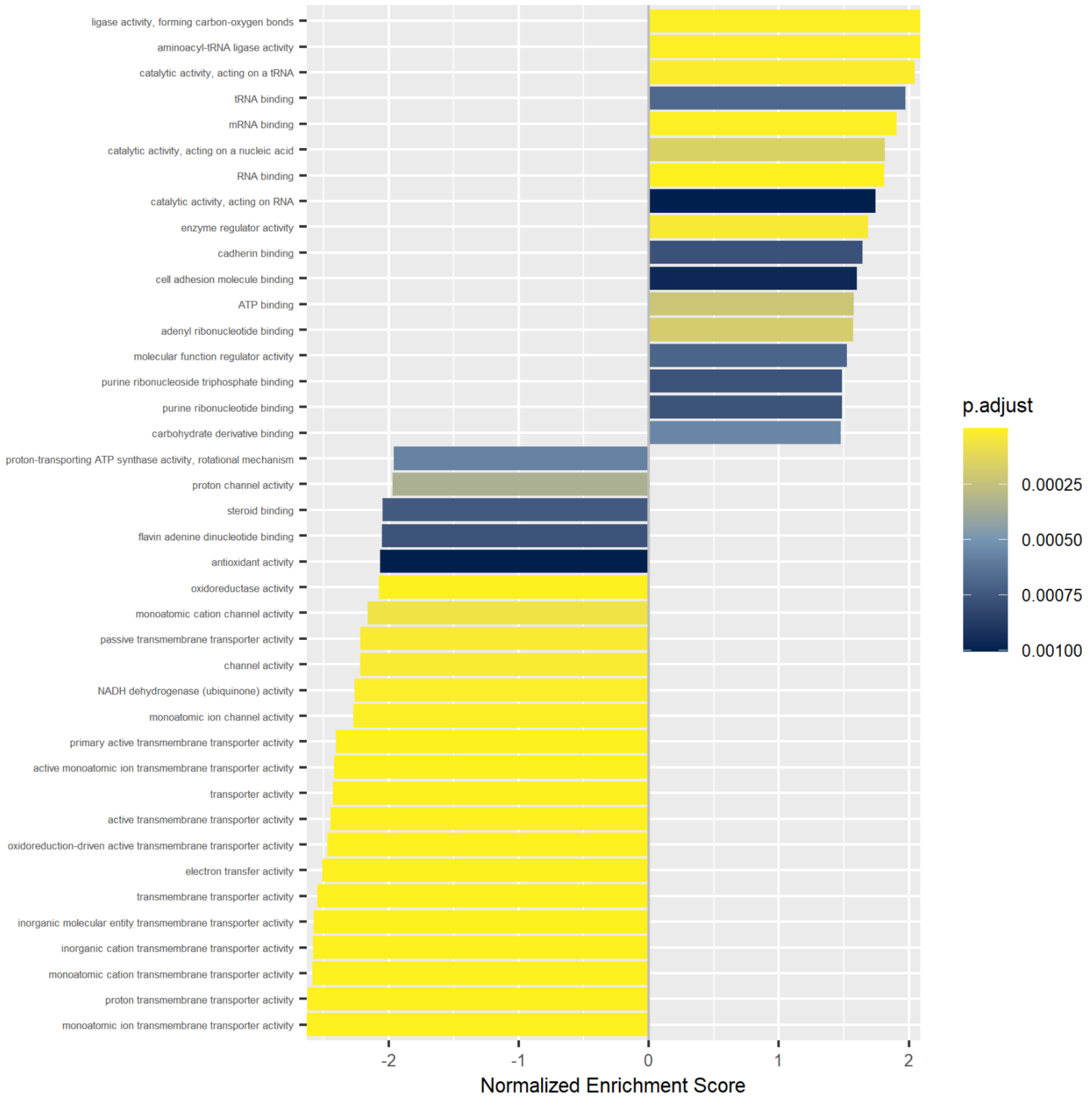

**S2 (continued).** (U) 4BD vs. 4C Barplot showing normalized enrichment score and p-value (adjusted according to Benjamini-Hochberg) for enriched processes based on Gene Ontology Molecular Function. Based on global proteomics.

*Dotplot*

**V**

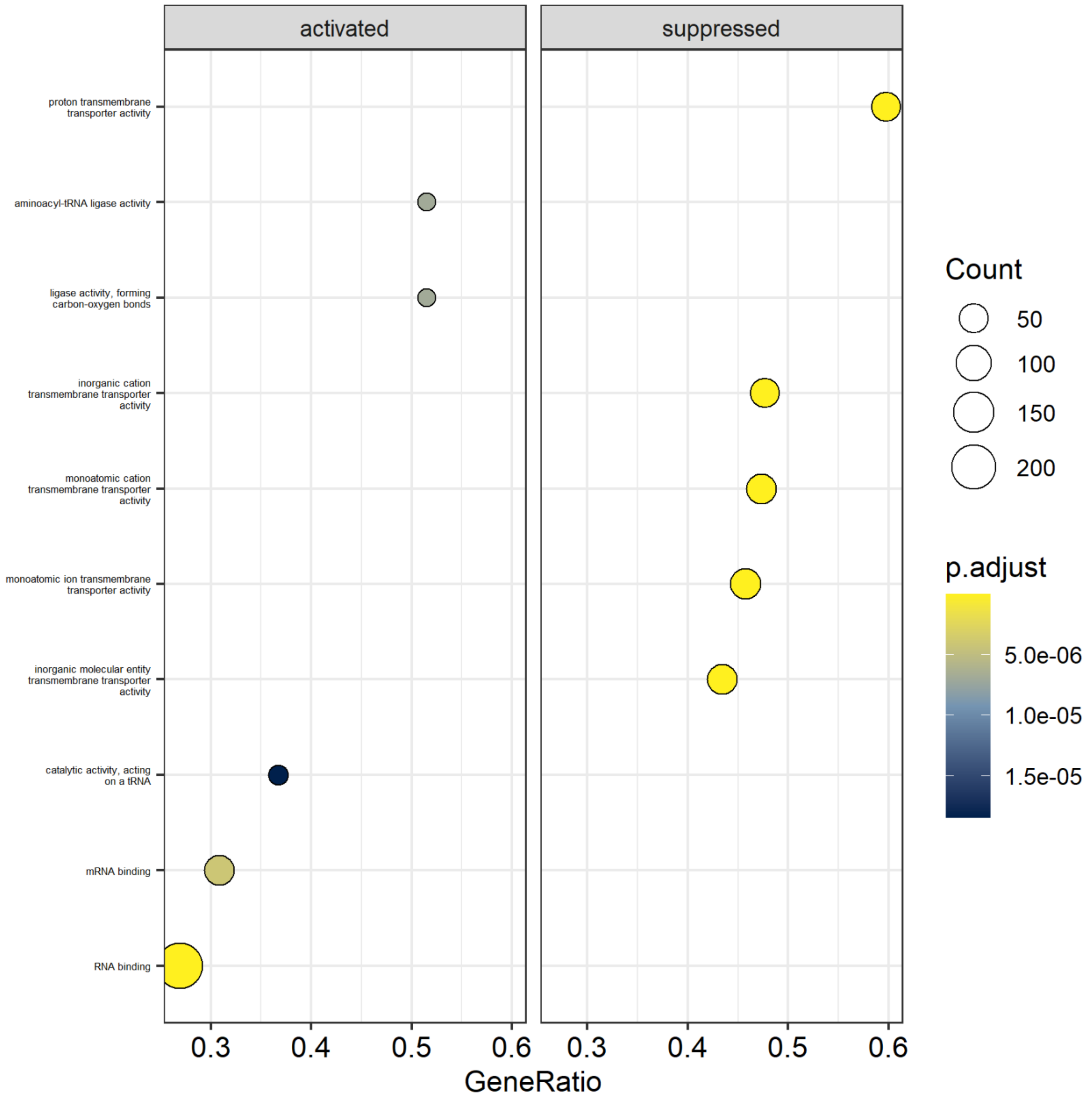

**S2 (continued).** (V) 4BD vs. 4C dotplot showing activated and suppressed processes, including gene count and p-value (adjusted according to Benjamini-Hochberg) per process, based on Gene Ontology Molecular Function. Based on global proteomics.

### Reactome – Pathway Analysis

*Barplot*

**W**

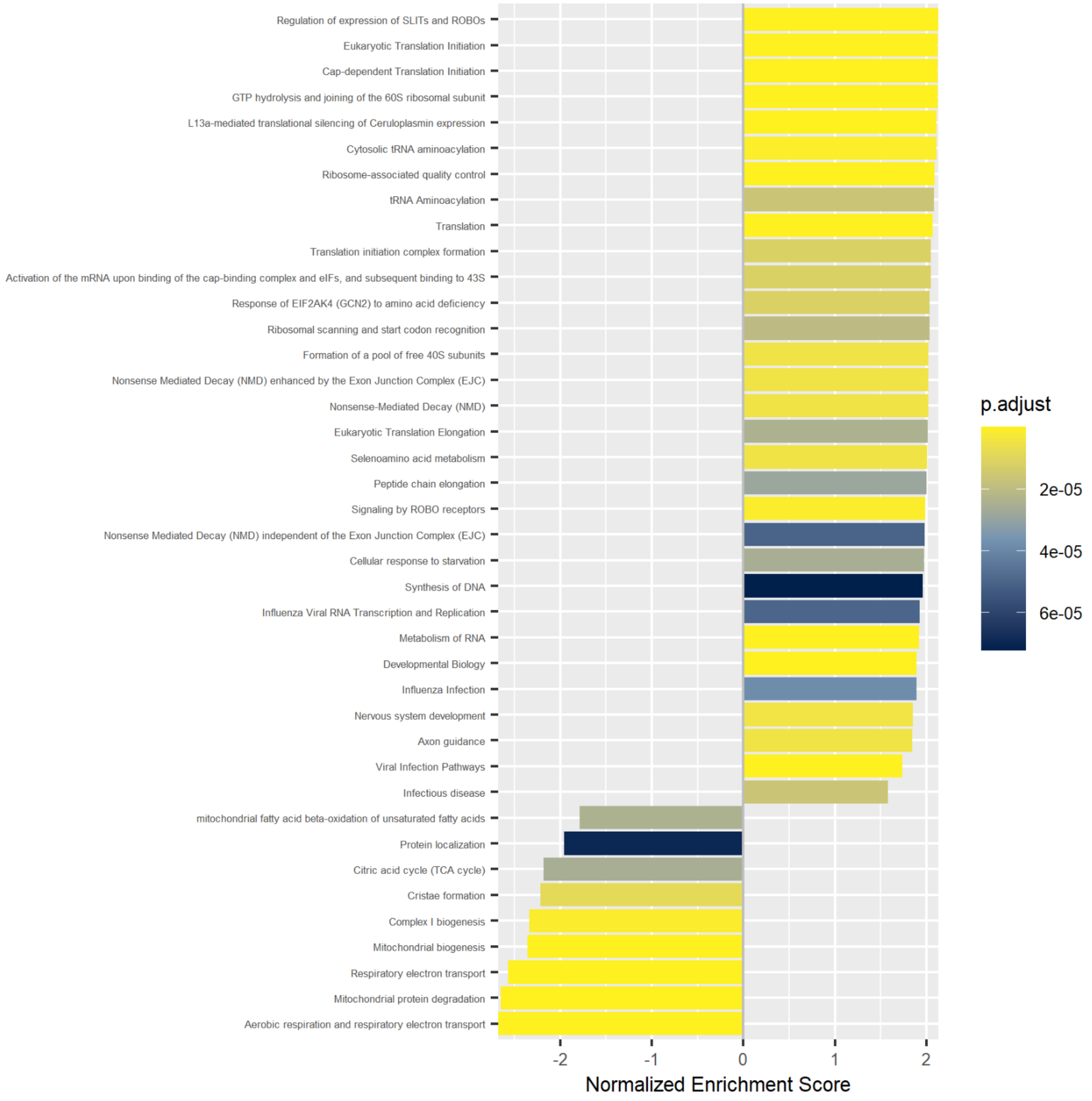

**S2 (continued).** (W) 4BD vs. 4C barplot showing normalized enrichment score and p-value (adjusted according to Benjamini-Hochberg) for enriched processes based on Reactome Pathway Analysis. Based on global proteomics.

*Dotplot*

**X**

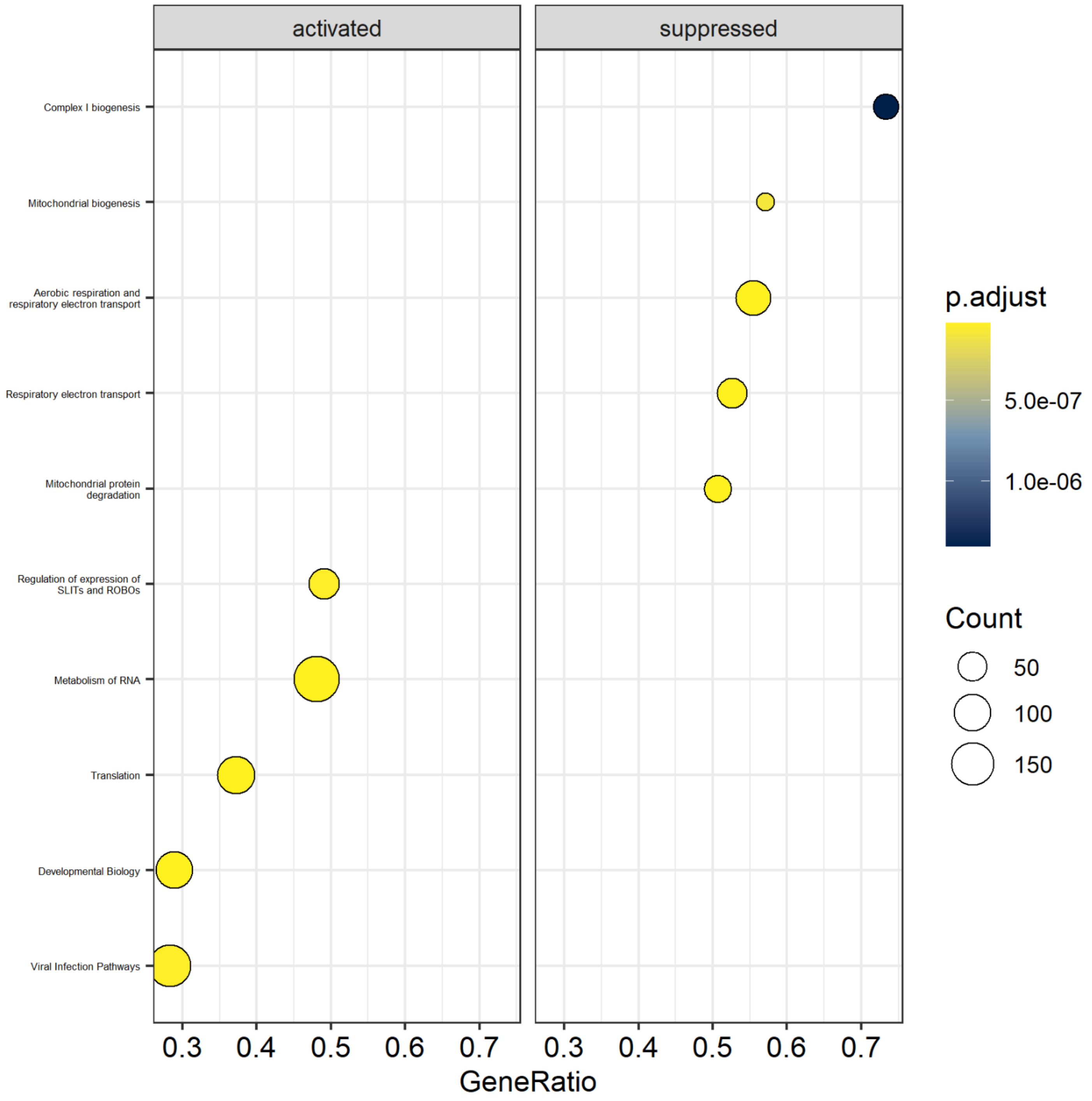

**S2 (continued).** (X) 4BD vs. 4C dotplot showing activated and suppressed processes, including gene count and p-value (adjusted according to Benjamini-Hochberg) per process, based on Reactome Pathway Analysis. Based on global proteomics.

## 4AG

#### Gene Ontology - Biological Process

*Barplot*

**Y**

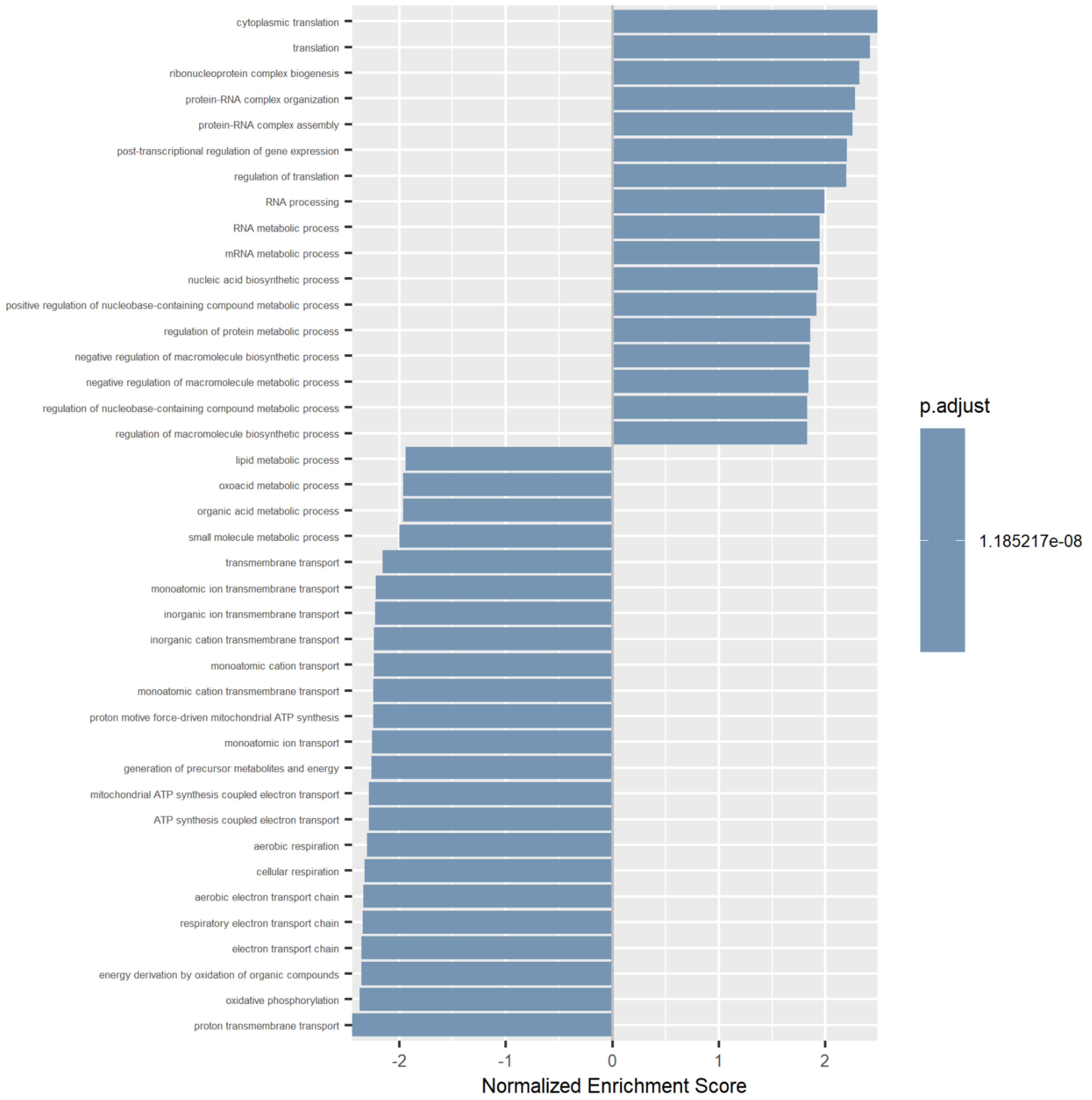

**S2 (continued).** (Y) 4AG vs. 4C barplot showing normalized enrichment score and p-value (adjusted according to Benjamini-Hochberg) for enriched processes based on Gene Ontology Biological Process. Based on global proteomics.

*Dotplot*

**Z**

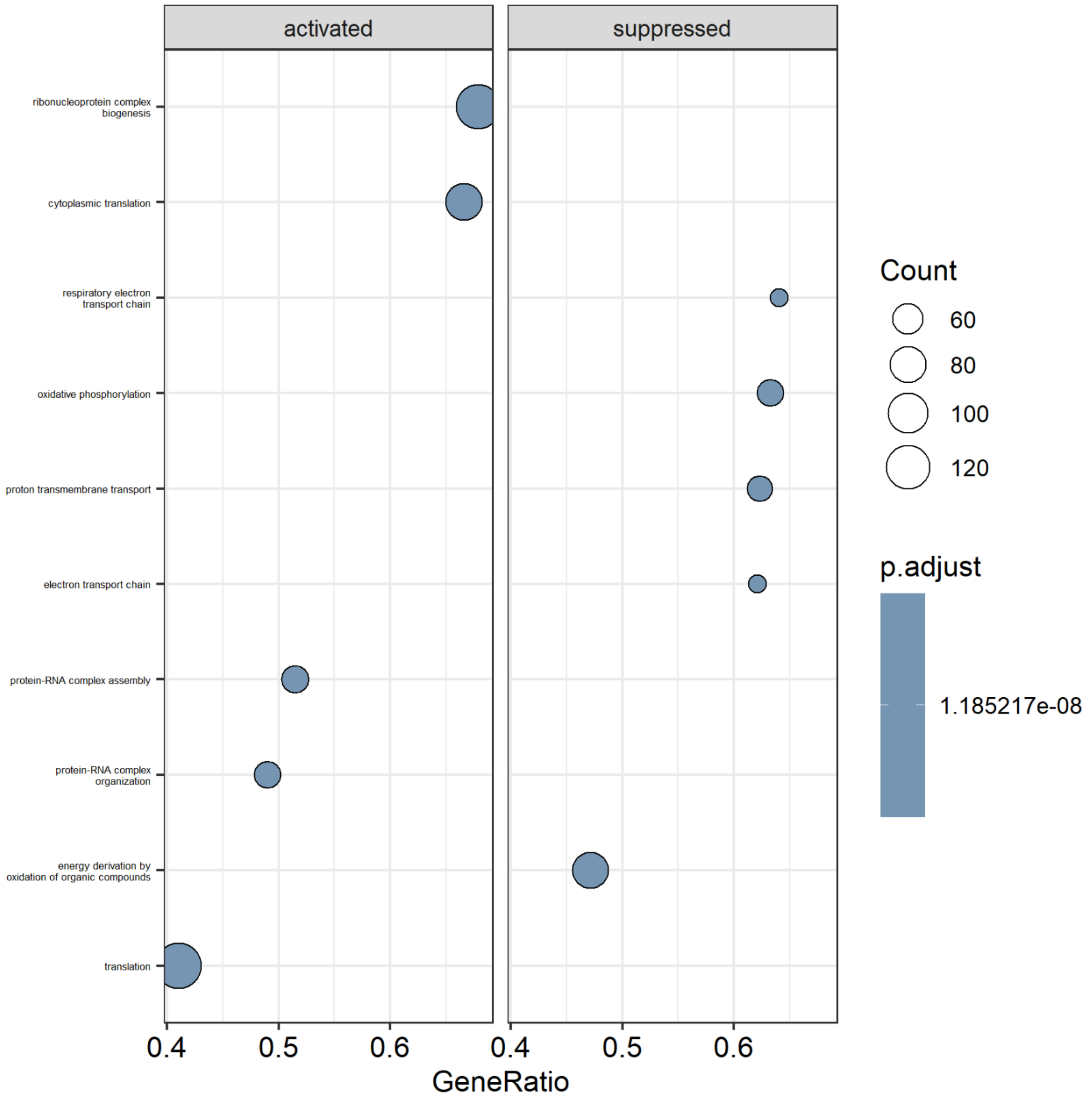

**S2 (continued).** (Z) 4AG vs. 4C dotplot showing activated and suppressed processes, including gene count and p-value (adjusted according to Benjamini-Hochberg) per process, based on Gene Ontology Biological Process. Based on global proteomics.

### Gene Ontology – Cellular Component

#### *Barplot*

**AA**

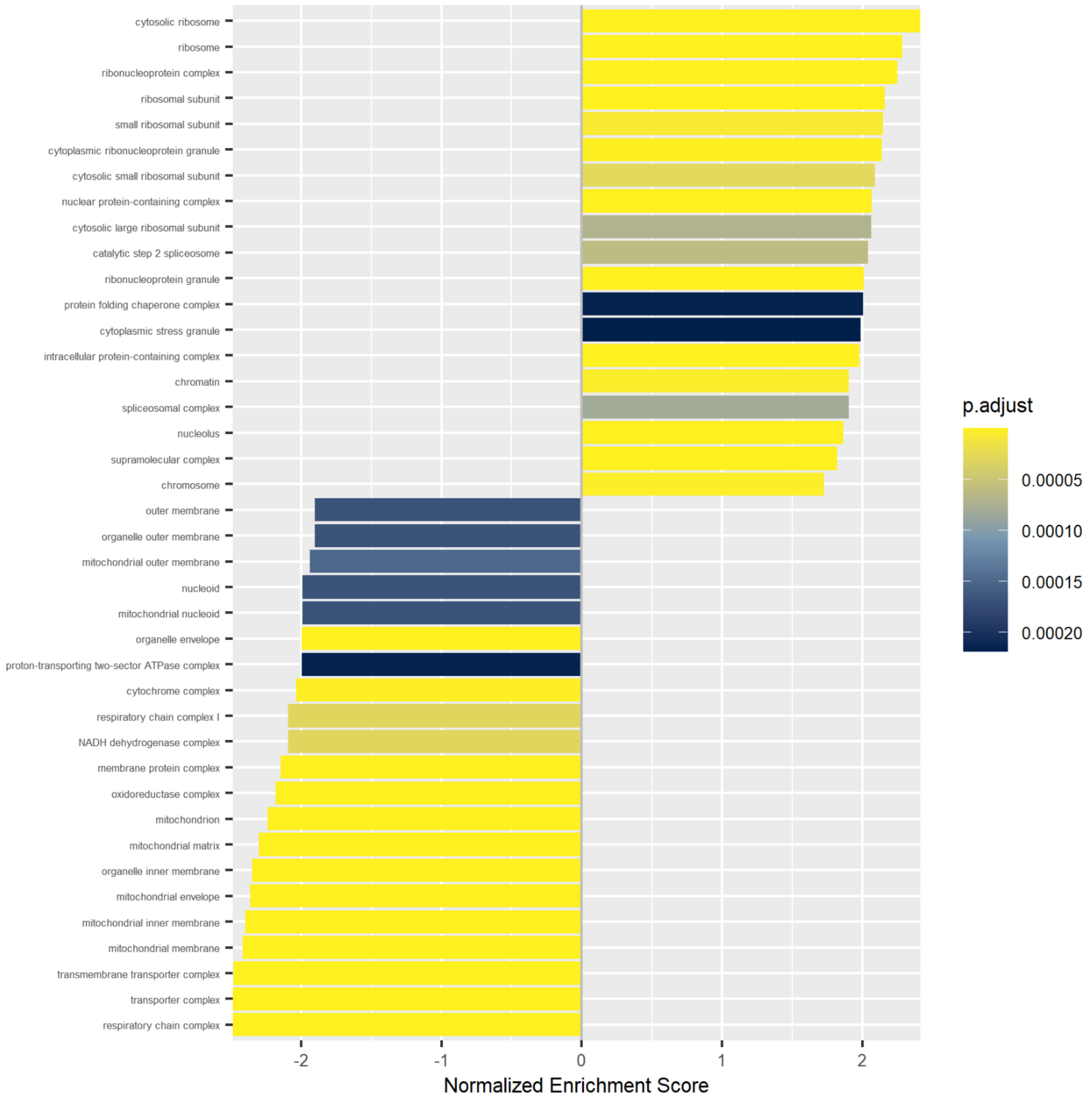

**S2 (continued).** (AA) 4AG vs. 4C barplot showing normalized enrichment score and p-value (adjusted according to Benjamini-Hochberg) for enriched processes based on Gene Ontology Cellular Component. Based on global proteomics.

*Dotplot*

**AB**

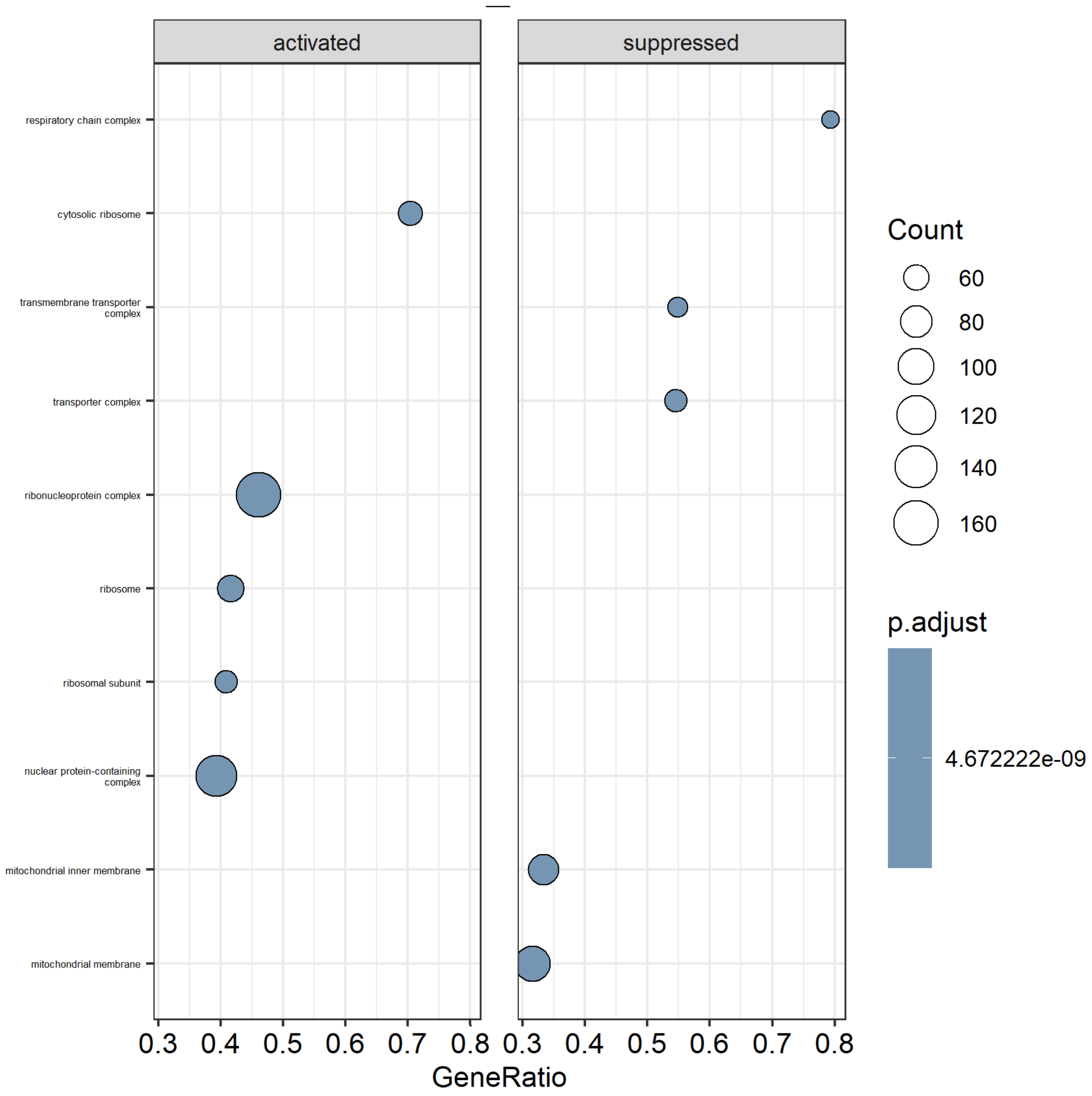

**S2 (continued).** (AB) 4AG vs. 4C dotplot showing activated and suppressed processes, including gene count and p-value (adjusted according to Benjamini-Hochberg) per process, based on Gene Ontology Cellular Component. Based on global proteomics.

### Gene Ontology – Molecular Function

*Barplot*

**AC**

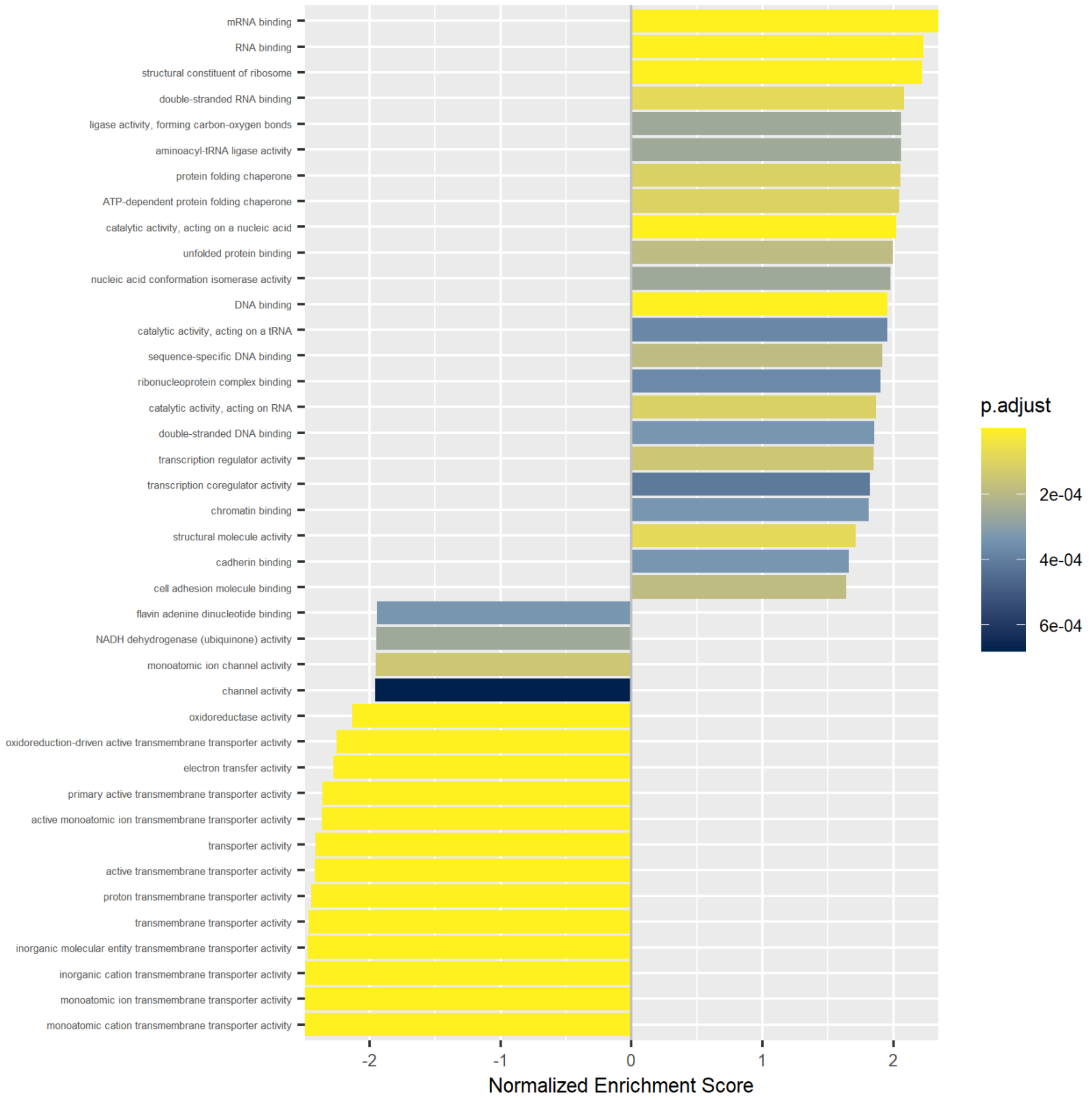

**S2 (continued).** (AC) 4AG vs. 4C barplot showing normalized enrichment score and p-value (adjusted according to Benjamini-Hochberg) for enriched processes based on Gene Ontology Molecular Function. Based on global proteomics.

*Dotplot*

**AD**

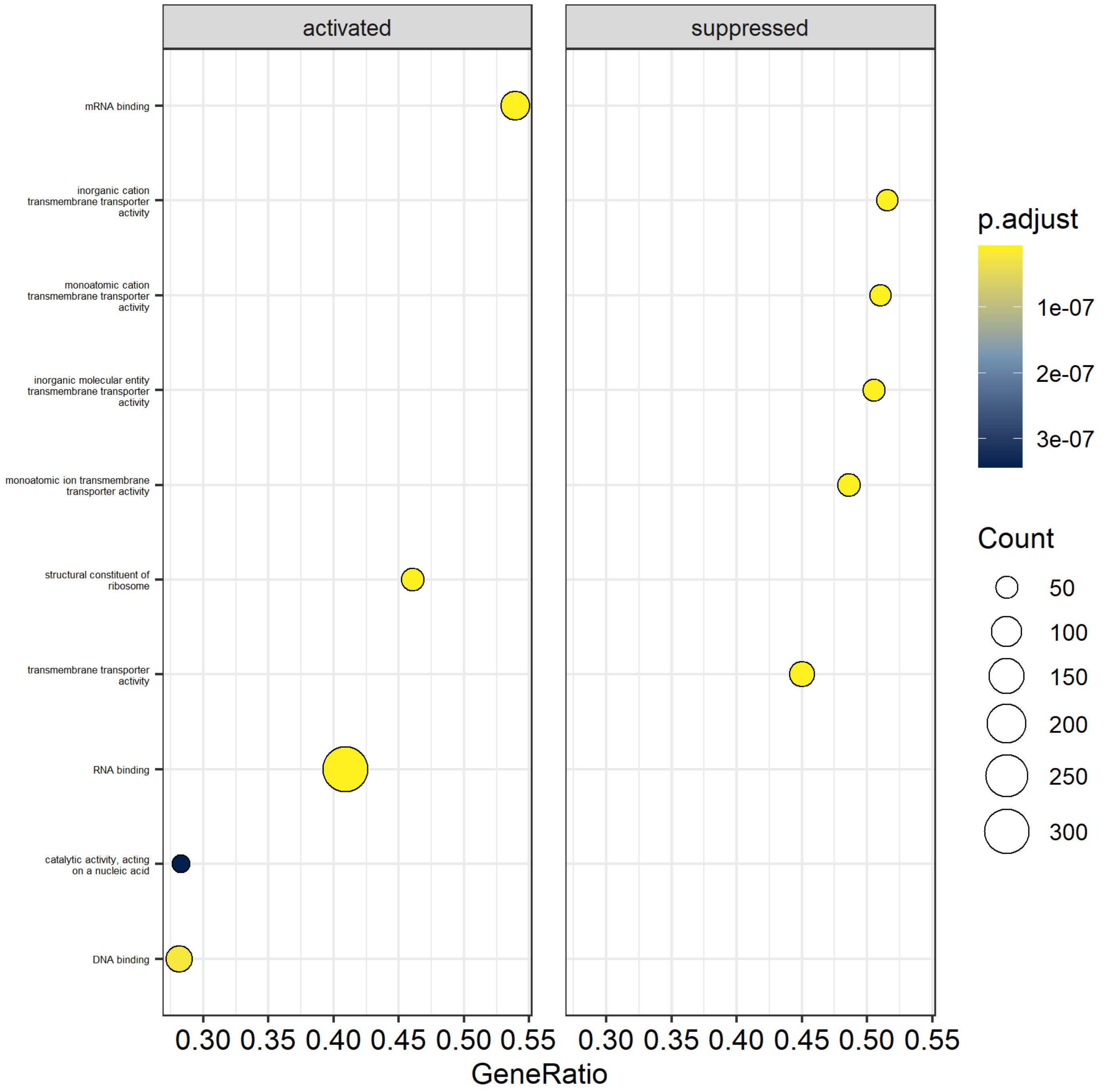

**S2 (continued).** (AD) 4AG vs. 4C dotplot showing activated and suppressed processes, including gene count and p-value (adjusted according to Benjamini-Hochberg) per process, based on Gene Ontology Molecular Function. Based on global proteomics.

### Reactome – Pathway Analysis

*Barplot*

**AE**

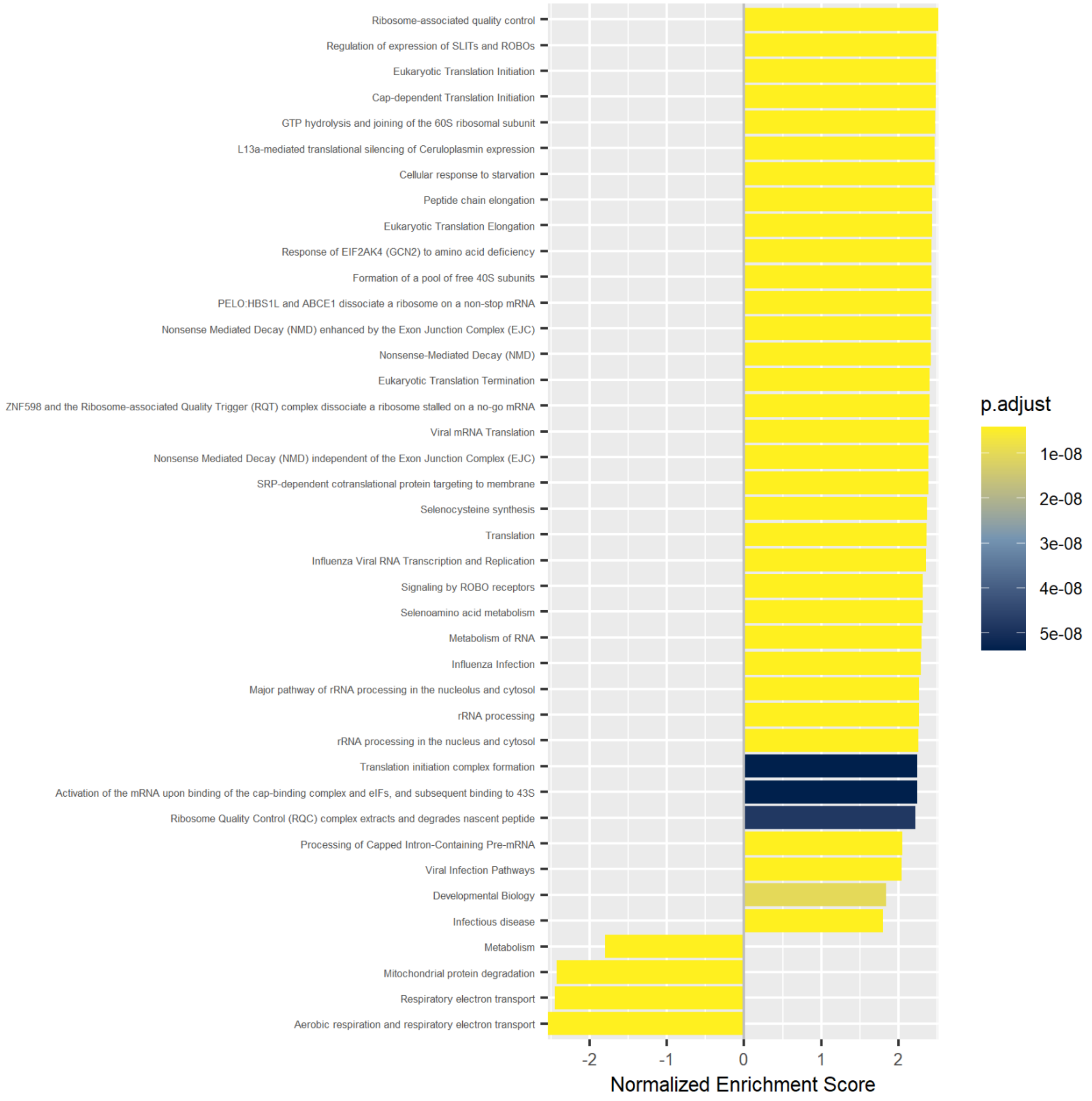

**S2 (continued).** (AE) 4AG vs. 4C barplot showing normalized enrichment score and p-value (adjusted according to Benjamini-Hochberg) for enriched processes, based on Reactome Pathway Analysis. Based on global proteomics.

*Dotplot*

**AF**

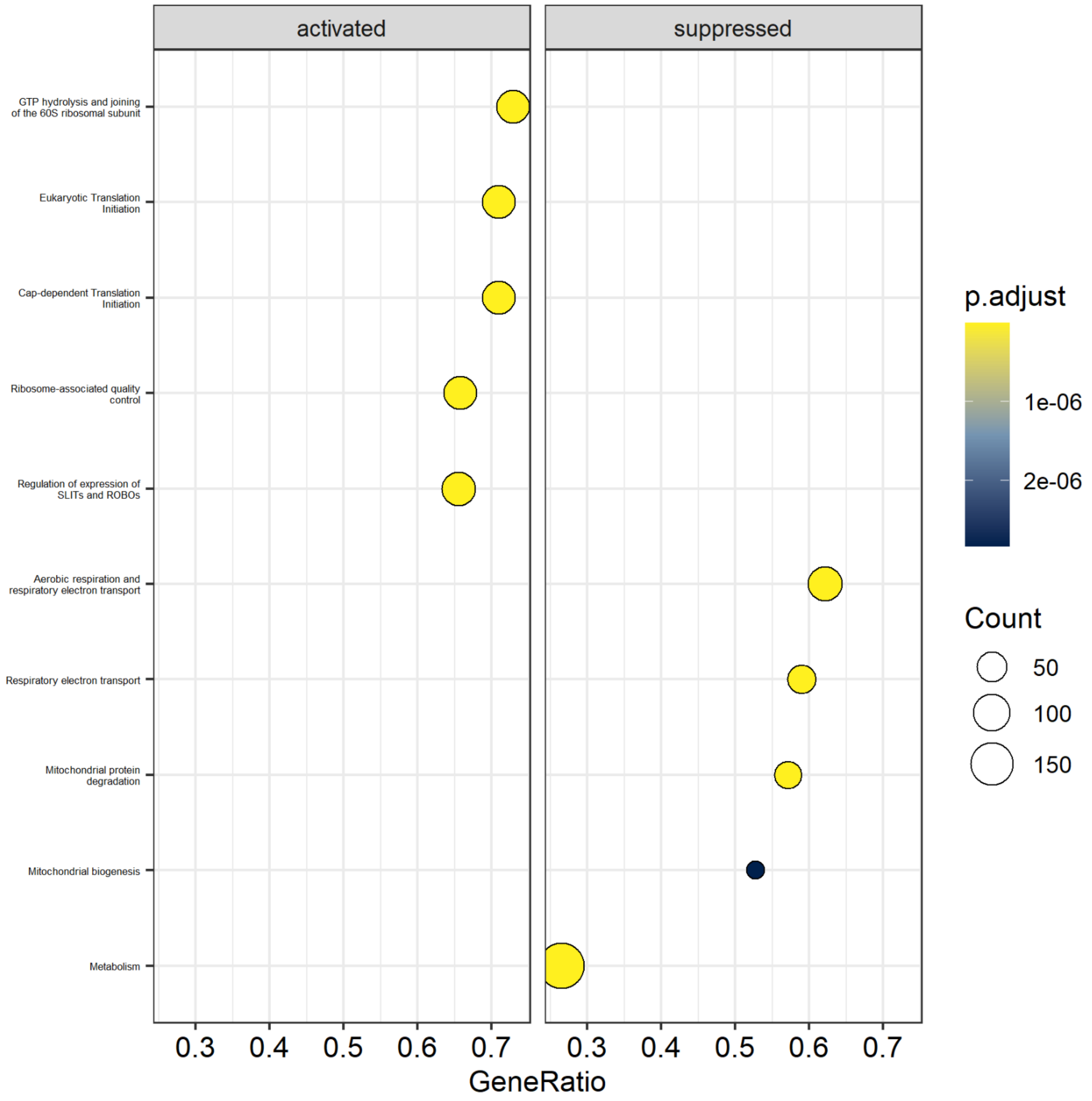

**S2 (continued).** (AF) 4AG vs. 4C dotplot showing activated and suppressed processes, including gene count and p-value (adjusted according to Benjamini-Hochberg) per process, based on Reactome Pathway Analysis. Based on global proteomics.

# 4BG

### Gene Ontology - Biological Process

#### *Barplot*

**AG**

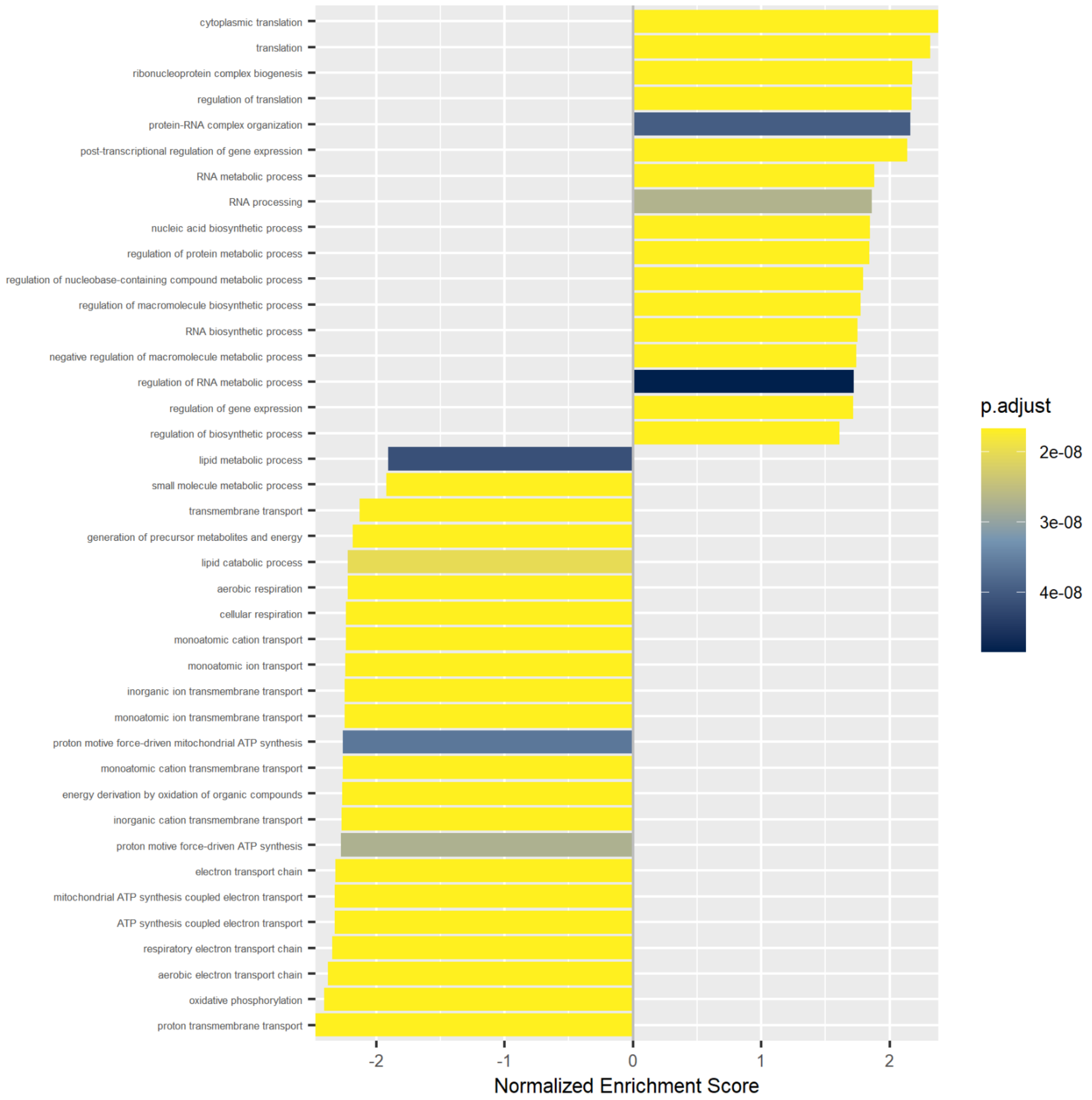

**S2 (continued).** (AG) 4BG vs. 4C barplot showing normalized enrichment score and p-value (adjusted according to Benjamini-Hochberg) for enriched processes based on Gene Ontology Biological Process. Based on global proteomics.

*Dotplot*

**AH**

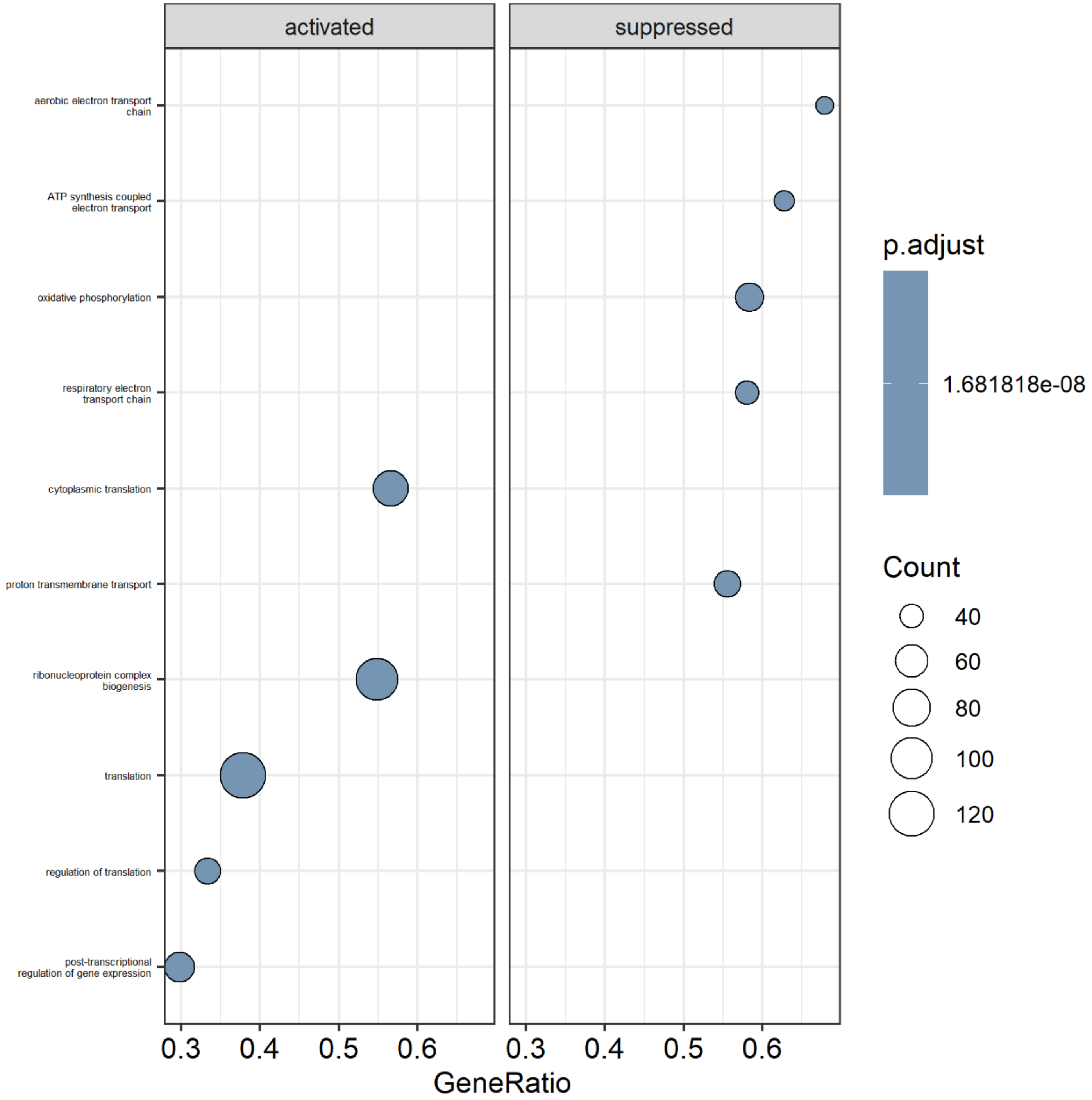

**S2 (continued).** (AH) 4BG vs. 4C dotplot showing activated and suppressed processes, including gene count and p-value (adjusted according to Benjamini-Hochberg) per process, based on Gene Ontology Biological Process. Based on global proteomics.

### Gene Ontology – Cellular Component

*Barplot*

**AI**

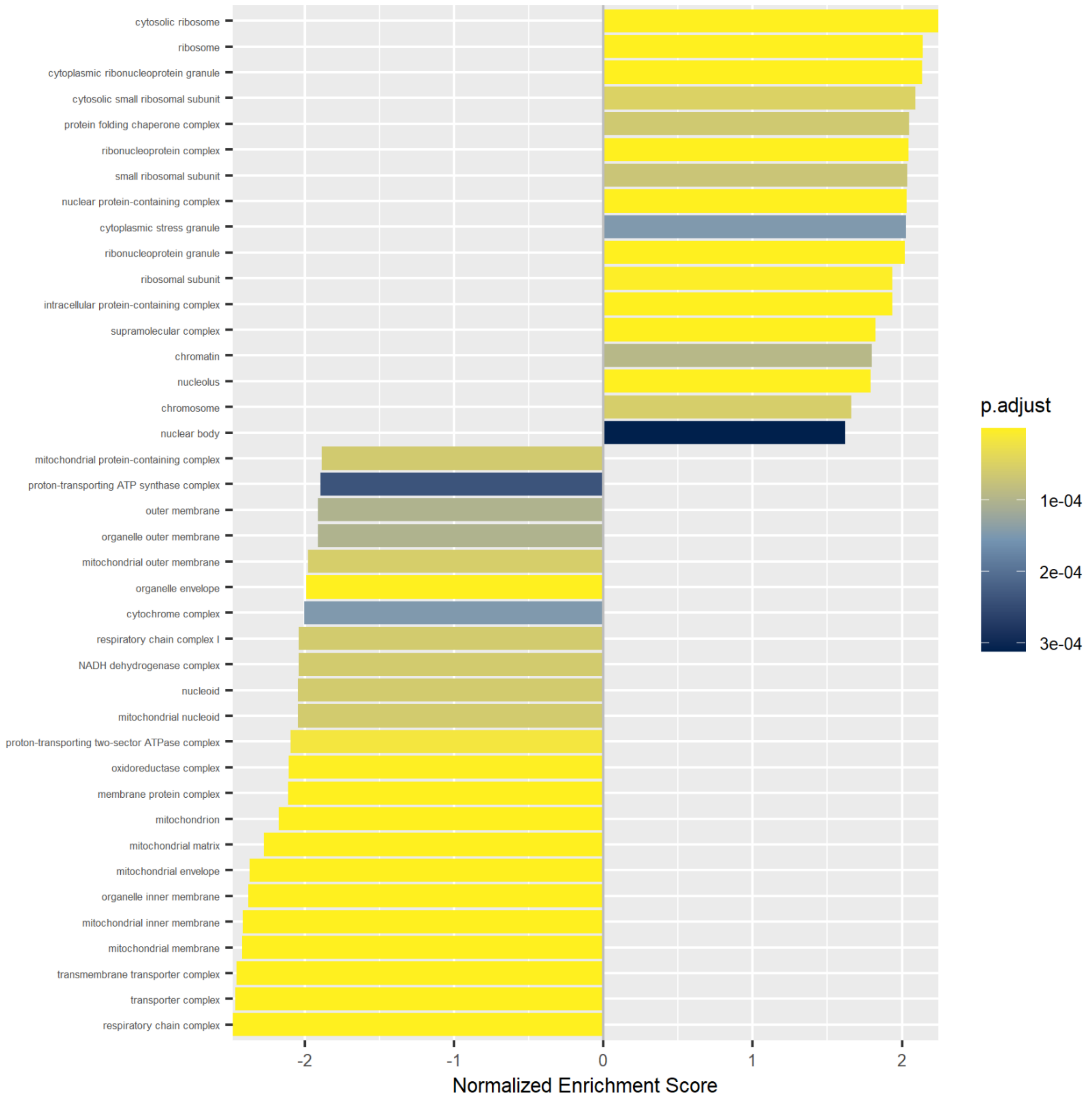

**S2 (continued).** (AI) 4BG vs. 4C barplot showing normalized enrichment score and p-value (adjusted according to Benjamini-Hochberg) for enriched processes based on Gene Ontology Cellular Component. Based on global proteomics.

*Dotplot*

**AJ**

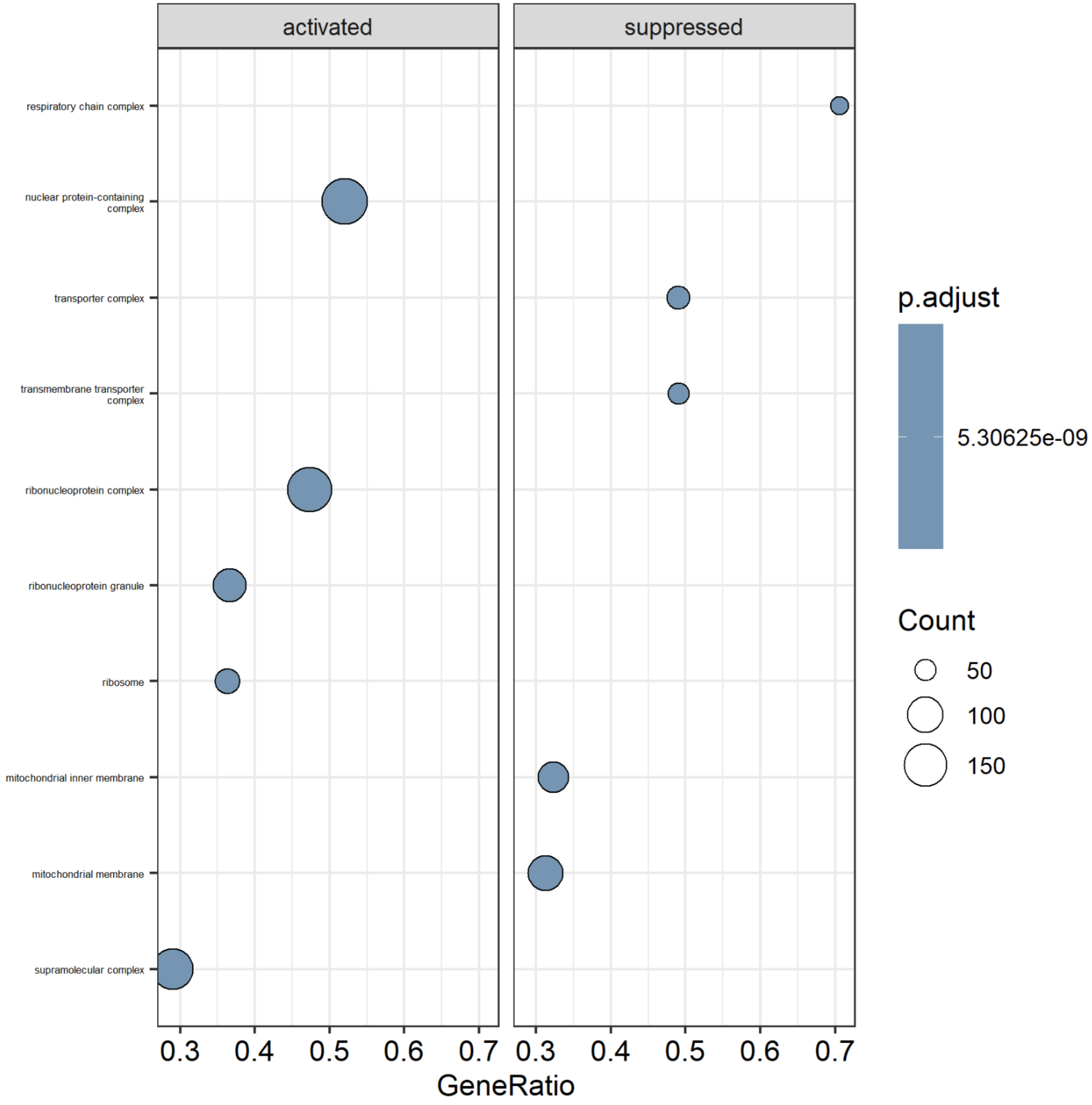

**S2 (continued).** (AJ) 4BG vs. 4C dotplot showing activated and suppressed processes, including gene count and p-value (adjusted according to Benjamini-Hochberg) per process, based on Gene Ontology Cellular Component. Based on global proteomics.

### Gene Ontology – Molecular Function

*Barplot*

**AK**

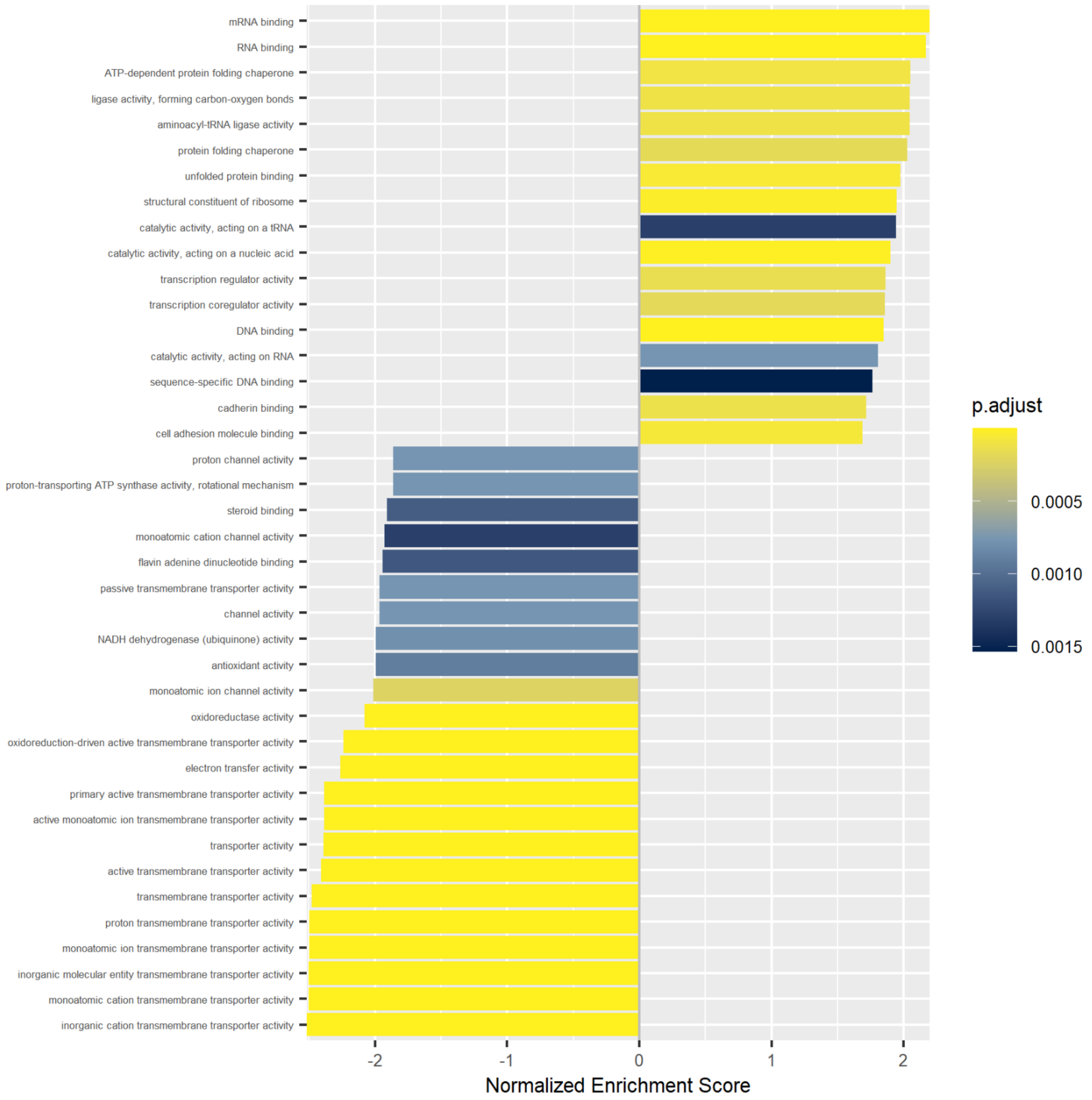

**S2 (continued).** (AK) 4BG vs. 4C barplot showing normalized enrichment score and p-value (adjusted according to Benjamini-Hochberg) for enriched processes based on Gene Ontology Molecular Function. Based on global proteomics.

*Dotplot*

**AL**

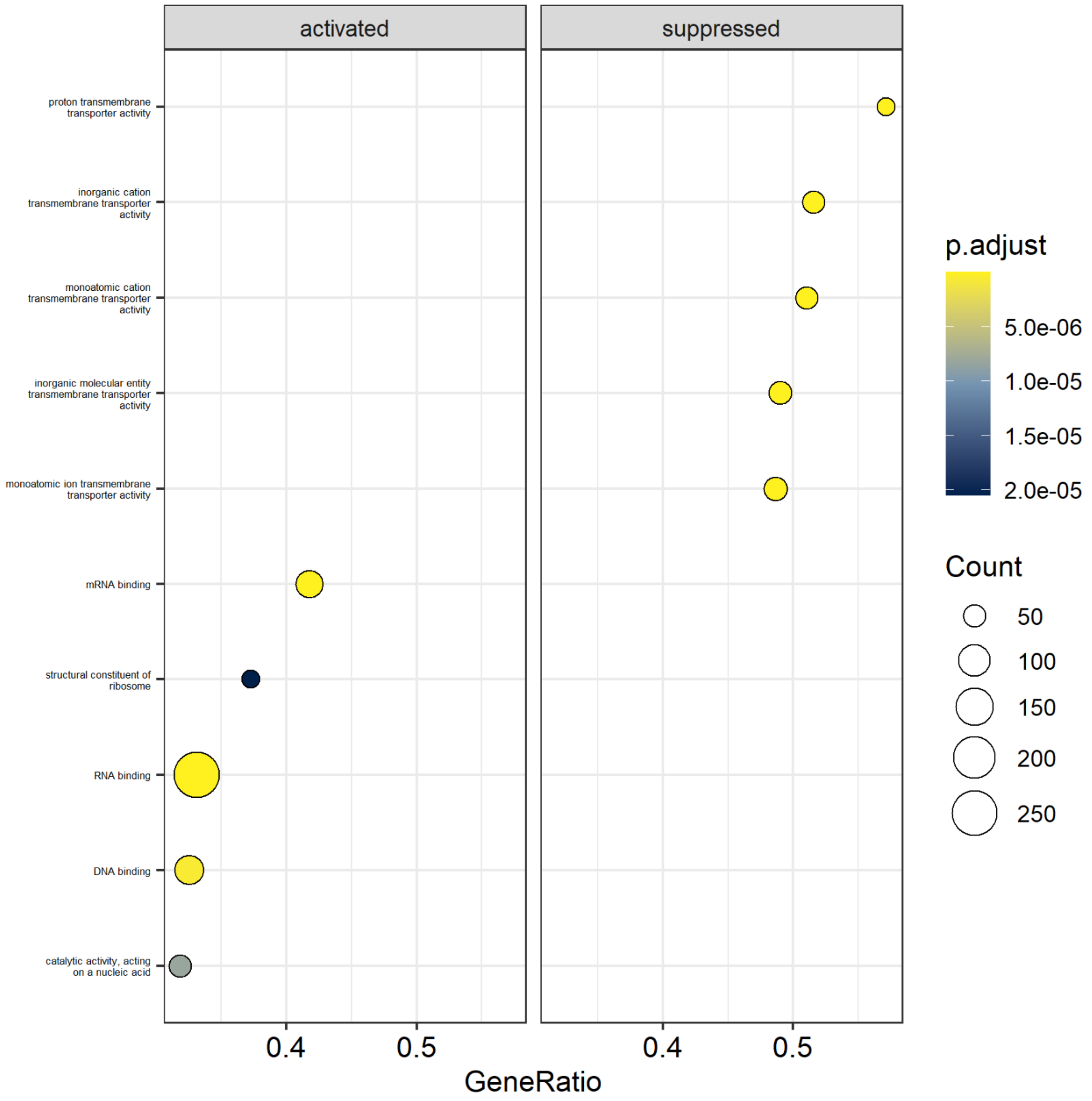

**S2 (continued).** (AL) 4BG vs. 4C dotplot showing activated and suppressed processes, including gene count and p-value (adjusted according to Benjamini-Hochberg) per process, based on Gene Ontology Molecular Function. Based on global proteomics.

### Reactome – Pathway Analysis

#### *Barplot*

### AM

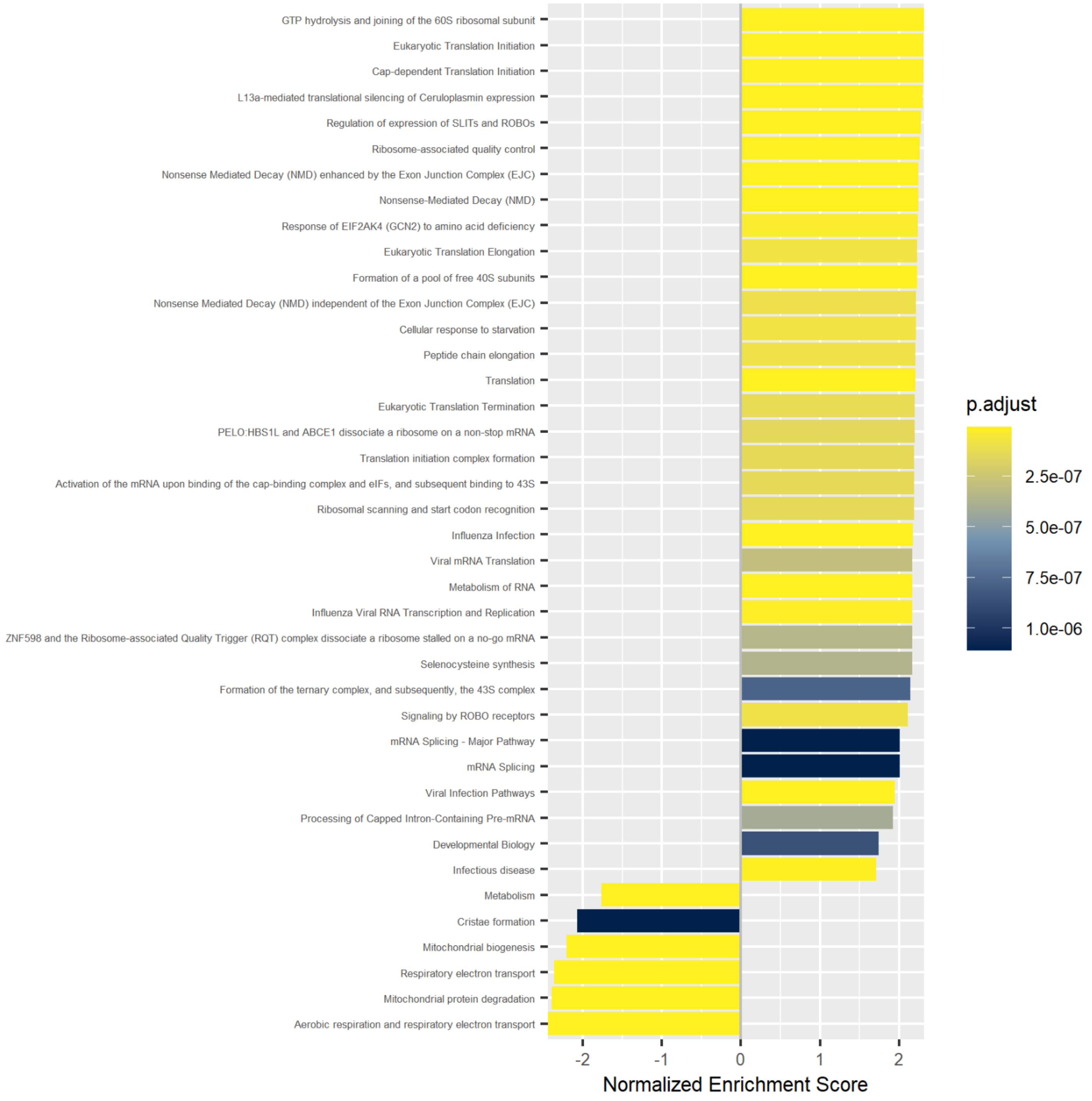

**S2 (continued).** (AM) 4BG vs. 4C barplot showing normalized enrichment score and p-value (adjusted according to Benjamini-Hochberg) for enriched processes based on Reactome Pathway Analysis. Based on global proteomics.

*Dotplot*

**AN**

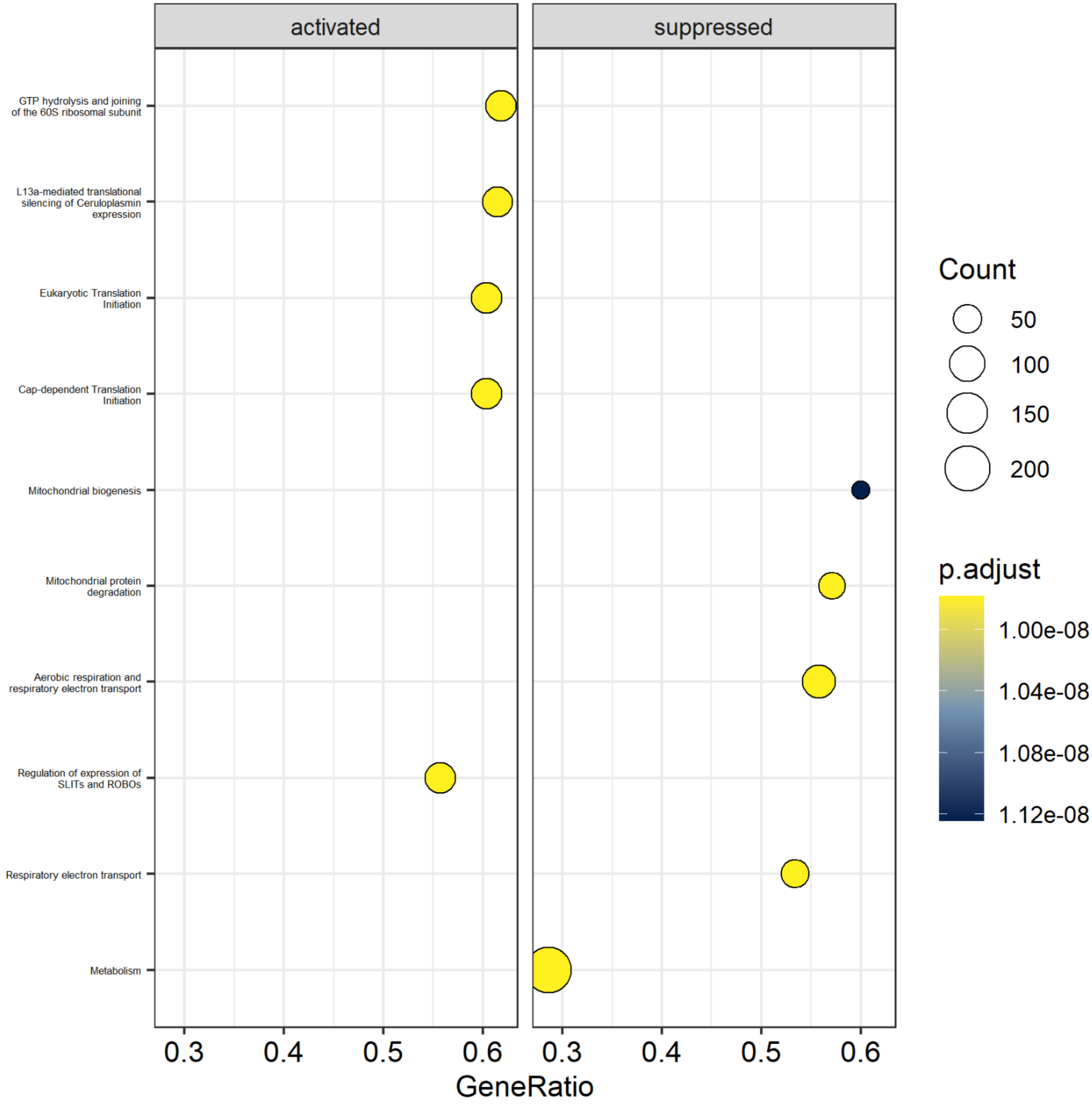

**S2 (continued).** (AN) 4BG vs. 4C dotplot showing activated and suppressed processes, including gene count and p-value (adjusted according to Benjamini-Hochberg) per process, based on Reactome Pathway Analysis. Based on global proteomics.

### References

**AO**

**S2 (continued).** (AO) The in-house R processing pipeline was based on the following publications:

[1] J.J. Allaire, C. Dervieux, quarto: R Interface to Quarto Markdown Publishing System, 2025. https://github.com/quarto-dev/quarto-r.

[2] J.M. Barbone, J.M. Garbuszus, openxlsx2: Read, Write and Edit xlsx Files, 2026. https://janmarvin.github.io/openxlsx2/.

[3] H. Bengtsson, matrixStats: Functions that Apply to Rows and Columns of Matrices (and to Vectors), 2025. https://github.com/HenrikBengtsson/matrixStats.

[4] B. Bolstad, preprocessCore: A collection of pre-processing functions, 2026. https://github.com/bmbolstad/preprocessCore.

[5] M. Carlson, org.Hs.eg.db: Genome wide annotation for Human, 2026.
